# Cell-autonomous and non-cell-autonomous effects of arginase-II on cardiac aging

**DOI:** 10.1101/2024.01.17.576017

**Authors:** Duilio M. Potenza, Xin Cheng, Guillaume Ajalbert, Andrea Brenna, Marie-Noelle Giraud, Aurelien Frobert, Stephane Cook, Kirsten D. Mertz, Zhihong Yang, Xiu-Fen Ming

**Author notes:** Corresponding authors: Duilio M. Potenza, Xiu-Fen Ming, Zhihong Yang. **Statements relating to relevant ethics:** Experimental work with animals was approved by the Ethical Committee of the Veterinary Office of Fribourg Switzerland (2020-01-FR, and 2019-21-FR) and performed in compliance with guidelines on animal experimentation at our institution. The study was conducted according to the principles expressed in the Declaration of Helsinki. Experiments with human heart biopsies were approved by the Ethics Committee of Northwestern and Central Switzerland (Project-ID 2021-01445).

## Abstract

Aging is a predominant risk factor for heart disease. Aging heart reveals low-grade chronic inflammation, cell apoptosis, cardiac fibrosis, and increased vulnerability to ischemic injury. The underlying molecular mechanisms responsible for the cardiac aging phenotype and its susceptibility to injury are far from being fully understood. Although previous literature reports a role of the mitochondrial enzyme arginase-II (Arg-II) in development of heart failure, contradictory results are reported and no systematic analysis of cellular expression and localization of Arg-II in the heart has been performed. Whether and how Arg-II participates in cardiac aging are still unknown. In this study, we demonstrate, to our surprise, that Arg-II is not expressed in cardiomyocytes from aged mice and human patients, but upregulated in non-myocytes of the aging heart, including macrophages, fibroblasts, endothelial cells. Mice with genetic deficiency of *arg-ii* (*arg-ii^−/−^*) are protected from age-associated cardiac inflammation, myocyte apoptosis, interstitial and perivascular fibrosis, endothelial-mesenchymal transition (EndMT), and susceptibility to ischemic injury. Further experiments show that Arg-II mediates IL-1β release from macrophages of old mice, contributing to the above-described cardiac aging phenotype. In addition, Arg-II enhances mitochondrial reactive oxygen species (mtROS) and activates cardiac fibroblasts that is inhibited by inhibition of mtROS. Thus, our study demonstrates a non-cell-autonomous effect of Arg-II on cardiomyocytes, fibroblasts, and endothelial cells mediated by IL-1β from aging macrophages as well as a cell-autonomous effect of Arg-II through mtROS in fibroblasts contributing to cardiac aging phenotype.

## Introduction

Aging is a predominant risk factor for cardiovascular disease (CVD) (***Moturi et al., 2022***). Cardiac aging is accompanied by low-grade chronic inflammation termed “inflammaging” (***Cevenini et al., 2013***), which is linked to gradual development of cardiac fibrosis and heart failure, whereby macrophages, cardiac fibroblasts, and endothelial cells play critical roles in age-associated cardiac remodeling and dysfunction (***Abdellatif et al., 2023; Meschiari et al., 2017***). Studies in experimental animal models and in humans provide evidence demonstrating that aging heart is more vulnerable to stressors such as ischemia/reperfusion injury and myocardial infarction and exhibits less efficient reparative capability after injury as compared to the heart of young individuals (***Bujak et al., 2008; Mariani et al., 2000***). Even in the heart of apparently healthy individuals of old age, chronic inflammation, cardiomyocyte senescence, cell apoptosis, interstitial/perivascular tissue fibrosis, endothelial dysfunction and endothelial-mesenchymal transition (EndMT), and cardiac dysfunction either with preserved or reduced ejection fraction rate are observed (***Abdellatif et al., 2023; Ruiz-Meana et al., 2020***). Despite the intensive investigation in the past, the underlying causative cellular and molecular mechanisms responsible for the cardiac aging phenotype and the susceptibility of aging heart to injurious stressors are not fully elucidated, yet. Moreover, most preclinical studies preferentially use relatively juvenile animal models which hardly recapitulate clinical scenario. It is therefore critical to investigate mechanisms of cardiac aging phenotype with a naturally advanced aging animal model.

Research in the past years suggests a wide range of functions of the enzyme arginase, including the cytosolic isoenzyme arginase-I (Arg-I) and the mitochondrial isoenzyme arginase-II (Arg-II) in organismal aging as well as age-related organ structural remodeling and functional changes (***Caldwell et al., 2015; Li et al., 2022***). These functions of arginase have been implicated in cardiac and vascular aging (***Xiong, Yepuri, Montani, et al., 2017; Yepuri et al., 2012***) and cardiac ischemia/reperfusion injury in rodents (***Jung et al., 2010***). Arg-I, originally found in hepatocytes, plays a crucial role in urea cycle in the liver to remove ammonia from the blood (***Sin et al., 2015***), whereas Arg-II is inducible in many cell types under pathological conditions, contributing to chronic tissue inflammation and remodeling (***Moretto et al., 2019***). Both enzymes are able to catalyze the divalent cation-dependent hydrolysis of L-arginine (***Ming et al., 2013***). Studies in the literature including our own, demonstrate that Arg-II is involved in aging process, and promotes organ inflammation and fibrosis and inhibition of arginase or genetic ablation of *arg-ii* slows down aging process and protects against age-associated organ degeneration such as lung (***Zhu et al., 2023***), pancreas (***Xiong, Yepuri, Necetin, et al., 2017***) and heart (***Xiong, Yepuri, Montani, et al., 2017***). However, controversial results on expression of arginase and its isoforms at the cellular levels in the heart are reported. While some studies linked Arg-I to the cardiac ischemia/reperfusion injury (***Jung et al., 2010***), others suggest a role of Arg-II in this context (***Heusch et al., 2010***). Even protective functions of Arg-II are suggested in some studies, based on correlation of Arg-II levels with cardiac injury or systemic administration of Arg-II in rat models (***Huang et al., 2018; Lu et al., 2012***). Importantly, cellular localization of arginase and its isoenzyme in the heart, i.e., expression of arginase in cardiomyocytes and non-cardiomyocytes have not been systematically analyzed and confirmed. Finally, whether and how Arg-II participates in cardiac aging and whether it is through cell-autonomous and/or paracrine effects are not known.

Therefore, our current study is aimed to investigate which cell types in the heart express arginase and what enzymatic isoforms. How arginase and this isoenzyme in these cells influence the cardiac aging process and enhance susceptibility of the aging heart to ischemia/reperfusion injury.

## Results

### 1. Increased *arg-ii* expression in aging heart

In heart from young (3-4 months in age) and old (22-24 months in age) mice, an age-associated increase in *arg-ii* expression in both *wt* male and female mice are observed (**Fig. 1A**). This age-associated increase in *arg-ii* expression is more pronounced in females than in males (**Fig. 1A**). In contrast to *arg-ii*, *arg-i* mRNA expression is very low in the heart (with very high Ct values, data not shown), and no age-dependent changes of *arg-i* gene expression are observed in *wt* and in *arg-ii^−/−^* mice of either gender (**Suppl. Fig. 1A** and **Suppl. Fig 1B**), suggesting a specific enhancement of Arg-II in aging heart. Further experiments are thus focused on Arg-II and female mice. We made effort to analyze Arg-II protein levels by immunoblotting in the heart tissues. However, immunoblotting is not sensitive enough to detect Arg-II protein in the heart tissue of either young or old animals. Therefore, further experiments are performed to detect Arg-II proteins at the cellular level with confocal microscopic immunofluorescence staining as described in later sections.

**Fig. 1.**
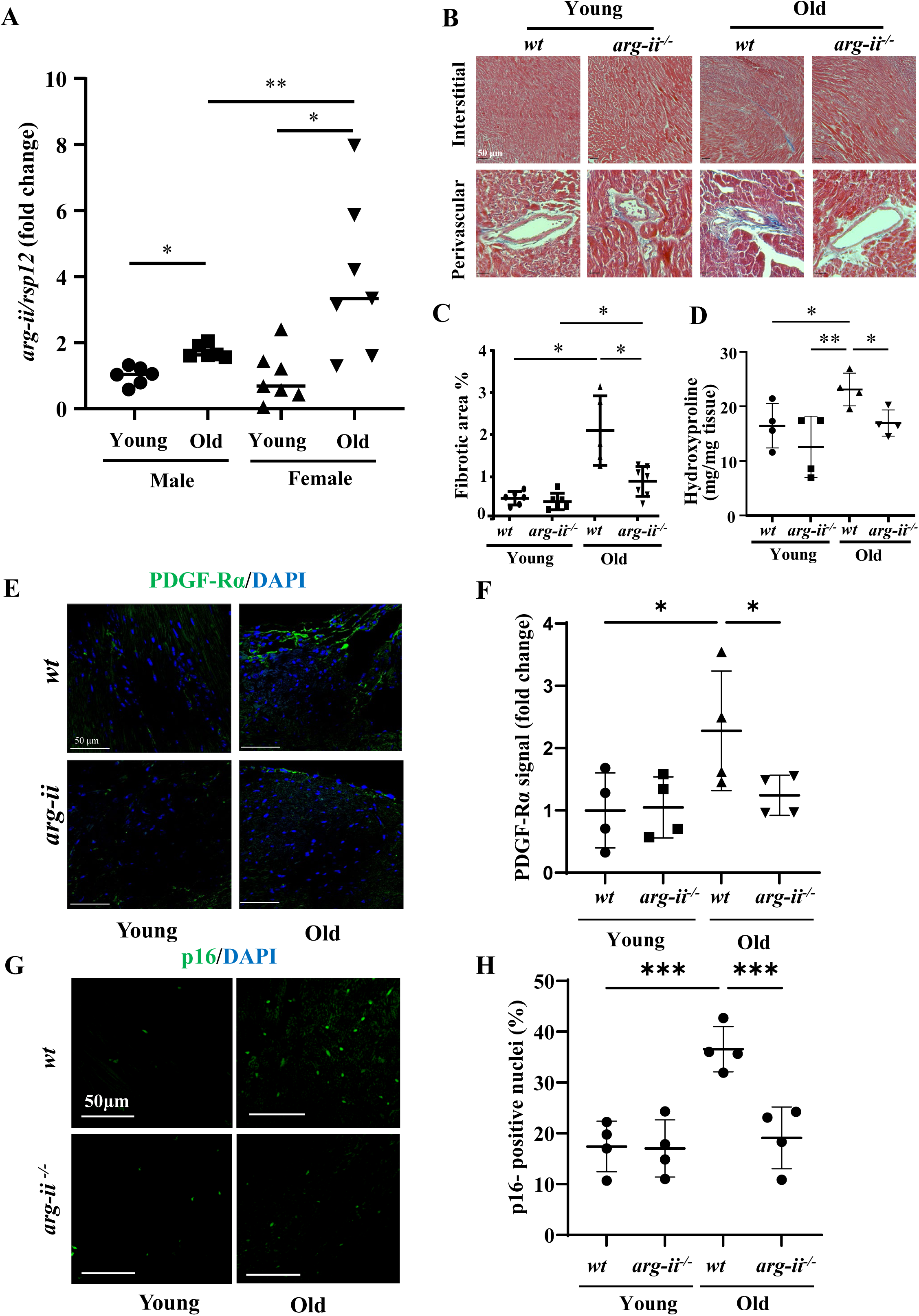
Age-associated increase in Arg-II levels in mouse heart. (**A**) *arg-ii* mRNA levels of male and female young (3-4 months) and old (20-22 months) wild type (*wt*) heart tissues analyzed by qRT-PCR. *rps12* served as the reference (n=6 to 7 animals per group); (**B**) Representative histological images of heart interstitial and perivascular fibrosis in young and old *wt* and *arg-ii^−/−^* female mice. Fibrosis is shown by the blue-colored Trichrome Masson’s staining. Scale bar = 50 µm. (n=5 to 7 mice per group); (**C**) Quantification of total fibrotic area in cardiac tissue (% of total area); (**D**) Hydroxyproline content of mouse heart from young and old *wt* and *arg-ii^−/−^*female mice. (n=4 mice in each group); (**E**) Representative confocal images showing immunofluorescence staining of PDGF-Rα (green, fibroblasts marker) in young and old *wt* and *arg-ii^−/−^* heart tissue. DAPI (blue) is used to stain nuclei. Scale bar = 50 µm; (**F**) Relative PDGF-Rα signal quantification of confocal images (n= 4 per each group). (**G**) Representative confocal images showing immunofluorescence staining of p16 (green, senescent marker) in young and old *wt* and *arg-ii^−/−^* heart tissue. DAPI (blue) is used to stain nuclei. Scale bar = 50 µm; (**H**) Percentage of p16^+^ nuclei in the four groups (n= 4). The values shown are mean ± SD. Data are presented as the fold change to the young-*wt* group, except for panel D. *P ≤ 0.05, **P ≤ 0.01, ***P ≤ 0.005 between the indicated groups. *wt*, wild-type mice; *arg-ii^−/−^*, *arg-ii* gene knockout mice.

### 2. *Arg-ii* ablation reduces cardiac aging phenotype

Next we investigated the impact of *arg-ii* ablation on cardiac phenotype. Masson’s staining reveals an enhanced peri-vascular and interstitial fibrosis in *wt* old mice, which is mitigated in age-matched *arg-ii^−/−^* animals (**Fig. 1B, 1C**). The age-associated increase in cardiac fibrosis and its inhibition by *arg-ii^−/−^* are further confirmed by quantitative measurements of collagen content as assayed by determination of hydroxyproline levels in the heart tissues (**Fig. 1D**). The increased cardiac fibrosis in aging is however, associated with decreased mRNA levels of *collagen-Iα* (*col-Iα*) and *collagen-IIIα* (*col-IIIα*), the major isoforms of pre-collagen in the heart (**Suppl. Fig. 2A and 2B**), which is a well-known phenomenon in cardiac fibrotic remodeling (***Besse et al., 1994; Horn et al., 2016***). The results demonstrate that age-associated cardiac fibrosis and prevention in *arg-ii^−/−^* mice is due to alterations of translational and/or post-translational regulations including collagen synthesis and/or degradation. Furthermore, an increased density of fibroblasts as characterized by positive staining of PDGF-Rα is observed in *wt* old mice, but not in age-matched *arg-ii^−/−^* animals (**Fig. 1E** and **Fig. 1F**). Interestingly, an age-associated increase in p16^ink4^ positive senescent cells in *wt* mouse heart is prevented in the *arg-ii^−/−^* animals (**Fig. 1G and 1H**).

**Fig. 2.**
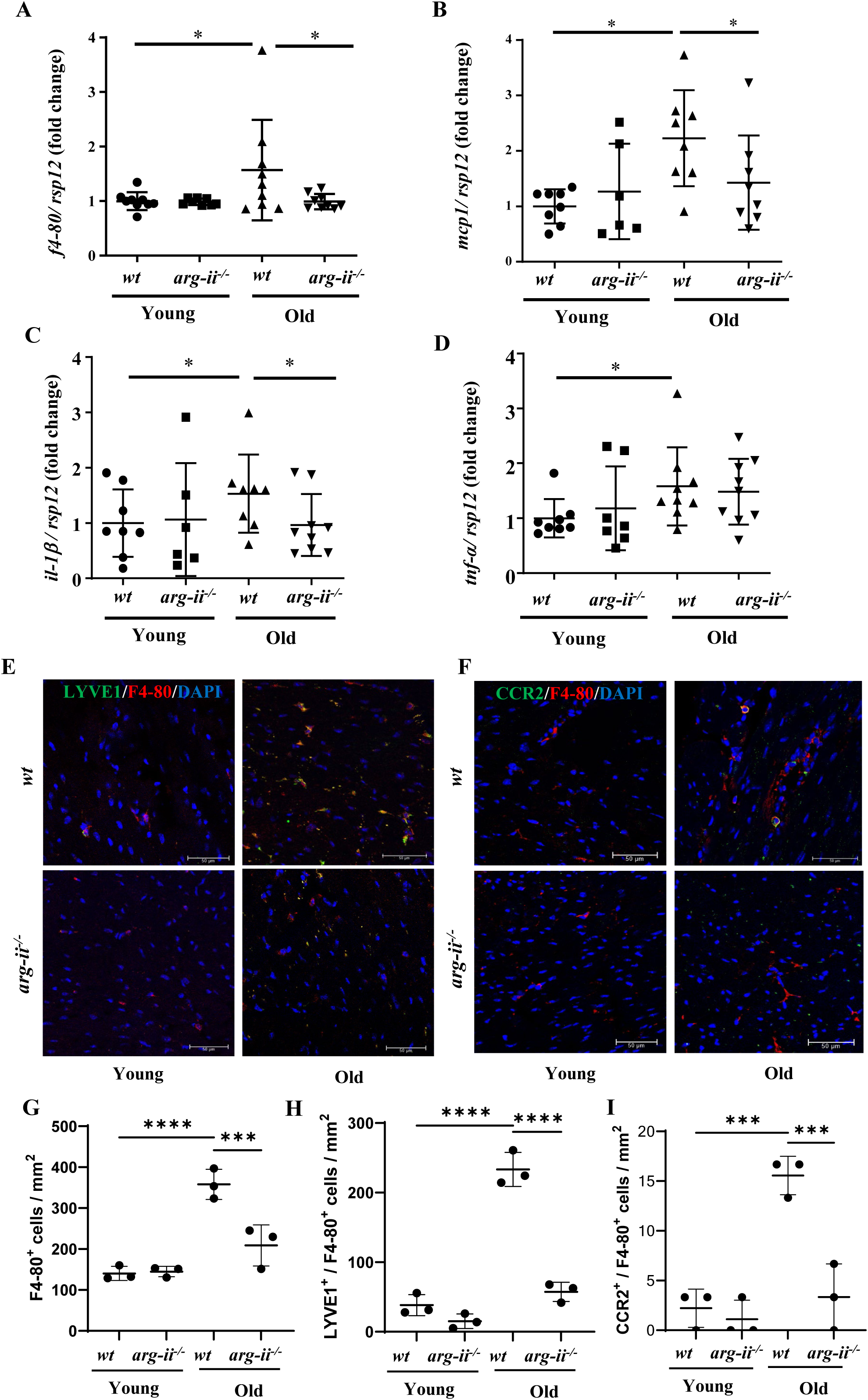
Age-associated elevation in inflammatory cytokines and apoptosis in heart is prevented in female *arg-ii^−/−^* mice. mRNA expression levels of (**A**) *f4/80*, (**B**) *mcp-1*, (**C**) *il-1β* and (**D**) *tnf-α* in young and old *wt* and *arg-ii^−/−^* female mouse hearts were analyzed by qRT-PCR. *rps12* served as the reference (n=6-8 mice per group); (**E**) Representative confocal images of young and old *wt* and *arg-ii^−/−^* heart tissue showing co-localization of LYVE1 (green) and F4-80 (red, mouse macrophage marker), (**F**) Representative co-localization images of CCR2 (green) and F4-80 (red) in *wt* and *arg-ii^−/−^* heart tissues. DAPI (blue) is used to stain nuclei. Scale bar = 50 µm. Graph showing the quantification of (**G**) F4-80^+^ cells, (**H**) LYVE1^+^/F4-80^+^ cells and (**I**) CCR2^+^/F4-80^+^ cells per mm^2^ in heart tissue of *wt* and *arg-ii^−/−^* young and old mice (n= 3 per each group). The values shown are mean ± SD. Data are expressed as fold change to the young *wt* group, expect for panels G to I. *p ≤ 0.05, ***p ≤ 0.005 and ****p ≤ 0.001 between the indicated groups. *wt*, wild-type mice; *arg-ii^−/−^*, *arg-ii* gene knockout mice.

Moreover, a significant increase in macrophage marker *f4/80* (**Fig. 2A**) and elevated gene expression of numerous pro-inflammatory cytokines such as *mcp1*, *il-1β* and *tnf-α* are observed in aging heart of *wt* mice (**Fig. 2B to Fig. 2D**). This age-associated increase in the inflammatory markers and cytokine gene expression (except *tnf-α*) are overall significantly reduced or prevented in *arg-ii^−/−^* mice (**Fig. 2A** to **Fig. 2D**). In line with the gene expression of *f4/80*, immunofluorescence staining reveals an age-associated increase in the numbers of F4/80^+^ cells in the *wt* mouse heart, which is reduced in the age-matched *arg-ii^−/−^* animals (**Fig. 2E to 2G**), demonstrating that *arg-ii* gene ablation reduces macrophage accumulation in the aging heart. Interestingly, resident macrophages as characterized by LYVE1^+^/F4-80^+^ cells (**Fig. 2E** and **2H**) are predominant in the aging heart as compared to the infiltrated CCR2^+^/F4-80^+^ cells (**Fig. 2F and 2I**). The increase in both LYVE1^+^/F4-80^+^ and CCR2^+^/F4-80^+^ macrophages in aging heart is reduced in *arg-ii^−/−^* mice (**Fig. 2E, 2F, 2H, and 2I**).

We also observed an age-associated increase in total numbers of apoptotic cells as demonstrated by TUNEL staining in the heart of *wt* mice, which is reduced in age-matched *arg-ii^−/−^*animals (**Fig. 3A** and **Fig. 3B**). Further experiments with wheat germ agglutinin (WGA)-Alexa Fluor 488 (used to define cell boarders) (**Fig. 3C**) show a higher percentage of apoptotic cardiomyocytes (CM) than non-cardiomyocytes (NCM) in *wt* mice (**Fig. 3D**). This increased percentage of cardiomyocyte apoptosis is also prevented in age-matched *arg-ii^−/−^*animals (**Fig. 3D**). The percentage of apoptotic non-cardiomyocytes in aged *arg-ii^−/−^* mice tended to be reduced as compared to *wt* mice, but it does not reach statistical significance (**Fig. 3D**).

**Fig. 3.**
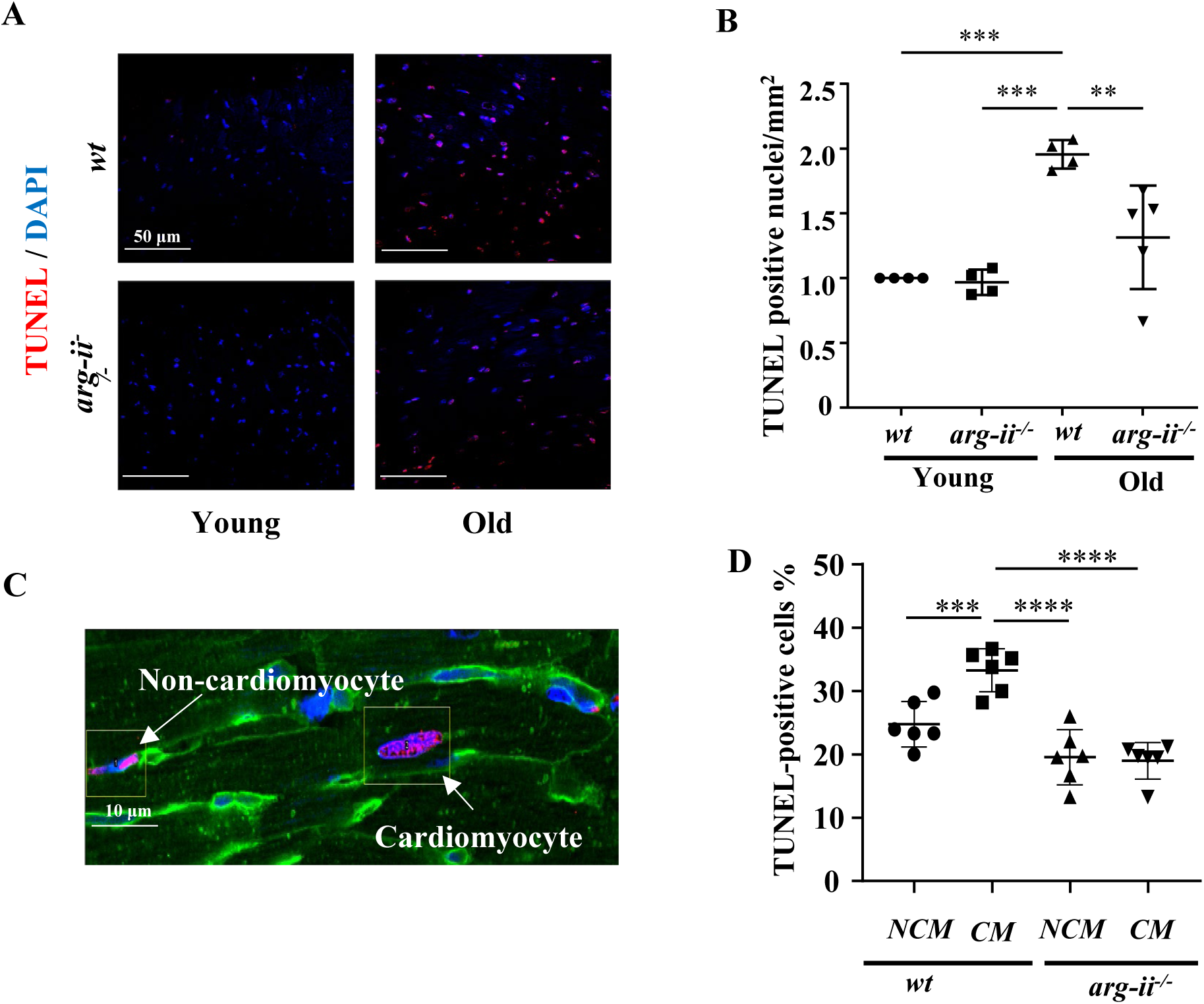
Age-associated elevation in apoptotic cardiomyocytes is prevented in female *arg-ii^−/−^* mice. **(A)** Representative confocal images and relative quantification of apoptotic cardiac cells in young and old *wt* and *arg-ii^−/−^* heart tissue. DAPI (blue) is used to stain nuclei. Scale bar = 50 µm; (**B**) graph showing the quantification of the TUNEL-positive cells in old *wt* and *arg-ii^−/−^* hearts; (**C**) Wheat germ agglutinin (WGA)-Alexa Fluor 488-conjugate was used to stain cell membrane, and separate cardiomyocytes and non-cardiomyocytes apoptotic cells. Cell distinction was based on cell size and shape. Scale bar = 10 µm; (**D**) Graphs showing the quantification of the TUNEL-positive cardiomyocytes (CM) and non-myocytes (NCM) in old *wt* and *arg-ii^−/−^* hearts. The values shown are mean ± SD. Data are expressed as fold change to the young *wt* group, expect for panel D. **p≤ 0.01, ***p ≤ 0.005 and ****p ≤ 0.001 between the indicated groups. *wt*, wild-type mice; *arg-ii^−/−^*, *arg-ii* gene knockout mice.

In addition, heart from old *wt* mice reveals elevated levels of SNAIL (the master regulator of End-MT process) and vimentin (mesenchymal marker), which is prevented in *arg-ii^−/−^*mice (**Fig. 4A** to **Fig. 4C**). No changes in CD31 (endothelial marker) levels are observed among the four groups (**Fig. 4A** and **Fig. 4D**). Immunofluorescence staining reveals that the age-related up-regulation of vimentin is mostly due to expansion of fibroblast population in left ventricle as shown in Fig. 1E and Fig. 1F). Confocal co-immunofluorescence staining shows that vimentin could be found in CD31^+^ endothelial cells of blood vessels in the heart of old *wt* but rarely in age-matched *arg-ii^−/−^* mice (**Fig. 4E**). The results suggest at least a partial End-MT occurring in cardiac aging that is prevented by *arg-ii* ablation. With aging, the heart weight to body weight ratio (HW/BW) increases to a similar extent between *wt* and *arg-ii^−/−^* mice (**Suppl. Fig. 3**), suggesting no difference in age-related cardiac hypertrophy between *wt* and *arg-ii^−/−^*mice.

**Fig. 4.**
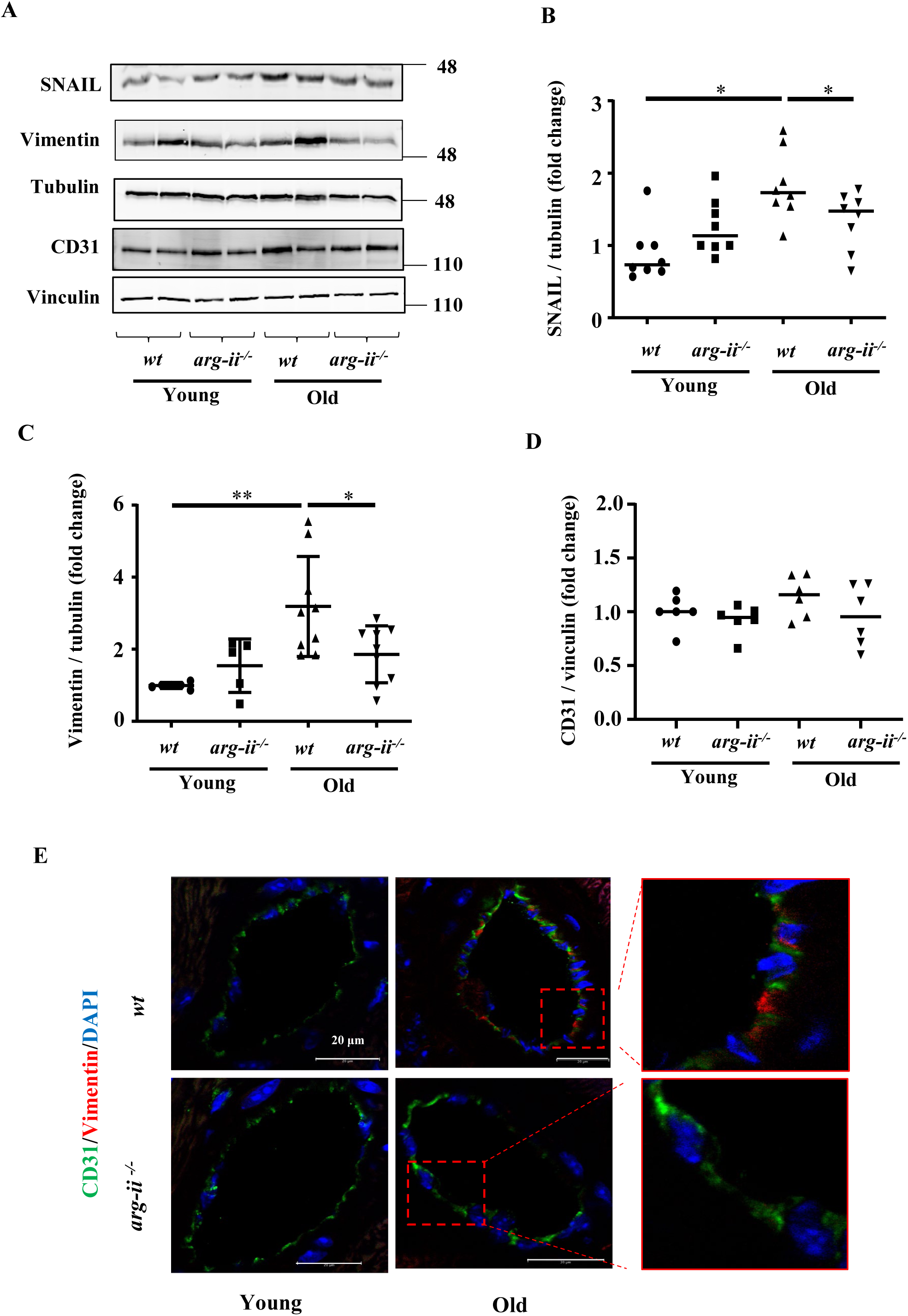
Age-related endothelial-to-mesenchymal transition (EndMT) in heart tissue. (**A**) Immunoblotting analysis of CD31 (endothelial marker), vimentin (mesenchymal marker) and SNAIL (master regulator of EndMT) in the heart of *wt* and *arg-ii^−/−^* female young and old mice; tubulin and vinculin served as protein loading controls. Molecular weight (kDa) is indicated at the side of the blots. The plot graphs show the quantification of the SNAIL (**B**), vimentin (**C**) and CD31 (**D**) signals on immunoblots (n=6-10 mice in each group); (**E**) Representative confocal images showing co-localization of CD31 (green) and vimentin (red) in young and old *wt* and *arg-ii^−/−^* heart tissues. DAPI (blue) is used to stain nuclei. Scale bar = 20 µm. *p≤0.05 and **p ≤ 0.01, between the indicated groups. *wt*, wild-type mice; *arg-ii^−/−^*, *arg-ii* gene knockout mice.

### 3. Cellular localization of Arg-II in aging heart

Next, the cellular localization of Arg-II in aging heart are investigated. Since *arg-ii* mRNA levels are upregulated mostly in aged females, heart tissues from old female mice are used mainly for this purpose. To our surprise, confocal co-immunofluorescence staining does not show Arg-II in cardiomyocytes as evidenced by lack of Arg-II staining in troponin T^+^-cells, a specific cardiomyocyte marker (**Fig. 5A**). Arg-II is only found in non-cardiomyocytes (**Fig. 5A**). In line with this notion, Arg-II is not detectable by immunoblotting in isolated primary cardiomyocytes from old *wt* mice at baseline or after exposure to hypoxia (1 % O_2_, 24 hours, **Fig. 5B**), a well know strong stimulus for Arg-II expression in many cell types (***Liang et al., 2019; Liang et al., 2021; Ren et al., 2022; Zhu et al., 2023***). In contrast to cardiomyocytes, cardiac fibroblasts isolated from old *wt* mice express Arg-II which is up-regulated by hypoxia (**Fig. 5B**). The absence of *arg-ii* mRNA in cardiomyocytes in contrast to fibroblasts isolated from *wt* old mouse hearts is also confirmed with qRT-PCR (**Fig. 5C**). The absence of Arg-II in cardiomyocytes are also demonstrated in the heart of other species such as rats in which no staining could be found in the cardiomyocytes (with or without infarction) but in non-cardiomyocytes i.e., cells between myocytes and endocardial cells (**Suppl. Fig. 4B**). The specificity of the antibody against Arg-II was confirmed in the old *wt* and *arg-ii^−/−^* mouse kidneys in which Arg-II is constitutively expressed in the S3 proximal tubular cells (**Suppl. Fig. 4A**) as demonstrated by our previous study (***Huang et al., 2016***). Again, Arg-II staining could be only found in cells between cardiomyocytes in the old *wt* mouse but not in *arg-ii^−/−^* animals (**Suppl. Fig. 4C**). The absence of Arg-II in cardiomyocytes are also confirmed in human heart tissue biopsies (please see the results in **Fig. 5H**). Moreover, Arg-II is detected in Mac-2^+^ macrophages (**Fig. 5D**), CD31^+^ endothelial cells (**Fig. 5E**), and PDGF-Rα^+^-fibroblasts (**Fig. 5F**), but not in α-SMA^+^ vascular smooth muscle cells (**Fig. 5G).** These results demonstrate that Arg-II in aging mouse heart is mostly expressed in non-cardiomyocytes such as macrophages, endothelial cells and fibroblasts. Similar to mouse heart, in human heart (obtained from a 66-year-old woman, and a 76-year-old man with no significant cardiac pathology), Arg-II is also absent in cardiomyocytes as proved by lack of co-localization with troponin-T^+^-cells (**Fig. 5H**), and is expressed in CD31^+^-endothelial cells (**Fig. 5I**), CD68^+^-macrophages (**Fig. 5J**), and vimentin^+^-(**Fig. 5K**) or α-SMA^+^-(**Fig. 5L**) fibroblasts/myofibroblasts. In contrast to mouse heart, Arg-II could be found in very few vascular smooth muscle cells in human heart (**Fig. 5M**).

**Fig. 5.**
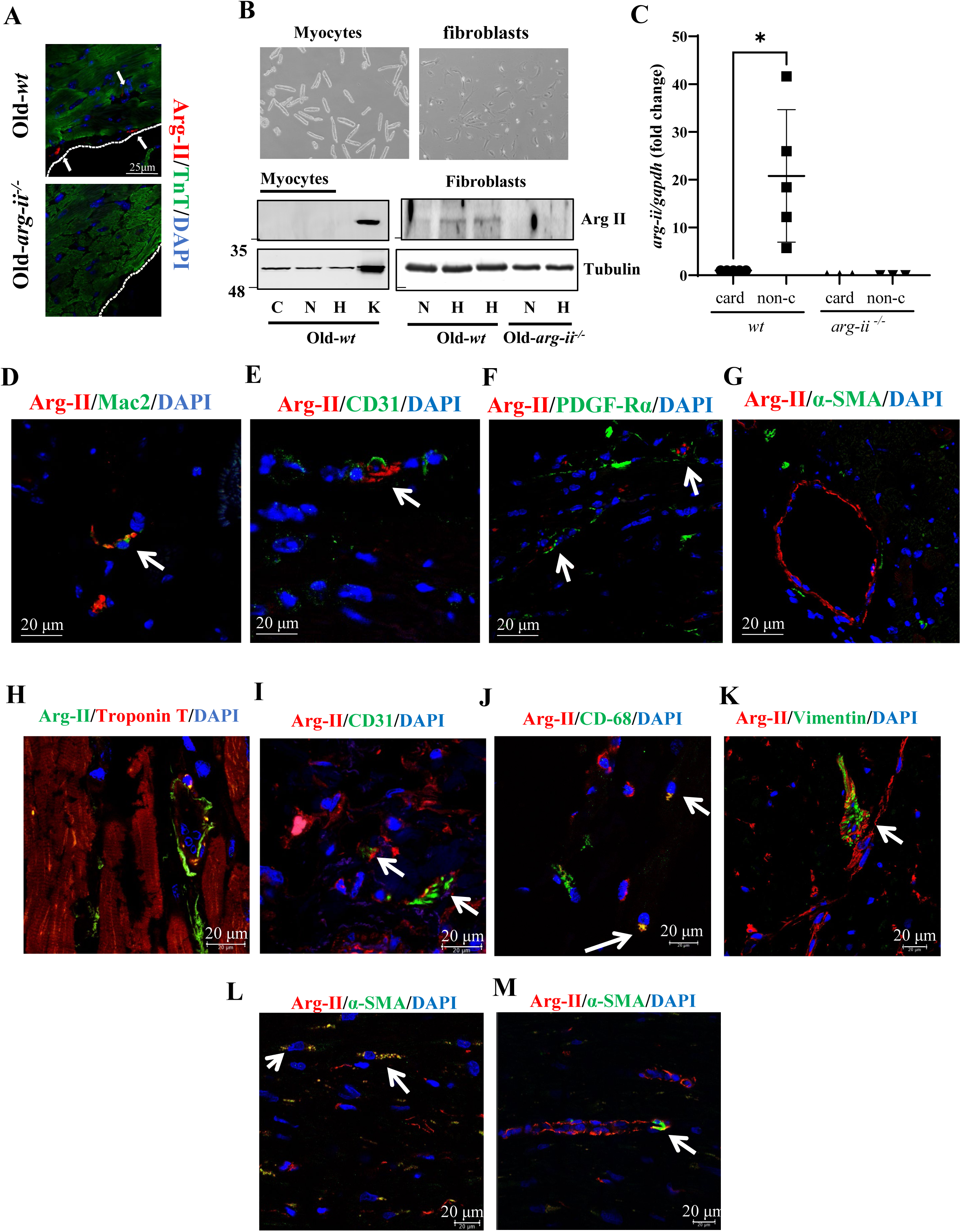
Cellular localization of Arg-II in aging heart of female mice. (**A**) Confocal microscopy illustration of immunofluorescence double staining of Arg-II (red) and tropoin-t (TnT, green; cardiomyocytes marker). Scale bar = 25 µm; (**B**) Bright-field microscopy images of isolated *wt* cardiomyocytes and primary cardiac fibroblasts. The immunoblot shows the level of Arg-II in both cardiomyocytes and fibroblasts upon exposure to hypoxia (1 % O_2_) for 24 h. C indicates freshly isolated cardiomyocytes used as control, N and H indicates respectively normoxia and hypoxia conditions, and K indicates kidney tissue extract used as positive control. Tubulin served as protein loading control; (**C**) mRNA expression levels of *arg-ii* in cardiomyocytes (card) and non-cardiomyocytes (non-c) cells isolated from old *wt* and *arg-ii^−/−^* female mouse hearts. *gapdh* served as the reference. (n=3-5 mice per group); (**D** to **G**) Representative confocal images of old *wt* mouse heart showing co-localization of (**D**) Arg-II (red) and Mac-2 (green, mouse macrophage marker), (**E**) Arg-II (red) and CD31 (green, endothelial marker), (**F**) Arg-II (red) and PDGF-Rα (green, fibroblasts marker), and (**G**) Arg-II and α-smooth muscle actin (α-SMA; green, smooth muscle cell/myofibroblasts marker); (**H** to **M**) Representative confocal images of human heart tissue showing co-localization of (**H**) Arg-II (green) and tropoin-t (red, cardiomyocytes marker), (**I**) Arg-II (red) and CD31 (green, endothelial marker), (**J**) Arg-II (red) and CD-68 (green, macrophage marker), (**K**) Arg-II (red) and vimentin (green, fibroblast marker), and (**L-M**) Arg-II (red) and α-smooth muscle actin (α-SMA; green, (**L**) myofibroblasts and (**M**) smooth muscle cell marker). DAPI (blue) stains cell nuclei. Scale bar = 20 µm. Each experiment was repeated with 3 to 5 animals.

### 4. Role of Arg-II in crosstalk between macrophages and cardiomyocytes through IL-1β

Given that age-associated cardiomyocyte apoptosis is prevented in *arg-ii^−/−^* mice despite absence of Arg-II in the cardiomyocytes, the non-cell-autonomous effects of Arg-II on cardiomyocytes are then investigated. As reported above, macrophage numbers and *il-1β* mRNA expression are increased in aging heart and reduced in *arg-ii^−/−^* mice (Fig. 2A, 2C, and 2G). Both immunoblotting and immunofluorescence staining confirm that IL-1β protein levels are higher in old *wt* heart tissues when compared to age-matched *arg-ii^−/−^*(**Fig. 6A** to **6D**). Furthermore, co-immunofluorescence staining reveals that IL-1β is localized in Mac2^+^-macrophages (**Fig. 6E**). Taking into account that our previous studies demonstrated a relationship of Arg-II and IL-1β in vascular disease and obesity (***Ming et al., 2012***) and in age-associated organ fibrosis such as renal and pulmonary fibrosis (***Huang et al., 2021; Zhu et al., 2023***), and IL-1β has been shown to play a causal role in patients with coronary atherosclerotic heart disease as shown by CANTOS trials (***Ridker et al., 2017***), we therefore focused on the role of IL-1β in crosstalk between macrophages and cardiac cells such as cardiomyocytes, fibroblasts and endothelial cells. Considering that ablation of *arg-ii* gene specifically reduces the number of TUNEL-positive cardiomyocytes (Fig. 3C and 3D), we hypothesized that Arg-II in macrophages may prime cardiomyocyte apoptosis. To test this hypothesis, splenic macrophages are isolated from young and old *wt* and age-matched *arg-ii^−/−^* mice, and conditioned medium from the cells are collected. Elevated Arg-II and IL-1β protein levels are observed in old *wt* macrophages as compared to the cells from young animals (**Fig. 7A** to **7C**). In line with this finding, IL-1β levels in conditioned medium from old *wt* mouse macrophages are higher as compared to the conditioned medium from young macrophages (**Fig. 7D**). Interestingly, *arg-ii* ablation substantially reduces IL-1β levels in the macrophages (**Fig. 7A, 7C,** and **7D**). Importantly, cardiomyocytes isolated from adult (five-months) female *wt* mice as bioassay cells, when incubated with conditioned medium from old (but not from young) *wt* macrophages for 24 hours (**Fig. 7E**), display more apoptosis as monitored by TUNEL assay (**Fig. 7F** and **7G**). This age-associated cardiac damaging effect of macrophages is not observed with *arg-ii^−/−^* macrophage conditioned medium (**Fig. 7F** and **7G**), suggesting a role of Arg-II expressing macrophages in mediating cardiomyocyte apoptosis in aging. To further examine whether IL-1β from macrophages is the paracrine mediator for cardiomyocyte apoptosis, the above described experiment is performed in presence of the IL-1 receptor antagonist IL-Ra. Indeed, cardiomyocyte apoptosis induced by the conditioned medium from old *wt* macrophages is abolished by IL-Ra (**Fig. 7F** and **7G**).

**Fig. 6.**
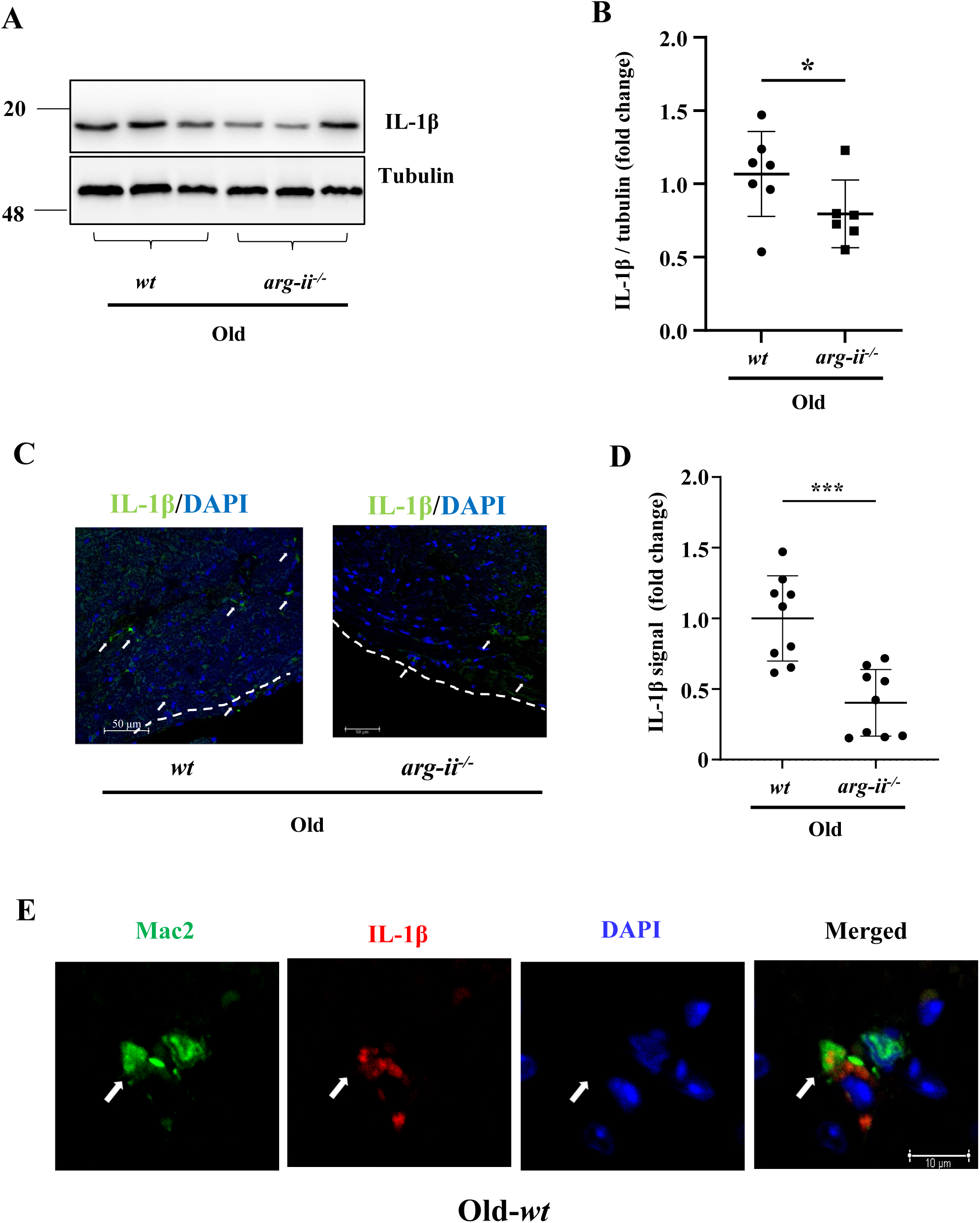
Arg-II ablation reduces Il-1β protein levels in aging heart. (**A**) Immunoblotting analysis of Il-1β (active form) in the heart of old *wt* and *arg-ii^−/−^* female mice; tubulin served as protein loading control. Molecular weight (kDa) is indicated at the side of the blots; (**B**) The plot graph shows quantification of the Il-1β signals on immunoblots (n=6-7 mice in each group); (**C**) Representative confocal images showing Il-1β localization in old *wt* and *arg-ii^−/−^* heart tissues. DAPI (blue) is used to stain nuclei. Scale bar = 50 µm; (**D**) Relative Il-1β signal quantification of confocal images (n= 9 per each group); (**E**) Representative confocal images showing co-localization of Mac-2 (green, mouse macrophage marker), and Il-1β (red) in *wt* heart tissues. This experiment was repeated with 3 animals. Scale bar = 10 µm. *p≤0.05 and ***p ≤ 0.005, between the indicated groups. *wt*, wild-type mice; *arg-ii^−/−^*, *arg-ii* gene knockout mice.

**Fig. 7.**
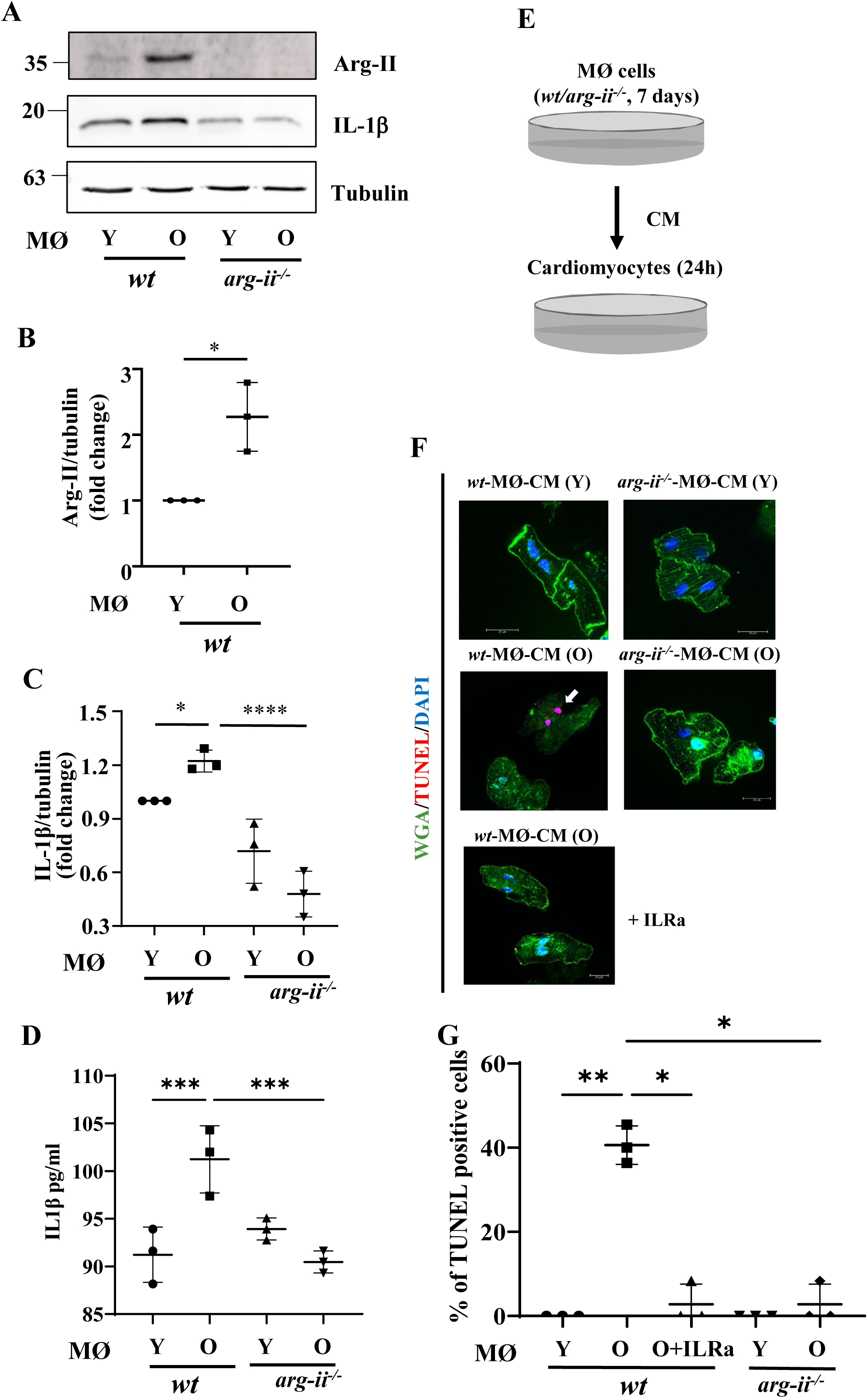
*In vitro* study of crosstalk between aging splenic macrophages and cardiomyocytes. (**A**) Immunoblotting analysis of Arg-II and Il-1β (active form) in mouse splenic macrophages isolated from young and old *wt* and *arg-ii^−/−^* female mice; tubulin served as protein loading control. Molecular weight (kDa) is indicated at the side of the blots. The plot graphs show the quantification of Arg-II (**B**) and Il-1β (**C**) protein signals on immunoblots (n=3 mice in each group); (**D**) IL-1β levels in the conditioned medium from young and old *wt* and *arg-ii^−/−^* splenic macrophages was measured by ELISA; (**E**) Schematic representation of *in vitro* crosstalk study; (**F**) Representative confocal images of adult isolated mouse cardiomyocytes stimulated with conditioned media (CM) from young and old, *wt* and *arg-ii^−/−^* splenic cells (24 h incubation). Wheat Germ Agglutinin (WGA)-Alexa Fluor 488-conjugate was used to stain cell membrane, and TUNEL was performed to identify apoptotic cells. DAPI is used to stain nuclei. Interleukin receptor antagonist (ILRa; 50 ng/mL) is used to prevent IL-1β binding to its receptor; (**G**) Quantification of TUNEL-positive cardiomyocytes (% in respect to total number of cells). Scale bar = 20 µm. *p≤0.05, **p ≤ 0.01, ***p ≤ 0.005 and ****p ≤ 0.001 between the indicated groups. MØ, splenic macrophage; Y, young; O, old; *wt*, wild-type mice; *arg-ii*^−/−^, *arg-ii* gene knockout mice.

Similar results are obtained with Raw264.7 cells, a widely used mouse macrophage model. Raw264.7 cells stimulated with lipopolysaccharide (LPS; 100 ng/mL) for 24 hours, show a significant increase in both Arg-II and IL-1β, which is inhibited by silencing *arg-ii* (**Suppl. Fig. 5A** to **Suppl. Fig. 5C**), confirming the role of Arg-II in the production of IL-1β in macrophages (**Suppl. Fig. 5A** and **Suppl. Fig. 5C**). In agreement with the results shown in Fig. 7, conditioned medium from LPS-treated Raw264.7 cells significantly enhances cardiomyocyte apoptosis as assessed by TUNEL staining, which is reduced by either *arg-ii* gene silencing or by IL-1 receptor blocker ILRa (**Suppl. Fig. 5D** and **Suppl. Fig. 5E**). It is of note that that LPS alone does not induce cardiomyocyte apoptosis at the concentration tested (data not shown), confirming the role of Arg-II-mediated IL-1β release from macrophages in cardiomyocyte apoptosis.

Next, we analyzed relationship between Arg-II and iNOS in regulation of IL-1β production in macrophages. In the human THP1 monocytes in which Arg-II but not iNOS is induced by LPS (100 ng/mL for 24 hours) (**Suppl. Fig. 6A**), mRNA and protein levels of IL-1β are markedly reduced in *arg-ii* knockout THP1*^arg-ii-/-^* as compared to the THP1*^wt^* cells (**Suppl. Fig. 6B and 6C**), further confirming that Arg-II promotes IL-1β production as already shown in RAW264.7 macrophages (Suppl. Fig. 5A and 5C). Moreover, in the mouse bone-marrow-derived macrophages, LPS-induced IL-1β production is inhibited by *inos* deficiency (BMDM*^inos-/-^* vs BMDM*^wt^*) (**Suppl. Fig. 6D and 6E**), while Arg-II levels are slightly enhanced in the BMDM*^inos-/-^* cells (**Suppl. Fig. 6D and 6F**). All together, these results suggest that Arg-II and iNOS are upregulated by LPS independently and iNOS slightly reduces Arg-II expression. Both Arg-II and iNOS are required for maximal IL-1β production upon LPS stimulation as illustrated in **Suppl. Fig. 6G**.

### 5. Role of Arg-II in crosstalk between macrophages and cardiac fibroblasts through IL-1β

Since cardiac fibrosis is accompanied with macrophage accumulation in our aging heart model, a crosstalk between macrophages and fibroblasts is then investigated. Cardiac fibroblasts isolated from adult (five-months) female *wt* mice as bioassay cells, produce more hydroxyproline when they are incubated with conditioned medium from old as compared to young *wt* macrophages (**Suppl. Fig. 7A**). This effect of old macrophages on fibroblasts is reduced with conditioned medium from *arg-ii^−/−^* macrophages (**Fig. 8A**). In parallel, expression levels of genes involved in extracellular matrix remodeling such as *fibronectin, collagen IIIα, tgf-β1* (**Fig. 8B** to **Fig. 8D**) as well as *mmp2* and *mmp9* (**Suppl. Fig. 7B** and **Suppl. Fig. 7C**) are reduced in fibroblasts stimulated with conditioned medium from *arg-ii^−/−^*macrophages as compared to the old *wt* macrophages, while expression of *collagen Iα* is not altered under this condition (**Suppl. Fig. 7D**). Importantly, the increase in hydroxyproline levels in cardiac fibroblasts incubated with conditioned medium from old *wt* macrophages is inhibited by ILRa (**Fig. 8E**).

**Fig. 8.**
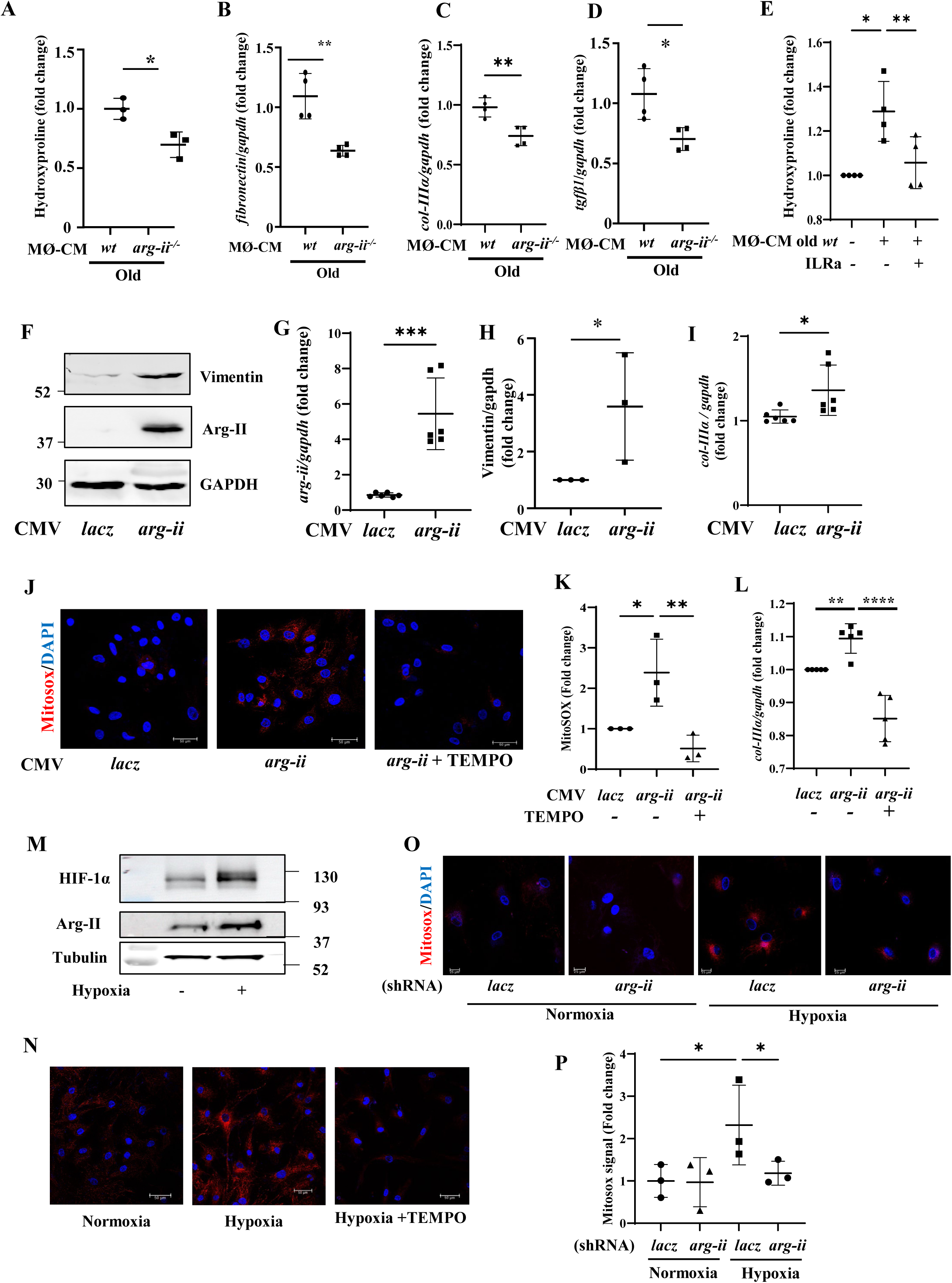
Crosstalk between splenic macrophages and cardiac fibroblasts and cell-autonomous effect of Arg-II in cardiac fibroblasts. (**A**) Collagen production measured as hydroxyproline content in mouse *wt* fibroblasts treated with conditioned media (CM) from old, *wt* and *arg-ii^−/−^* splenic cells (96 h incubation) (n=3 mice in each group). The values shown are mean ± SD. mRNA expression levels of (**B**) *fibronectin*, (**C**) *collagen-IIIα (col-IIIα*) and (**D**) *tgf-β1* in *wt* fibroblasts treated with conditioned media (CM) from old *wt* and *arg-ii^−/−^* splenic cells (96 h incubation) were analyzed by qRT-PCR. *gapdh* served as the reference. (n=4 mice per group); (**E**) Hydroxyproline content in fibroblasts treated with CM from old *wt* splenic cells (96 h incubation). ILRa (50 ng/mL) is used to prevent IL-1β binding to its receptor (n=4 independent experiments). (**F**) Immunoblotting analysis of Arg-II and vimentin in human cardiac fibroblasts (HCFs) upon *arg-ii* gene overexpression. GAPDH served as protein loading control; (**G**) qRT-PCR analysis of mRNA expression levels *arg-ii* in HCF cells; (**H**) The plot graph shows the quantification of the vimentin signals on immunoblots shown in panel F. (n=3 independent experiments); (**I**) qRT-PCR analysis of mRNA expression levels of *collagen-IIIα* in HCF cells. *gapdh* served as the reference. (n=6 independent experiments); (**J**) Representative confocal images of human cardiac fibroblasts (HCF) upon transfection with rAd-CMV-Con/Arg-II for 48 h. MitoSOX (Red) is used to stain mitochondrial ROS (mtROS). TEMPO (10 μmol/L) is used to prevent mtROS generation. DAPI (blue) stains cell nuclei. Scale bar = 50 µm. (**K**) Mitosox signal quantification (n=3 independent experiments); (**L**) qRT-PCR analysis of mRNA expression levels of *col-IIIα* in HCF cells treated as indicated. *gapdh* served as the reference. (n=5 independent experiments). (**M**) Immunoblotting analysis of Arg-II and Hif-1α in human cardiac fibroblasts (HCFs) upon 1% hypoxia incubation for 48h. Tubulin served as protein loading control; (**N**) Representative confocal images of HCFs under normoxia (21% O_2_) or hypoxia (1% O_2_) for 48 h. MitoSOX (Red) is used to stain mitochondrial ROS (mtROS). TEMPO (10 μmol/L) is used to prevent mtROS generation. DAPI (blue) stains cell nuclei. Scale bar = 50 µm. (**O**) Representative confocal images of HCFs upon transfection with shRNA for *arg-ii* gene silencing under normoxia or hypoxia. MitoSOX (Red) is used to stain mitochondrial ROS (mtROS). DAPI (blue) stains cell nuclei. Scale bar = 25 µm. (**P**) MitoSOX signal quantification of images shown in panel O (n=3 independent experiments). Data are expressed as fold change to respective control group. *p≤0.05, **p ≤ 0.01, ***p ≤ 0.005 and ****p ≤ 0.001 between the indicated groups. MØ, splenic macrophage; Con, control.

### 6. Cell-autonomous effects of Arg-II in cardiac fibroblasts: role of mtROS

Since Arg-II is expressed in fibroblasts, a cell-autonomous effect of Arg-II in these cells is also investigated in cultured human cardiac fibroblasts (HCF). Overexpression of *arg-ii* in the cells (**Fig. 8F nd 8G**) increases *vimentin* and *collagen IIIα* expression levels (**Fig. 8H and 8I),** demonstrating a cell-autonomous effect of Arg-II in cardiac fibroblasts. Interestingly, upregulation of *collagen IIIα* expression by *arg-ii* is accompanied with increased mitochondrial ROS (mtROS) generation, and inhibition of mtROS by mito-TEMPO (10 μmol/L) prevents *col-IIIa* upregulation (**Fig. 8J, 8K and 8L**), suggesting a role of Arg-II-induced mtROS in activation of fibroblasts. The effect of Arg-II in promoting mtROS is also confirmed under hypoxic conditions in which endogenous Arg-II is upregulated accompanied with elevated HIF-1α levels (**Fig. 8M and 8O**). The enhanced mtROS generation under hypoxic conditions are inhibited either by mito-Tempo (**Fig. 8N**), the specific inhibitor of mtROS or by *arg-ii* silencing (**Fig. 8O and 8P**).

### 7. Role of Arg-II in crosstalk between macrophages and endothelial cells through IL-1β

Results in Fig. 4 suggests EndMT in aging heart, which is regulated by Arg-II. The role of Arg-II-expressing macrophages in End-MT of endothelial cells is then investigated in cultured cells. Human endothelial cells exposed to macrophage-derived conditioned medium from old (not young) *wt* mice for 96 hours has no effect on VE-cadherin but enhances N-cadherin and vimentin (mesenchymal markers), and also Arg-II protein levels (**Fig. 9A** to **Fig. 9E**), suggesting a partial End-MT induced by aged macrophages. Interestingly, these effects of macrophages are prevented in old *arg-ii^−/−^* mice (**Fig. 9A** to **Fig. 9E**) or abolished in the presence of IL-1 receptor antagonist IL-Ra (**Fig. 9F** to **Fig. 9J**). The results suggest a role of IL-1β from old *wt* macrophages in EndMT process.

**Fig. 9.**
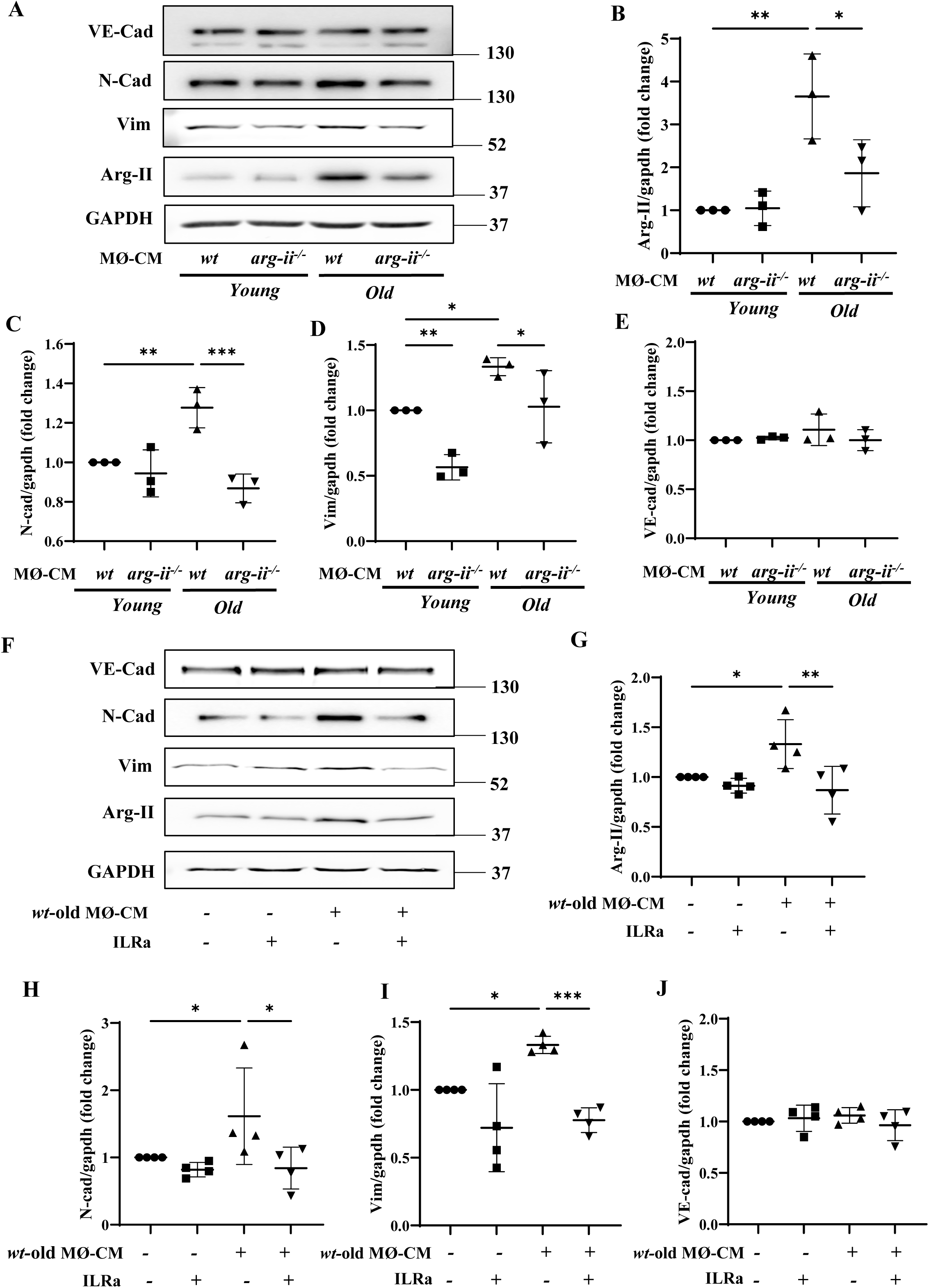
Aging *wt* macrophages induces EndMT in HUVEC. (**A**) Immunoblotting analysis of VE-Cadherin (endothelial marker), N-Cadherin and vimentin (both mesenchymal markers), and Arg-II in HUVEC cells upon incubation with conditioned media (CM) from young and old *wt* and *arg-ii^−/−^*splenic cells (4 days incubation); GAPDH served as protein loading control. Molecular weight (kDa) is indicated at the side of the blots. The plot graphs show the quantification of Arg-II (**B**), N-Cadherin (**C**), vimentin (**D**), and VE-Cadherin (**E**) protein levels (n=3 independent experiments); (**F**) Immunoblotting analysis of VE-Cadherin, N-Cadherin, vimentin and Arg-II in HUVEC cells upon incubation with CM from old *wt* splenic cells (4 days incubation); Interleukin receptor antagonist (IL-Ra; 50 ng/mL) is used to prevent IL-1β effect. GAPDH served as protein loading control. Molecular weight (kDa) is indicated at the side of the blots. The plot graphs show the quantification of Arg-II (**G**), N-Cadherin (**H**), vimentin (**I**), and VE-Cadherin (**J**) protein signals on immunoblots (n=4 independent experiments); Data are expressed as fold change to respective control group. *p≤0.05, **p ≤ 0.01 and ***p ≤ 0.005 between the indicated groups. MØ, splenic macrophage; *wt*, wild-type mice; *arg-ii^−/−^*, *arg-ii* gene knockout mice.

### 8. Arg-II gene ablation reduces ischemic damage in aging heart

To study the specific role of Arg-II in heart ischemia/reperfusion (I/R) injury during aging, *ex vivo* experiments with Langendorff-perfused heart from old *wt* and *arg-ii*^−/−^ mice were performed followed by cardiac functional analysis. The experimental protocol of global ischemia/reperfusion is shown in **Fig. 10A**. Under normoxic baseline condition, the cardiac functions, i.e., dP/dt_max._ (maximal rate of contraction), dP/dt_min._ (maximal rate of relaxation) and left ventricular developed pressure (LVDP) are comparable between *wt* and *arg-ii*^−/−^ old mice (**Suppl. table 1**). *Arg-ii* ablation improves cardiac function following I/R injury, i.e., the inotropic contractile function as measured by LVDP and dP/dt_max_, lusitropic dP/dt_min_ (**Fig. 10B**). The post I/R recovery is significantly improved in *arg-ii*^−/−^ mice when compared to age-matched *wt* animals (**Fig. 10B**). In addition to functional recovery, infarct size after I/R-injury was analyzed by triphenyl tetrazolium chloride (TTC) staining in old mice. Under the ischemic/reperfusion condition (20 minutes ischemia and 30 minutes reperfusion), no myocardial infarct was detected (**Suppl. Fig. 8**), while longer ischemia and reperfusion time (30 minutes and 45 minutes respectively) is required to induce myocardial infarct, i.e., 10.5 ± 2.9% in *wt* old mice, which is significantly reduced in the age-matched *arg-ii^−/−^*mice (3.2 ± 0.9%, **Fig. 10C** and **Fig. 10D**). It is to note that following ischemia/reperfusion injury, male mice display cardiac dysfunction, i.e., reduced left ventricular developed pressure (LVDP), as well as the inotropic and lusitropic states (expressed as dP/dt max and dP/dt min, respectively) (**Suppl. Fig. 9**). As previously reported (***Murphy et al., 2007***), we also found that old male mice are more prone to I/R injury than age-matched female animals. Specifically, 15 minutes of ischemic time is enough to significantly affect the left ventricle contractile function in the male mice (**Suppl. Fig. 9**). As opposite, age-matched old female mice are relatively resistant to I/R injury, and at least 20 min of ischemia are necessary to induce a significant impairment of the contractile function (**Fig. 10**). Similar to females, the post I/R recovery of cardiac function is also significantly improved in the male *arg-ii^−/−^* mice as compared to age-matched *wt* animals. In addition to functional recovery, triphenyl tetrazolium chloride (TTC) staining (myocardial infarction) upon I/R-injury in males is significantly reduced in the age-matched male *arg-ii^−/−^* animals (**Suppl. Fig. 9C** and **9D**). All together, these results reveal a role for Arg-II in heart function impairment during aging in both genders with a higher vulnerability in males (**Suppl. Fig. 9**).

**Fig. 10.**
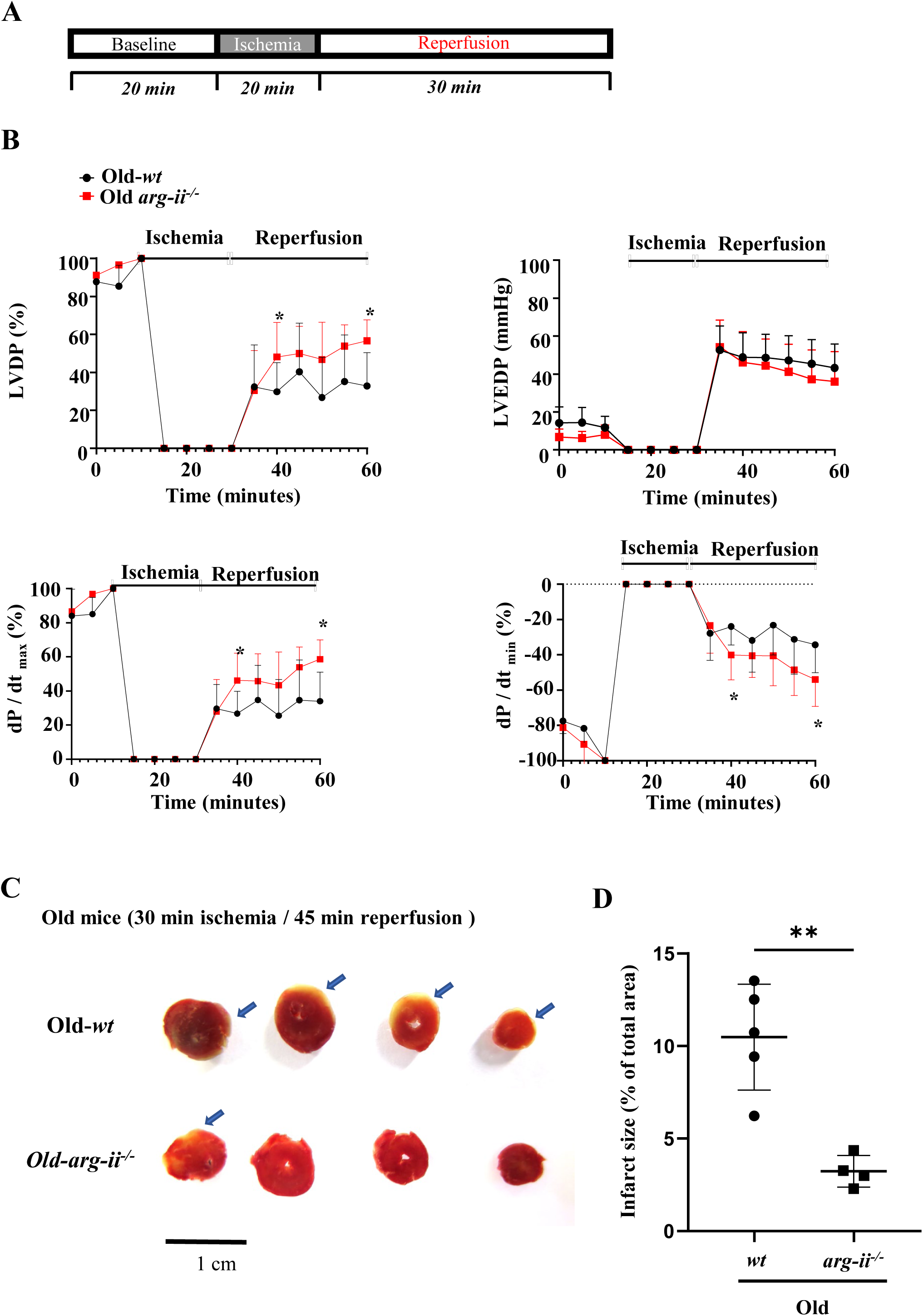
Arg-II ablation improves heart function recovery from global ischemia/reperfusion injury (I/R-I). (**A**) Protocol applied for e*x vivo* Langendorff-heart functional assessment of old *wt* and age-matched *arg-ii^−/−^* female mice. After baseline recordings, 20 minutes global ischemia is followed by 30 minutes of reperfusion; (**B**) *Ex vivo* Langendorff-heart assessment of old female *wt* (black line) and *arg-ii^−/−^* (red line) hearts. The graphs show the functional recovery of the left ventricular developed pressure (LVDP), left ventricular end diastolic pressure (LVED) and maximal rate of contraction (dP/dt_max_) and relaxation (dP/dt_min_). The data are expressed as % of recovery in respect to baseline values and represent the mean ± SD of data from 5 mice per group; (**C**) Representative sections of *wt* and *arg-ii^−/−^* female hearts stained with 2,3,5-TTC. The healthy/normal tissue appear deep red. The arrows indicate the white infarct tissue showing the absence of living cells. (**D**) relative quantification of infarcted areas. *p≤0.05 and **p ≤ 0.01 between the indicated groups. *wt*, wild-type mice; *arg-ii^−/−^*, *arg-ii* gene knockout mice.

## Discussion

Previous studies have reported the presence of both Arg-I and Arg-II isoenzymes in the cardiac tissues of various animal models including feline, pig, rats, rabbit, and mice (***Gonon et al., 2012; Heusch et al., 2010; Jung et al., 2006; Jung et al., 2010; Steppan et al., 2006***). These isoenzymes have been implicated in cardiac ischemic injury and cardiac aging (***Jung et al., 2010; Xiong, Yepuri, Montani, et al., 2017***). It has been widely assumed that Arg-I and Arg-II are expressed in cardiomyocytes, where they play a role in cardiac injury via a cell-autonomous effect on the cardiomyocytes. However, the cellular localization of arginase isoforms has not been convincingly demonstrated due to the following two reasons: species-specific expression of arginase isoforms and lack of including negative controls with gene knockout samples.

Our current study systematically investigated the expression and cellular localization of the mitochondrial Arg-II in the heart of a natural aging mouse model and in human biopsies. We found an age-associated increase in *arg-ii* but not *arg-i* in heart of male and female mice. Data extracted from a high throughput sequencing database of male mice (no data on females are available, unfortunately) confirms our findings showing the age-dependent enhancement of *arg-ii* but not *arg-i* expression in the heart (GSE201207) (***Wolff et al., 2023***) (**Suppl. Fig. 1C and 1D**). Surprisingly, we found that neither *arg-ii* gene expression nor its protein levels were detectable in cardiomyocytes from either young or old mice. Despite increased *arg-ii* mRNA (but not *arg-i* mRNA) expression in the aging heart, mitochondrial Arg-II was absent from cardiomyocytes. This observation aligns with previous findings in vascular endothelial cells from mice and humans, where Arg-II (not Arg-I) is the predominant isoenzyme (***Ming et al., 2004***). It is to note that a previous study reported that Arg-II is exclusively expressed in isolated cardiomyocytes from in rats (***Steppan et al., 2006***). However, we could not confirm this finding. The reason of this discrepancy is not clear. However, we made great effort to confirm the cellular localization of Arg-II in the heart from different species including mice, rats and humans. Importantly negative controls. i.e., *arg-ii^−/−^* samples are always included in our present study avoid any possible background signals. Our experiments confirm that Arg-II could be found only in non-cardiomyocytes but not in cardiomyocytes from different species. The experiments with freshly isolated primary cells also confirm that even under hypoxia, a well know strong stimulus for Arg-II protein levels in different cell types (***Liang et al., 2019; Zhu et al., 2023***), no Arg-II protein expression could be detected in the cardiomyocytes but elevated in fibroblasts from old female mice. Furthermore, RT-qPCR could not detect *arg-ii* mRNA in cardiomyocytes but in non-cardiomyocytes. All together, these results demonstrate that Arg-II are not expressed or at negligible levels in cardiomyocytes but expressed in non-cardiomyocytes. Further experiments identified that Arg-II is expressed in macrophages, fibroblasts, and endothelial cells in aged mouse heart and human cardiac biopsies. These results suggest that Arg-II exerts non-cell-autonomous effects on aging cardiomyocytes.

It is well known that the aging heart is associated with accumulation of senescent cells, increased cardiomyocyte apoptosis, chronic inflammation, oxidative stress, and cardiac fibrosis (***Vakka et al., 2023***). In the current study, we observe that *arg-ii^−/−^* mice are protected from cardiac aging phenotype and reveal a decrease in apoptotic cells, particularly cardiomyocytes but also non-cardiomyocytes. Since cardiomyocytes do not express Arg-II in aging and Arg-II could not even be induced in these cells under hypoxic condition, a non-cell-autonomous effect of Arg-II on cardiomyocyte apoptosis must be involved, meaning that the increased cardiomyocyte apoptosis in aging must be due to paracrine effects of other cell types such as macrophages in which Arg-II levels are elevated in aging. It is evident that crosstalk between cardiomyocytes and non-cardiomyocytes are critical in cardiac functional changes and remodeling (***Hulsmans et al., 2016***). Substantial evidence demonstrates that immune cells, particularly, monocytes and/or macrophages, are important contributors to cardiac remodeling in aging and disease conditions (***Hulsmans et al., 2016; Jimenez et al., 2022; Wang et al., 2020***). Under acute and chronic ischemic heart disease conditions, blood monocytes can be mobilized from bone marrow and spleen, and recruited into the injury site of the heart where they play an important role in cardiac remodeling (***Honold et al., 2018; Swirski et al., 2009***). An increased accumulation of myeloid cells including in aging heart has been reported and contributes to cardiac inflammaging (***Esfahani et al., 2021***). Besides infiltrated macrophages, tissue resident macrophages also contribute to cardiac inflammation and remodeling in heart disease (***Suku et al., 2022; Weinberger et al., 2024***). Interestingly, we demonstrate that total numbers of macrophages in the heart are significantly increased in aging heart, which is due to both enhanced accumulation of resident tissue macrophages and infiltrated macrophages and both macrophage populations are reduced in *arg-ii^−/−^* mice. The enhanced infiltrated macrophages must be due to sustained cardiac injury occurring with aging in the heart accompanied with repairing process which is manifested by resident macrophage proliferation and fibroblast activation (***Weinberger et al., 2024***). Our previous study demonstrated that Arg-II plays a role in macrophage infiltration in different organ/tissues in high-fat diet and high-cholesterol diet-induced atherosclerosis and obesity mouse models (***Ming et al., 2012***). It remains to be investigated whether Arg-II regulates resident macrophage proliferation in the aging heart and what are the underlying mechanisms. Whether infiltrated macrophages can be switched to resident ones and regulated by Arg-II is also an interesting question to be investigated.

As shown above an increased macrophage accumulation in aging heart associates with elevated expression of numerous inflammatory cytokines in macrophages. This age-associated cardiac inflammation is inhibited in *arg-ii^−/−^* animals. Among these inflammatory cytokines, an increase in IL-1β levels in cardiac tissue and macrophages or monocytes/macrophages derived from spleen of old mice are demonstrated accompanied with elevated Arg-II levels as compared to young animals. Importantly, we showed that Arg-II in the aged macrophages enhances IL-1β expression and release mediating cardiomyocyte apoptosis, since the cardiomyocyte apoptosis caused by the aged macrophages is inhibited in *arg-ii^−/−^*mice and prevented by IL-1β receptor antagonist. This results further confirm our previous findings demonstrating a pro-inflammatory role of Arg-II in macrophages in vascular atherosclerosis and high-fat-diet-induced type 2 diabetes mouse models (***Ming et al., 2012***). This conclusion is also further supported by the findings with a mouse macrophage cell line RAW264.7. The question what are the underlying mechanisms that upregulate Arg-II levels in macrophages in aging remains to be investigated. Moreover, we also found increased apoptosis of other cell types in aging heart, which is also prevented in *arg-ii^−/−^*mice. What are these cells, how this cell apoptosis is triggered, and what is the impact on age-associated cardiac vulnerability to stressors require further investigation. Moreover, research has provided evidence that cellular senescence including cardiomyocytes, endothelial cells, fibroblasts and immune cells all contributes to the cardiac aging and age-associated cardiac dysfunction (***Luan et al., 2024***). Future research shall investigate which cell types are protected from *arg-ii* deficiency from senescence and what are the underlying mechanisms.

One of the interesting questions is the relationship between Arg-II and iNOS in regulation of IL-1β production in macrophages. Both iNOS and Arg-II (not Arg-I) are highly induced in macrophages in response to several pro-inflammatory stimuli such as LPS, TNFα, and IFN-γ, etc. (***Bogdan, 2015; Ming et al., 2012***) and considered as pro-inflammatory M1-macrophage markers (***Martinez et al., 2014; Ming et al., 2012***). Since arginase and iNOS share the same metabolic substrate L-arginine, an increase in arginase levels is proposed to decrease intracellular L-arginine bioavailability for iNOS, and in turn to reduce pro-inflammatory responses of macrophages (***Chang et al., 1998***). Hence, arginase including the two isoforms Arg-I and Arg-II is regarded as anti-inflammatory (***Yang et al., 2014***). Arg-I is indeed associated with M2-anti-inflammatory macrophages (***Yang et al., 2014***), while Arg-II (but not Arg-I) along with iNOS is enhanced in pro-inflammatory macrophages upon LPS stimulation as shown in the previous study (***Ming et al., 2012***) and also confirmed in our present study. Importantly, knockout or knockdown of *arg-ii* gene reduces IL-1β production as shown in our present study and decreases pro-inflammatory cytokine levels in vitro and in vivo in various chronic inflammatory disease models and inflammaging (***Atawia et al., 2019; Ming et al., 2012; You et al., 2013***). These results demonstrate that Arg-II, in contrast to Arg-I, exerts pro-inflammatory functions in macrophages. Despite controversies published in the literature (***Bogdan, 2015***), iNOS is generally considered to exert pro-inflammatory effects in macrophages (***Eslami et al., 2017; Yao et al., 2022***). This is confirmed by our present study showing that LPS-stimulated IL-1β production in murine macrophages is inhibited when *inos* is ablated. Since Arg-II promotes IL-1β production and Arg-II levels are increased under *inos*-deficiency condition, one would expect an increased IL-1β production in *inos^−/−^* macrophages. This is however not the case. A strong inhibition of IL-1β production in *inos^−/−^* macrophages is observed. These results implicate that iNOS promotes IL-1β production independently of Arg-II and the inhibiting effect of IL-1β by *inos* deficiency is dominant and able to counteract Arg-II’s stimulating effect on IL-1β production. On the other hand, the question whether Arg-II promotes IL-1β production through iNOS could also be excluded due to the following reasons. First, in the human THP1 cells in which Arg-II but not iNOS is induced by LPS, ablation of *arg-ii* gene reduces IL-1β mRNA and protein levels; second, both Arg-I and Arg-II is believed to inhibits endothelial NOS-mediated NO synthesis, *arg-ii* deficiency would increase L-arginine bioavailability for NOS activity (***Durante et al., 2007***). Since iNOS induction by LPS is promoting IL-1β production as demonstrated in our current study, an increase in IL-1β production by *arg-ii* deficiency would be expected under this condition. We observe, however, a decreased IL-1β production in *arg-ii^−/−^* cells. Hence, our results demonstrate that Arg-II promotes IL-1β production in macrophages independently of iNOS (This concept is illustrated in the **Suppl. Fig. 6G**). Future study shall further confirm the iNOS-independent stimulating effect on IL-1β production by Arg-II in the *inos^−/−^* mouse model. It remains to be investigated what are the exact molecular mechanisms of IL-1β production promoted by Arg-II and iNOS in macrophages, respectively. Since IL-1β is produced and released via activation of inflammasome (***Agostini et al., 2004; Cullen et al., 2015***), whether Arg-II and iNOS promotes macrophage pro-inflammatory responses and IL-1β release through inflammasome activation remains to be studied.

Previous studies suggest that Arg-II exerts both L-arginine metabolizing-dependent and - independent (pleiotropic) effects depending on cell types. For example, in vascular smooth muscle cells in which no NOS existing, even inactive mutant of Arg-II is capable of inducing cell senescence (***Xiong et al., 2013***), while in endothelial cells, eNOS-uncoupling, i.e., decreased NO and increased superoxide anion production, is dependent on the L-arginine metabolizing activity of Arg-II (***Yepuri et al., 2012***). It is understandable that endothelial dysfunction caused by enhanced Arg-II levels would reduce coronary perfusion and in turn decrease cardiac contractile function as reported in the literature (***Khan et al., 2012; Luo et al., 2014***). Whether Arg-II promotes inflammatory responses in macrophages through the pleiotropic effects remains to be investigated.

It is of note that a study reported that Arg-II is required for IL-10 mediated-inhibition of IL-1β in mouse BMDM upon LPS stimulation (***Dowling et al., 2021***), which suggests an anti-inflammatory function of Arg-II. The results of our study, however, demonstrate that a pro-inflammatory effect of Arg-II in macrophages. Our findings are supported by the study from another group, which shows decreased pro-inflammatory cytokine production including IL-6 and IL-1β in *arg-ii^−/−^*BMDM most likely through suppression of NFκB pathway, since *arg-ii^−/−^* BMDM reveals decreased activation of NFκB and IL-1β levels upon LPS stimulation (***Uchida et al., 2023***). Most importantly, our previous study also showed that re-introducing *arg-ii* gene back to the *arg-ii^−/−^*macrophages markedly enhances LPS-stimulated pro-inflammatory cytokine production (***Ming et al., 2012***), providing further evidence for a pro-inflammatory role of *arg-ii* under LPS stimulation. In support of this conclusion, chronic inflammatory diseases such as atherosclerosis and type 2 diabetes (***Ming et al., 2012***), inflammaging in lung (***Zhu et al., 2023***), kidney (***Huang et al., 2021***) and pancreas (***Xiong, Yepuri, Necetin, et al., 2017***) of aged animals or acute organ injury such as acute ischemic/reperfusion or cisplatin-induced renal injury are reduced in the *arg-ii^−/−^* mice (***Uchida et al., 2023***). The discrepant findings between these studies and that with IL-10 may implicate dichotomous functions of Arg-II in macrophages, depending on the experimental context or conditions. Nevertheless, our results strongly implicate a pro-inflammatory role of Arg-II in macrophages in the inflammaging in aging heart.

One of the important features of cardiac aging is fibrosis which is due to chronic inflammation process (***Lu et al., 2017***). There are convincing evidences demonstrating that chronic inflammation facilitates development of myocardial fibrosis in heart failure which is associated with aging (***Lillo et al., 2023***). A crosstalk between macrophages and fibroblasts has been demonstrated to be important for cardiac fibrosis in diseased conditions (***Hartupee et al., 2016***). Studies show that monocyte-derived macrophages from bone marrow and spleen are recruited in advanced age, contributing to cardiac fibrosis and myocardial dysfunction (***Hulsmans et al., 2018***). In our present study using an in vitro cellular model, we show a pivotal role of Arg-II-expressing macrophages in activation of cardiac fibroblasts in aging heart. First, Arg-II protein is present in the cardiac macrophages and enhanced in monocyte-derived macrophages in aging mice; second, conditioned medium from aged *arg-ii^−/−^* macrophages exhibits decreased potential of stimulating cardiac fibroblasts to produce components that are essentially involved in fibrosis such as hydroxyproline, *fibronectin, coll-IIIa,* and *tgfb1.* Interestingly, the enhanced hydroxyproline levels stimulated by old *wt* mouse macrophage-derived conditioned medium is inhibited by the IL-1β receptor blocker, demonstrating a paracrine effect of IL-1β release from aged macrophages on cardiac fibroblast activation. Moreover, the age-associated cardiac fibrosis and increase in fibroblast number in the heart is prevented in *arg-ii^−/−^* mice, demonstrating a crosstalk between macrophage and fibroblasts in cardiac aging.

The fact that cardiac fibroblasts in aging heart express elevated Arg-II, indicates a contribution of a cell-autonomous effect of Arg-II in this cell type to cardiac fibrosis in aging. This concept is confirmed by the experiments showing that overexpression of *arg-ii* in cardiac fibroblasts enhances vimentin levels associated with increased expression of *col-IIIa* and mitochondrial ROS generation. Importantly, inhibition of mitochondrial ROS by TEMPO abolished the effect of *arg-ii* overexpression in the fibroblasts, demonstrating that Arg-II exerts its cell-autonomous stimulating effects in fibroblasts through mitochondrial ROS. The role of Arg-II in mitochondrial ROS generation is further confirmed by the results showing that silencing *arg-ii* abolished miROS generation in fibroblasts when Arg-II is upregulated under hypoxic condition. These findings are supported by the studies showing that ROS favors cardiac fibroblast proliferation and activation to produce collagen (***Alili et al., 2014; Ohtsu et al., 2005***). With these results, we demonstrate that cardiac fibroblasts are activated in aging by a paracrine effect of macrophages, that is mediated by Arg-II / IL-1β and also by a cell-autonomous effect of Arg-II through mtROS.

Endothelial cells play a key role in regulation of blood flow to organs through eNOS pathway, one of the most important regulatory mechanisms in cardiovascular system (***Godo et al., 2017***). Previous studies demonstrated a causal role of Arg-II in promotion of vascular endothelial aging through eNOS uncoupling (***Yepuri et al., 2012***). In line with this, our current study showed that Arg-II is expressed in the coronary vascular endothelium of old mice and also in human heart. Emerging evidence also suggests that endothelial cells have a significant capacity for plasticity and can change their phenotype, including endothelial-to-mesenchymal transition (EndMT), a process that endothelial cells undergoing extensive alterations, including changes in gene and protein expression, loss of the endothelial phenotype, and switch to mesenchymal phenotype such as increased cell migration, proliferation, and increased collagen production, contributing to cardiovascular disease and fibrosis (***Murdoch et al., 2014; Zeisberg et al., 2007; Zhang et al., 2021***). Although EndMT in cardiac fibrotic models, particularly in pro-atherosclerotic mouse models are at least partially evidenced, it is less known in cardiovascular aging (***Evrard et al., 2016; Kidder et al., 2023; Xu et al., 2023***). When compared to young animals, aged *wt* heart tissues show enhanced levels of SNAIL, the master regulator of End-MT process. Moreover, immunofluorescence reveals that CD31^+^ endothelial cells also co-express vimentin (mesenchymal marker). No changes of endothelial marker expression such as CD31 and vascular-endothelial cadherin (data not shown) are appreciated, suggesting that partial End-MT occurs during aging. Ablation of *arg-ii* blunts the age-related elevation of both vimentin and SNAIL, thereby reducing the number of CD31^+^/vimentin^+^ endothelial cells. This data hints that lower cardiac tissue fibrosis in *arg-ii^−/−^* is at least partially contributed by dampened End-MT process. Indeed, conditioned medium from old *wt* macrophages is capable of enhancing mesenchymal markers N-cadherin and vimentin, as well as Arg-II protein levels in endothelial cells without changes in endothelial markers such as VE-cadherin, confirming a partial End-MT induced by aged macrophages. Interestingly, this effect of macrophages is prevented when *arg-ii* was ablated. Furthermore, the paracrine effects of the old macrophages on EndMT are prevented by IL-1 receptor antagonist ILRa, demonstrating a role of IL1β from old *wt* macrophages expressing Arg-II in EndMT process. Moreover, Arg-II is expressed and upregulated in endothelial cells with aging (***Yepuri et al., 2012***). Overexpression of *arg-ii* in the endothelial cells did not affect the End-MT markers (data not shown), suggesting that Arg-II alone is not sufficient to induce a cell-autonomous effect in End-MT. Finally, we demonstrate that *arg-ii^−/−^* mice in aging exhibit improved cardiac function and are more resistant to ischemic/reperfusion stress and reveal a better recovery and less myocardial infarct area as compared to the old *wt* animals. This protection against age-related cardiac dysfunction and vulnerability to ischemic stress in *arg-ii^−/−^* animals could be explained by the above described cardiac aging phenotype such as increase in apoptosis of cardiac cells, decrease in macrophage-mediated inflammation, endothelial dysfunction and EndMT, and fibroblast activation.

Previous studies provided sex-related difference in Arg-II levels at both mRNA (***Xiong, Yepuri, Montani, et al., 2017***) and protein levels in various organs (***Huang et al., 2021; Xiong, Yepuri, Necetin, et al., 2017***). We further confirmed the observation that in female mice, there is a greater age-associated upregulation of *arg-ii* levels in the heart, which is specifically found in non-cardiomyocytes but not in cardiomyocytes. A possible explanation for this sex-specific phenomenon may lie in differences in hormonal patterns between male and female animals throughout the aging process. Indeed, sex hormones, including estrogen and testosterone, have been shown to regulate *arg-ii* expression in various tissues. Estrogen seems to downregulate Arg-II (***Hayashi et al., 2006***). The age-related greater increase in *arg-ii* in the female mice might be due to loss or lack of estrogen in old female animals, which requires further investigation. However, testosterone has been reported to upregulate *arg-ii* expression (***Levillain et al., 2005***). This observation, however, does not seem to support the hypothesis that increased *arg-ii* expression in aged male mouse heart is related to sex hormones. Nevertheless, how age enhances Arg-II in sex-specific manner remains a topic of further investigation. Of note, previous studies showed that sex-related difference in pathological phenotypes in the aging kidney, pancreas, and lung are associated with higher levels of Arg-II in the females (***Huang et al., 2021; Xiong, Yepuri, Necetin, et al., 2017; Zhu et al., 2023***). The fact that aged females have higher Arg-II but are more resistant to I/R injury seems contradictory to the detrimental effect of Arg-II in I/R injury. It is presumable that cardiac vulnerability to injuries stressors depends on multiple factors/mechanisms in aging. Other factors/mechanisms associated with sex may prevail and determine the higher sensitivity of male heart to I/R injury, which requires further investigation. Nevertheless, the results of our study show that Arg-II plays a role in cardiac I/R injury also in males.

In conclusion, our study demonstrates that age-related upregulation of Arg-II in macrophage, fibroblasts, and endothelial cells contributes to the cardiac aging phenotype. This includes manifestations, such as cardiac inflammatory response, partial EndMT, fibroblast activation, resulting in cardiomyocyte apoptosis, cardiac fibrosis, and ultimately, myocardial dysfunction. Moreover, it enhances the vulnerability of aging heart to ischemic/reperfusion injury, primarily through a non-cell-autonomous paracrine release of IL-1β from macrophages acting on cardiomyocytes and endothelial cells. This effect is in addition to a cell-autonomous effect of Arg-II in cardiac fibroblasts (**Fig. 11**). Future research shall develop the macrophage-specific *arg-ii*^−/−^ mouse model to confirm this conclusion with aging animals. Since Arg-II is also expressed in fibroblasts and endothelial cells and exerts cell-autonomous and paracrine functions, aging mouse models with conditional *arg-ii* knockout in the specific cell types would be the next step to elucidate cell-specific function of Arg-II in cardiac aging. In this context, another interesting aspect is the cross-talk between macrophages and vascular SMC in the aging heart. In our present study, we could not detect Arg-II in vascular SMC of mouse heart but in that of human heart. This could be due to the difference in species-specific Arg-II expression in the heart or related to the disease conditions in human heart which is harvested from patients with cardiovascular diseases. Indeed, in the *apoe^−/−^* mouse atherosclerosis model, aortic SMCs do express Arg-II (***Xiong et al., 2013***). It is interesting to note that rodents hardly develop atherosclerosis as compared to humans. Whether this could be partly contributed by the different expression of Arg-II in vascular SMC between rodents and humans requires further investigation. In our present study, the aspect of the cross-talk between macrophages and vascular SMC is not studied. Since the crosstalk between macrophages and vascular SMC has been implicated in the context of atherogenesis as reviewed (***Gong et al., 2025***), further work shall investigate whether Arg-II expressing macrophages could interact with vascular SMC in the coronary arteries in the heart and contribute to the development of coronary artery disease and/or vascular remodelling and the underlying mechanisms. Nevertheless, our study strongly suggests that targeting Arg-II has a wide spectrum of direct and indirect effects on multiple cell types in aging heart including macrophages, fibroblasts, and endothelial cells, and cardiomyocytes, resulting in improved cardiac resistance to ischemia/reperfusion injury, which is due to inhibition of inflammation, fibrosis, and cell apoptosis. Thus, Arg-II shall represent a promising therapeutic target in treatment of age-related cardiac dysfunction and protect heart under diseased conditions.

**Fig. 11.**
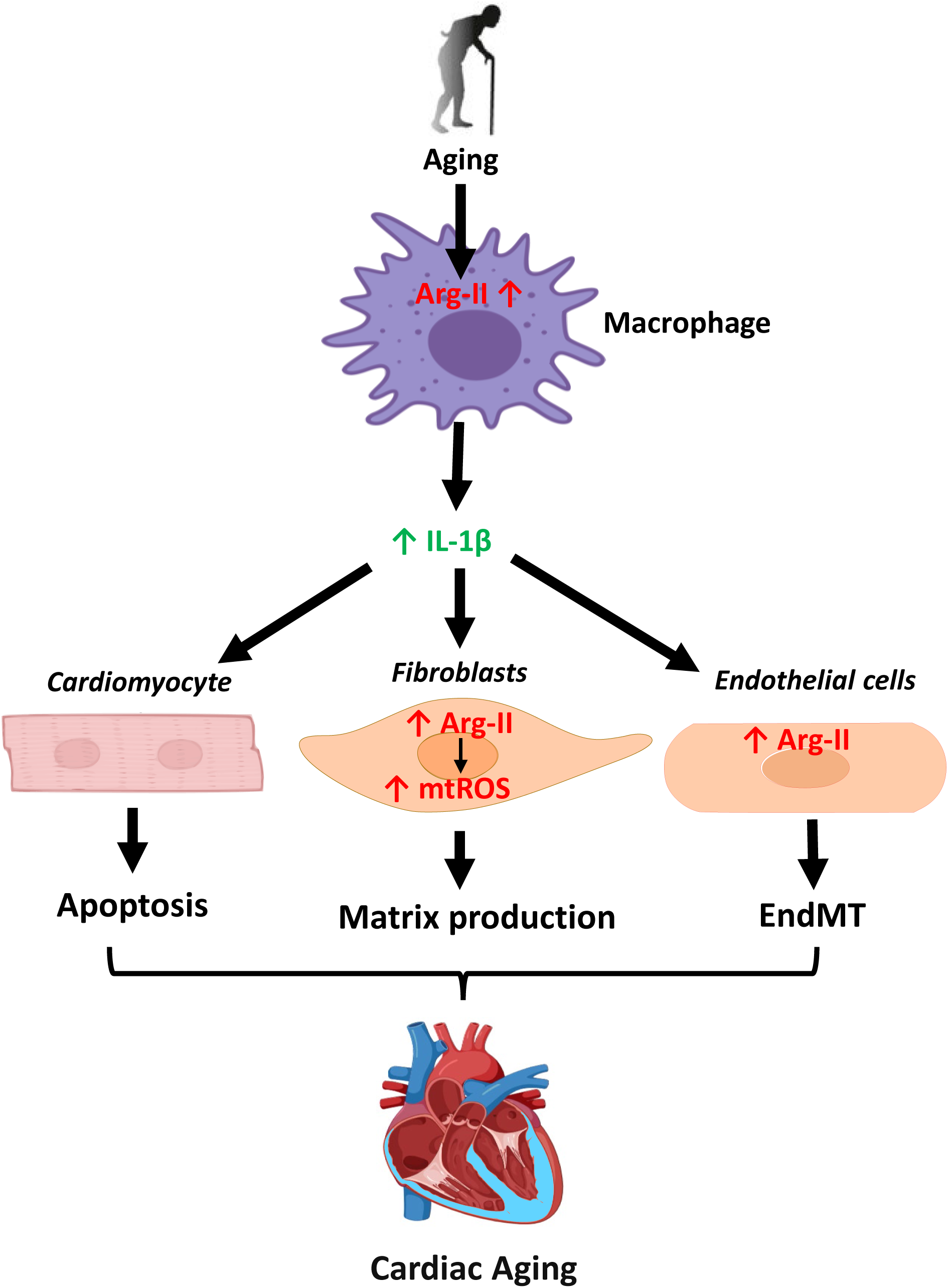
Schematic illustration of the role of Arg-II and underlying mechanisms in cardiac aging.

## Materials and methods

### Arg-II^−/−^ mouse and sample preparation

Wild type (*wt*) and *arg-ii* knockout (*arg-ii^−/−^*) mice were kindly provided by Dr. William O’Brien (***Shi et al., 2001***) and back crossed to C57BL/6 J for more than 10 generations. Genotypes of mice were confirmed by polymerase chain reaction (PCR) as previously described (***Ming et al., 2012***). Offspring of *wt* and *arg-ii^−/−^*mice were generated by interbred from hetero/hetero cross. Mice were housed at 23°C with a 12-h-light-dark cycle. Animals were fed a normal chow diet and had free access to water and food. The animals were selected and included in the study according to age, gender, and genotype. Only healthy animals, based on scoring of activity, posture, general appearance (coat, skin), locomotion, eyes/nose, body weight loss, were used. In this study, confounders such as the animal/cage location were controlled. All experimenters were aware of the group allocation at the different stages of the experiments. Male and female mice at the age of 3–4 months (young) or 20–22 months (old) were euthanized under deep anesthesia (i.p. injection of a mixture of ketamine/xylazine 50 mg/kg and 5 mg/kg, respectively) and death was confirmed by absence of all the reflexes and by exsanguination. The heart was then isolated and either snap frozen in liquid nitrogen and kept at −80 °C until use or fixed with 4% paraformaldehyde (pH 7.0), and then embedded in paraffin for immunofluorescence staining experiments. Experimental work with animals was approved by the Ethical Committee of the Veterinary Office of Fribourg Switzerland (2020-01-FR) and performed in compliance with guidelines on animal experimentation at our institution and in accordance with the updated ARRIVE guidelines (***Percie du Sert et al., 2020***). Experiments with human heart biopsies were approved by the Ethics Committee of Northwestern and Central Switzerland (Project-ID 2021-01445). Rat heart tissues were obtained from previous study (***Borrego et al., 2022***) approved by the Ethical Committee of the Veterinary Office of Fribourg Switzerland (2019-21-FR).

### Langendorff-perfused mouse heart preparation

Ex vivo functional assessments were performed by using an isolated heart system (Langendorff apparatus – EMKA technologies, France). Briefly, after euthanasia as above described, the hearts of *wt* and *arg-ii^−/−^* male and female mice (20–22 months old) were excised, cannulated and retrogradely perfused on the Langendorff system. 150 µL of heparin (2,000 units/ml) was injected 15 minutes before the heart removal to prevent blood coagulation. Hearts trimmed of extracardiac tissue were weighed immediately after surgical removal prior cannulation and heart weight to body weight ratio (HW/BW) was calculated as hypertrophy index. The aorta was cannulated on a shortened and blunted 21-gauge needle and perfusion initiated at a constant pressure of 80 mmHg. The hearts were perfused with a modified Krebs–Henseleit solution composed of (mmol/L): 118 NaCL, 4.7 KCl, 1.2 MgSO_4_, 1.2 KH_2_PO_4_, 25 NaHCO_3_, 11 Glucose, 2 CaCl_2_. The fluid was bubbled with a mix of 95% O_2_ and 5% CO_2_ at 37°C to give a pH of 7.4 and was filtered through an in-line 0.45 μm Sterivex-HV filter before delivery to the heart. A fluid-filled balloon was introduced into the left ventricle through an incision in the left atrial appendage. The ventricular balloon was connected via fluid-filled tubing to a pressure transducer for continuous assessment of ventricular function. The balloon was inflated to yield a left ventricular end-diastolic pressure between 5 to 15 mmHg. Hearts were immersed in warmed perfusate in a jacketed bath maintained at 37°C, and perfusate delivered to the coronary circulation was maintained at the same temperature. After 20 minutes stabilization, hearts were subjected to 15 minutes (males), 20 minutes and 30 minutes (females) of no-flow ischemia followed by 30-45 minutes of reperfusion. Recovery of contraction and relaxation rates (dt/dp_min_) as well as the recovery of the left ventricle developed pressure (LV-DP) and left ventricular end diastolic pressure (LV-EDP) were measured to investigate tolerance to ischemia/reperfusion (I/R)-injury. In addition to functional recovery, triphenyl tetrazolium chloride (TTC) staining was also performed to investigate a potential reduction in infarct size upon I/R-injury in *arg-ii^−/−^* animals. TTC is a colorless water-soluble dye that is reduced by the mitochondrial enzyme succinate dehydrogenase of living cells into a water-insoluble, light sensitive compound (formazan) that turns healthy/normal tissue deep red. In contrast, damaged/dead tissue remains white showing the absence of living cells, and thereby indicating the infarct region (***Bederson et al., 1986***). After completing the I/R protocol, heart was carefully removed from the experimental set-up. After a freeze-thaw cycle (−20°C for 1 hour), the heart was cross-sectioned from the apex to the atrioventricular groove into four 2.5 mm thick slices. The slices were first incubated in TTC (20 minutes at 37°C), and then fixed in 10% formalin for 30 minutes. A cover glass was then placed over the tissues and images were acquired (***Redfors et al., 2012***). ImageJ software was used to analyze the images and assess the infarcted area.

### Real-time quantitative RT-PCR

mRNA expression level of genes from mouse and human origin was measured by two-step quantitative Real Time-PCR as described previously (***Ming et al., 2012***). Ribosomal Protein S12 (*rps12*), and glyceraldehyde-3-phosphate dehydrogenase (*gapdh*) were used as reference genes. Total RNA from mouse tissue, splenic macrophages, and human cardiac fibroblasts was extracted with Trizol Reagent (TR-118, Molecular Research Center) following the manufacturer’s protocol. Real-time PCR reaction was performed with the GOTaq® qPCR Master Mix (A6001, Promega) and CFX96 Real-Time PCR Detection System (Bio-Rad). The mRNA expression level of all genes was normalized to the reference gene *rps12* or *gapdh*. All the qRT-PCR primer sequences are shown in the **Suppl. Table 2.**

### Confocal immunofluorescence microscope

Immunofluorescence staining was performed with heart tissues. Briefly, mouse hearts or human cardiac biopsies were fixed with 4% paraformaldehyde (pH 7.0) and embedded in paraffin. After deparaffinization in xylene (2 times of 10 minutes), the sections were treated in ethanol (twice in 100% ethanol for 3 minutes and twice in 95% ethanol and once in 80%, 75%, 50% ethanol for 1 minute sequentially) followed by antigen retrieval (EDTA buffer, pH 8.0) for Arg-II, troponin-T (TropT), CD31, p16, Mac-2, IL-1β, vimentin (Vim), CD-68, PDGF-Rα and α-smooth muscle actin (αSMA) in a pressure cooker. For co-immunofluorescence staining of Arg-II/TropT, Arg-II/CD31, Arg-II/Vim, Arg-II/CD-68, Arg-II/Mac-2, Arg-II/PDGF-Rα, Arg-II/αSMA, IL-1β/Mac-2, F4-80/CCR2, F4-80/LYVE1 (Suppl. Fig. 10 shows the validation of CCR2 and LYVE1 antibodies) and CD31/Vim, primary antibodies of different species were used. Transverse sections (5 μm) were blocked with mouse Ig blocking reagent (M.O.M, Vector laboratories) for 2 hours and then with PBS containing 1% BSA and 10% goat serum for 1 hour. For CD31 and PDGF-Rα (both polyclonal Goat IgGs), a blocking reagent containing 10% BSA was used. The sections were then incubated overnight at 4°C in a dark/humidified chamber with target primary antibodies, and subsequently incubated for 2 hours with the following secondary antibodies: Alexa Fluor 488–conjugated goat anti-rabbit IgG (H + L) and Alexa Fluor 568-conjugated goat anti-mouse IgG (H + L); Alexa Fluor 488–conjugated goat anti-mouse IgG (H + L) and Alexa Fluor 594-conjugated goat anti-rabbit IgG (H + L); Alexa Fluor 488–conjugated donkey anti-goat IgG (H + L) and Alexa Fluor 594-conjugated goat anti-rabbit IgG (H + L). All sections were finally counterstained with 300 nmol/L DAPI for 5 minutes. Immunofluorescence signals were visualized under Leica TCS SP5 confocal laser microscope. The antibodies used are shown in **Suppl. Table 3.**

### Isolation and cultivation of ventricular cardiomyocytes

Ventricular myocytes were isolated from five-month old female mice according to an established protocol (***Louch et al., 2011***). Briefly, the hearts were excised, cannulated and retrogradely perfused on a Langendorff system as described above. 150 µl of heparin (2,000 units/ml) was injected 15 minutes before the heart removal to prevent blood coagulation. To isolate the cardiomyocytes, hearts were perfused at 37°C for 20 minutes with a Ca^2+^-free solution composed of (in mmol/ liter): 130 NaCl, 5 KCL, 0.5 NaH_2_PO_4_, 10 HEPES, 10 Glucose, 10 2,3 butanedione monoxime (BDM), 10 Taurine, 1 MgCl_2_ (pH 7.4, adjusted with NaOH). Cells were enzymatically dissociated using a cocktail of collagenase type II (210 U/ml, Worthington), collagenase type IV (260 U/ml, Worthington) and protease type XIV (0.21 U/ml; Sigma-Aldrich). After isolation, cardiomyocytes were separated by gravity settling and Ca^2+^ was restored to a final concentration of 1 mmol/L. The cardiomyocytes were finally plated onto laminin-coated coverslips (5 μg/ml, Thermo Scientific) and cultured in the M-199 medium supplemented with bovine serum albumin (BSA; 0.1 % - A1470, Sigma-Aldrich), BDM (10 mmol/L), insulin, transferrin, selenium (ITS; I3146 - Sigma-Aldrich), CD lipid (11905-031 - Sigma-Aldrich) and penicillin / streptomycin mix (1 %).

### Cell culture, adenoviral transduction, and inos^−/−^ and arg-ii^−/−^ macrophages

Human cardiac fibroblasts (HCF) from adult human heart tissue were purchased from Innoprot (P10452) and cultured in the proprietary fibroblast culture medium (P60108-2, Innoprot) composed as follow: 500 ml of fibroblast basal medium, 25 ml of FBS, 5 ml of fibroblast growth supplement-2, and 5 ml of penicillin/streptomycin in the Poly-L-Lysine (PLL) coated flasks and dishes. The cells were maintained at 37°C in a humidified incubator (5% CO_2_ and 95% atmosphere air). To overexpress *arg-ii*, the cells were seeded on 6-cm dishes for 24 hours and transduced with rAd-CMV-Con/*arg-ii* for 48 hours as described previously (***Ming et al., 2012***).

Human umbilical vein endothelial cells (HUVEC) cells were cultured in RPMI-1640 (with 25 mmol/L HEPES and stable glutamine; Thermo Fisher Scientific) containing 5 % fetal calf serum (FCS; Gibco) and 1% streptomycin and penicillin in gelatin (1%) coated dishes.

Raw 264.7 cells (mouse macrophage cell line) were cultured in Dulbecco modified Eagle medium/F12 (DMEM/F12; Thermo Fisher Scientific) containing 10% fetal bovine serum (FBS; Gibco) and 1% streptomycin and penicillin. To silence *arg-ii*, the cells were seeded on 6-cm dishes for 24 hours and transduced first with the rAd at titers of 100 Multiplicity of Infection (MOI) and cultured in the complete medium for 2 days and then in serum-free medium for another 24 hours before experiments. The rAd expressing shRNA targeting *arg-ii* driven by the U6 promoter (rAd/U6-*arg-ii*-shRNA) and control rAd expressing shRNA targeting *lacz* (rAd/U6-*lacz*-shRNA) were generated as described previously (***Ming et al., 2009***).

*Wt* and *inos* gene deficient bone-marrow-derived macrophage cell lines (MØ*^wt^* and MØ*^inos-/-^*) were purchased from Kerafast (ENH166-FP and ENH176-FP) and cultured in Dulbecco modified Eagle medium (DMEM; Thermo Fisher Scientific) containing 10% fetal bovine serum (FBS; Gibco) and 1% streptomycin and penicillin. The cells were treated with LPS (100 ng/mL) for 24 hours before harvest. Gene deficiency of *inos* was verified by immunoblotting analysis.

Knockout of *arg-ii* gene in human THP-1 cells was generated by CRISPR-UTM-mediated genome engineering with gRNA targeting exon 1 (Guangzhou Ubigene Biosciences Co., Ltd.). The control THP1 cell line was generated with scramble gRNA and Cas9. Gene deficiency of *arg-ii* was verified by immunoblotting analysis.

### TUNEL Staining

Mouse hearts were isolated and fixed with 4% paraformaldehyde (pH 7.0), embedded in paraffin and sliced at 5 μm of thickness. After deparaffinization (xylene, 2 times for 10 minutes) and rehydration (sequential washing in 100%, 95%, 80%, 75% and 50% ethanol), TUNEL staining was performed using a commercial kit (in situ Cell Death Detection Kit, TMR red – Roche, Switzerland). Briefly, 50 μl of TUNEL mixture, containing TdT and dUTP in reaction buffer were added onto the sections and incubated in a humidified chamber for 60 minutes at 37°C in darkness. Positive controls were generated by incubating samples at room temperature (RT) for 10 minutes with DNase in PBS (1 mg/ml) to induce strand breaks. Negative controls were generated by incubation with TUNEL mixture lacking TdT. DAPI counterstaining was finally performed (300 nmol/l for 5 minutes).

TUNEL staining was also performed on isolated ventricular cardiomyocytes. After tissue digestion, the cardiomyocytes were plated onto murine laminin-coated coverslips to promote cell adhesion. One hour after plating, several PBS washing steps were performed to remove damaged and partially adherent cells, ensuring that only well-shaped, viable, and strongly adherent cells remained as bioassay cells. These “healthy” cells were then selected for the experiments. After washing, cardiomyocytes were incubated for 24 hours with either splenic macrophage conditioned media (CM; see below) or with RAW cells CM, and the following protocol was applied: cells are fixed using 4% paraformaldehyde for 1 hour at RT, and then permeabilized via Triton X (0.1 %). As for tissues, 50 μl of TUNEL mixture, containing TdT and dUTP in reaction buffer were added onto each slide, and then incubated at 37°C for 60 minutes in darkness. DAPI counterstaining was then performed.

Upon plating onto murine laminin-coated coverslips, the cardiomyocytes were first incubated for 24 hours with either splenic macrophage conditioned media (CM; see below) or with RAW cells CM, and the following protocol was applied: cells are fixed using 4% paraformaldehyde for 1 hour at RT, and then permeabilized via Triton X (0.1 %). As for tissues, 50 μl of TUNEL mixture, containing TdT and dUTP in reaction buffer were added onto each slide, and then incubated at 37°C for 60 minutes in darkness. DAPI counterstaining was then performed.

For experiments aiming to define cell borders, TUNEL-protocol was followed by a staining with Wheat Germ Agglutinin (WGA)-Alexa Fluor 488 (10 μg/ml for 30 minutes). For heart tissue, five representative images of each section are taken using a Leica TCS SP5 confocal system, and WGA staining was performed to differentiate cardiomyocytes and non-cardiomyocytes according to cell size and presence or absence of striations (characteristics of cardiomyocytes). Data was reported as number of TUNEL-positive nuclei over the total number of nuclei in the images.

### Masson’s trichrome staining

To analyze cardiac fibrosis, heart sections (5 µm) were subjected to Masson’s trichrome (ab150686, Abcam) staining according to the manufacturer’s instructions (***de Guzman et al., 2023***).

### Hydroxyproline colorimetric assay

Collagen content was analyzed by determination of hydroxyproline levels in left ventricles or in cell homogenates using the Hydroxyproline Assay kit (MAK008, Sigma) according to the manufacturer’s instructions.

### Isolation of splenic macrophages

Splenic cells were isolated as previously described (***Wang et al., 2013***). Briefly, young and old *wt* and *arg-ii^−/−^* female mice were euthanized as described and spleens were dissected from abdominal cavity and filtered through a 70-μm nylon strainer. Red blood cell lysis buffer was used to remove red blood cells. The cell suspension obtained was plated onto gelatin-coated dishes at a density of 2 × 10^6^ cells/ml. After 3 days, cells were washed to ensure macrophage enrichment. Following washing, the macrophages were cultured for further 4 days in RPMI culture media with 20% of L929 cell line conditioned medium (source of the macrophage growth factor M-CSF) to allow cell maturation. Medium was changed every two days.

### Crosstalk between macrophages and heart cells

For crosstalk experiments of splenic macrophages from *wt* and *arg-ii^−/−^* mice with cardiac cells, conditioned medium from the isolated splenic macrophages was collected and transferred to cardiomyocytes, fibroblasts and endothelial cells in 1:1 dilution with the specific cell culture medium for the indicated time. To study effects of IL-1β in the conditioned medium of splenic macrophages (CM-SM), cardiac cells, fibroblasts or endothelial cells were pre-treated with IL-1 receptor antagonist IL-1ra (50 ng/ml) for 2 hours followed by the specific experimental protocols for analysis of cardiac apoptosis, fibroblast activation, and EndMT transition, respectively.

For the experiments with mouse macrophage cell line Raw 264.7, the cells were seeded on six-well plates with a density of 2 × 10^5^ cells per well. Upon transduction with rAd expressing *arg-ii^shRNA^* (see above) and over-night serum starvation, the mouse macrophages were polarized toward a pro-inflammatory secretion profile by stimulation with lipopolysaccharide (LPS; 100 ng/mL) for 24 hours. The conditioned medium (CM) from non-treated control and LPS-activated RAW cells (referred to as CM-RAW) was then filtered and transferred onto cardiomyocytes for 24 hours to investigate cell apoptosis. To study effects of IL-1β in CM-RAW on cardiac cells apoptosis, myocytes were pre-treated with IL-1 receptor antagonist IL-1ra (280-RA, R&D systems) for 2 hours.

### Hypoxia experiments with murine cardiac fibroblasts and cardiomyocytes

Cardiac cells were isolated from *wt* mice according to an established protocol (***Louch et al., 2011***). Briefly, after euthanasia of mice, hearts were excised, cannulated and retrogradely perfused on a Langendorff system. Cells were enzymatically dissociated using a cocktail of collagenase type II, collagenase type IV and protease type XIV. After isolation, cardiomyocytes were separated from the pull of cardiac cells by gravity settling and plated onto laminin-coated coverslips. After 2 hours, the cardiomyocytes were washed to remove non-adhering cells and incubated for 24 hours at 1% oxygen level in a Coy In Vitro Hypoxic Cabinet System (The Coy Laboratory Products, Grass Lake, MI USA). The rest of cardiac cells were plated on 12-well plates and cultured in DMEM (10% FBS) for 3 days, allowing cardiac fibroblasts to adhere. The fibroblasts were passaged once and subjected to hypoxia as for the cardiomyocytes.

### Hypoxia experiments with human cardiac fibroblasts and mitochondrial superoxide detection (MitoSOX staining)

Human cardiac fibroblasts (HCF) from adult human heart tissue were incubated for 48 hours at 1% oxygen level in a Coy In Vitro Hypoxic Cabinet System (The Coy Laboratory Products, Grass Lake, MI USA). To silence *arg-ii*, HCFs were seeded on 6-cm dishes for 24 hours and transduced first with the rAd at titers of 100 Multiplicity of Infection (MOI) and cultured in the complete medium for 2 days and then in serum-free medium for another 24 hours before experiments. The rAd expressing shRNA targeting *arg-ii* driven by the U6 promoter (rAd/U6-arg-ii-shRNA) and control rAd expressing shRNA targeting lacz (rAd/U6-lacz-shRNA) were generated as described previously (Ming et al., 2009). Mitochondrial superoxide generation was monitored by using MitoSOX. Briefly, the cells were incubated with MitoSOX at the concentration of 1 μmol/L for 30 minutes. After washing, the cells were then fixed with 3.7% paraformaldehyde followed by counterstaining with DAPI and then subjected to imaging through 40× objectives with Leica TCS SP5 confocal laser microscope. To ensure the mitochondria as source of the ROS, cells were treated with Mito TEMPO (10 μmol/L, 1 hour) followed by MitoSOX staining.

### ELISA for IL-1β detection in conditioned media

IL-1β concentrations in the conditioned medium from murine splenic cells were measured by ELISA kits (mouse IL-1β, 432604, BioLegend, San Diego, USA) according to the manufacturer’s instructions. IL-1β concentrations in the conditioned medium from THP1 human cells were measured by ELISA kits (human IL-1β, 437004, BioLegend, San Diego, USA) according to the manufacturer’s instructions.

### Immunoblotting

Heart tissue or cell lysate preparation, SDS-PAGE and immunoblotting, antibody incubation, and signal detection were performed as described previously (***Ming et al., 2012***). To prepare heart homogenates, frozen cardiac tissues were crushed into the fine powder using a mortar and pestle in liquid nitrogen on ice. A portion of the fine powder was then homogenized (XENOX-Motorhandstück MHX homogenizer) on ice in 150 µl of ice-cold lysis buffer with the following composition: 10 mmol/L Tris-HCl (pH 7.4), 0.4 % Triton X-100, 10 μg/ml leupeptin, and 0.1 mmol/l phenylmethylsulfonyl fluoride (PMSF), protease inhibitor cocktail (B14002, Bio-Tool) and phosphatase inhibitor cocktail (B15002; Bio-Tool). Homogenates were centrifuged (Sorvall Legend Micro 17R) at 13,000 × *g* for 15 minutes at 4 °C. Protein concentrations of supernatants were determined by Lowry method (500-0116, Bio-Rad). Equal amount of protein from each sample was heated at 75°C for 15 minutes in loading buffer and separated by SDS-PAGE electrophoresis. Proteins in the SDS-PAGE gel were then transferred to PVDF membranes which are blocked with PBS-Tween-20 supplemented with 5% skimmed milk. The membranes were then incubated with the corresponding primary antibody overnight at 4°C with gentle agitation. After washing with blocking buffer, the membranes were then incubated with the corresponding anti-mouse or anti-rabbit secondary antibody. Signals were visualized using the Odyssey Infrared Imaging System (LI-COR Biosciences), or the FUSION FX Imaging system (Witec AG) for chemiluminescence, and quantified by Image Studio Lite (5.2, LI-COR Biosciences). The antibodies used in this study are listed in the **Suppl. Table 3.**

### Statistical analysis

Data were presented as mean ± SD. Data distribution was determined by Kolmogorov-Smirnov test and statistical analysis for normally distributed values was performed with Student’s unpaired t-test or analysis of variance (ANOVA) with Bonferroni post hoc test. For non-normally distributed values, Mann–Whitney test or the Kruskal–Wallis test were used. Differences in mean values were considered statistically significant at a two tailed *p*-value ≤ 0.05.

## Author contributions

DMP, XC, GA, AB, M-NG, AF, and SC performed experiments, data acquisition, analysis, and interpretation, KDM provided human samples; DMP, ZY and X-FM designed the project and prepared figures and drafted the manuscript; ZY and X-FM received funding for the study; All the authors discussed the results and critically revised and approved the manuscript for important intellectual contents.

## Availability of data and materials

Data supporting the present study are available from the corresponding author upon reasonable request.

---

**Suppl. Table 1.** Baseline comparison between *wt* and *arg-ii^−/−^* mice under Langendorff recordings.

| **N = 5**<br>**Each**<br>**group** | **Pmax**<br>**(mm**<br>**Hg)** | **Pmin**<br>**(mm**<br>**Hg)** | **DP**<br>**(ms)** | **SEP**<br>**(ms)** | **DFP**<br>**(ms)** | **CT**<br>**(ms)** | **RT**<br>**(ms)** | **dPmax** | **dPmin** | **CI** | **HR**<br>**(beats/**<br>**min)** |
| --- | --- | --- | --- | --- | --- | --- | --- | --- | --- | --- | --- |
| **wtO F** | 71.8±<br>22.8 | 9.8±<br>2.7 | 65.8±<br>19.7 | 43.6±<br>15.5 | 227±<br>115 | 28.4±<br>2.15 | 68.2±<br>3.9 | 1807±<br>854 | -1133±<br>566 | 63±<br>5.4 | 214.8±<br>55.6 |
| ***arg-ii<sup>-/-</sup>***<br>**O F** | 91.2±<br>20.6 | 7.4±5.<br>2 | 85.0±<br>23.3 | 41.8±<br>6.8 | 270±<br>135 | 31.8±<br>1.2 | 67.4±<br>6.7 | 2340±<br>707 | -1759<br>±569 | 57.6±<br>2.6 | 189.6±<br>47.6 |

\* DP = developed pressure
\* SEP = systolic ejection period (ms)
\* DFP = diastolic filling time (ms)
\* CT = contraction time (ms)
\* RT = relaxation time (ms)
\* CI = contractility Index
\* HR = Heart rate

**Suppl Table 2.**
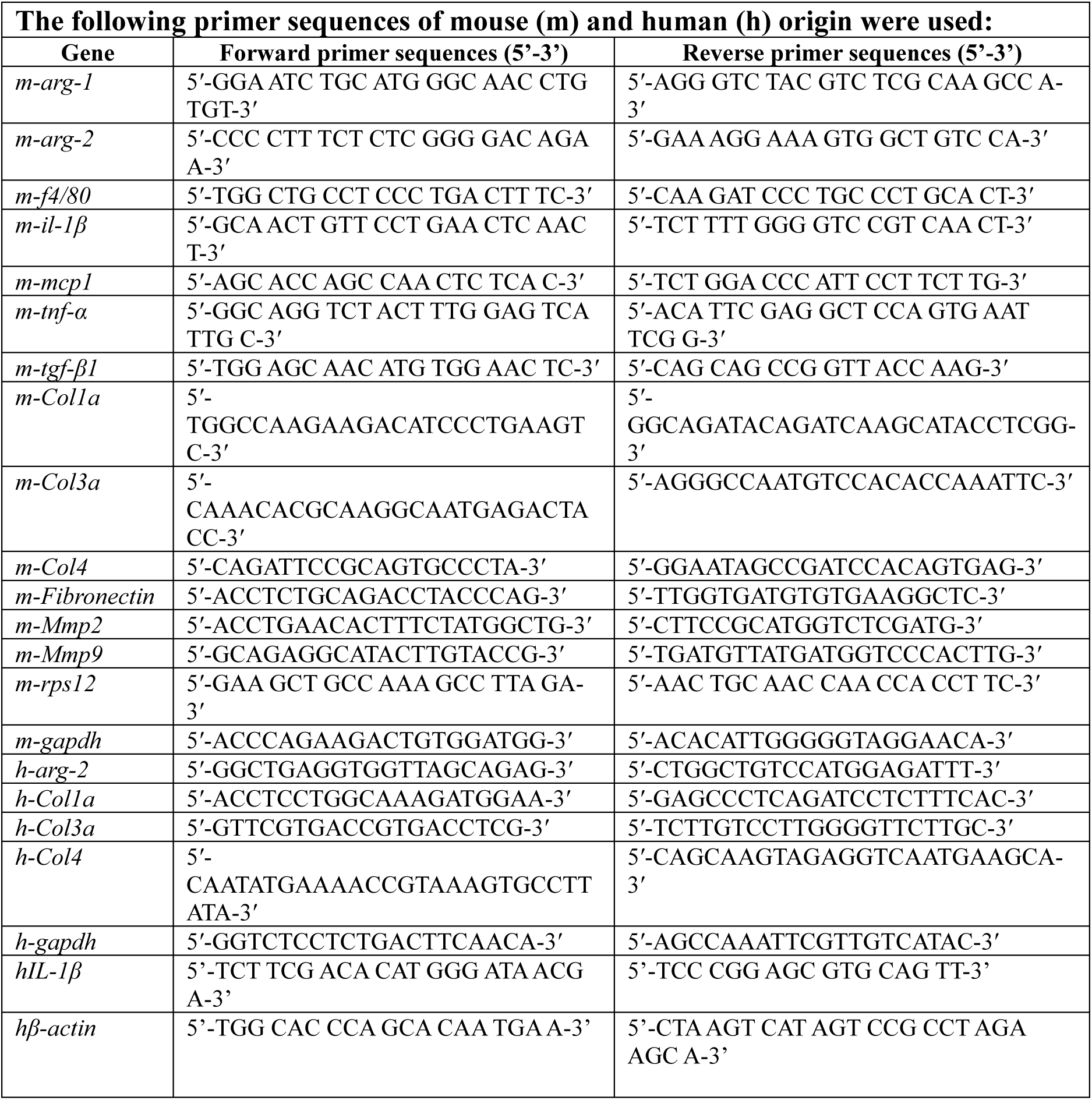
The RT-PCR primer sequences.

**Suppl. Table 3.** Antibody dilutions used for immunoblotting and immunofluorescence staining.

| **Antibody target** | **Dilution** |
| --- | --- |
| Arg-II (#55003, cell signaling) | WB 1:1000;<br>IF 1:100 |
| IL-1β (ab9722, Abcam) | WB 1:1000<br>IF 1:100 |
| Hif-1α (#36169, cell signaling) | WB 1:1000 |
| α-SMA (ab7817, Abcam) | IF 1:200 |
| α-Tubulin (T5168, Sigma) | WB 1:4000 |
| iNOS (ab178945, Abcam) | WB 1:1000 |
| Cardiac Troponin T (ab8295, Abcam) | IF 1:100 |
| Mac-2 (14-5301-82, Invitrogen) | IF 1:200 |
| CD31 (sc-1506, Santa Cruz) | WB 1:1000;<br>IF 1:100 |
| P16 (sc-81156, Santa Cruz) | IF 1:50 |
| LYVE1 (14-0443-82; ThermoFisher) | IF 1:100 |
| CCR2 (ab223050, Abcam) | IF 1:100 |
| F4-80 (30325S, cell signaling) | IF 1:100 |
| F4-80 (BM4007S, OriGene) | IF 1:100 |
| CD68 (MA5-13324KP1; ThermoFisher) | WB 1:100 |
| PDGF-Rα (AF1062, R D systems) | IF 1:200 |
| Vimentin (ab8978, Abcam) | WB 1:1000;<br>IF 1:100 |
| Vinculin (MCA465GA, Bio-Rad) | WB 1:5000 |
| VE-Cadherin (ab33168, Abcam) | WB 1:1000 |
| N-Cadherin (#13116, cell signaling) | WB 1:1000 |
| GAPDH (10R-G109A; Fitzgerald Biosciences) | WB 1:100,000 |
| Snail + SLUG (ab180714, Abcam) | WB 1:500 |
| IRDye 800-conjugated affinity purified goat anti-rabbit IgG (9263221, BioConcept) | WB 1:5,000 |
| Alexa fluor 680-conjugated goat anti-mouse IgG (A-21057, Invitrogen) | WB 1:5,000 |
| Alexa Fluor 488-conjugated goat anti-rabbit IgG (H+L) secondary Ab (A-11008, Thermo Fisher Scientific) | IF 1:400 |
| Alexa Fluor 594-conjugated goat anti-rabbit IgG (H+L) secondary Ab (A-11012, Thermo Fisher Scientific) | IF 1:400 |
| Alexa Fluor 488-conjugated goat anti-mouse IgG (H+L) secondary Ab (A-11001, Thermo Fisher Scientific) | IF 1:400 |
| Alexa Fluor 568-conjugated goat anti-mouse IgG (H+L) secondary Ab (A-11031, Thermo Fisher Scientific) | IF 1:400 |
| Alexa Fluor 488-conjugated donkey anti-goat IgG (H+L) secondary Ab (A11055, Thermo Fisher Scientific) | IF 1:400 |

**Suppl. Fig. 1.**
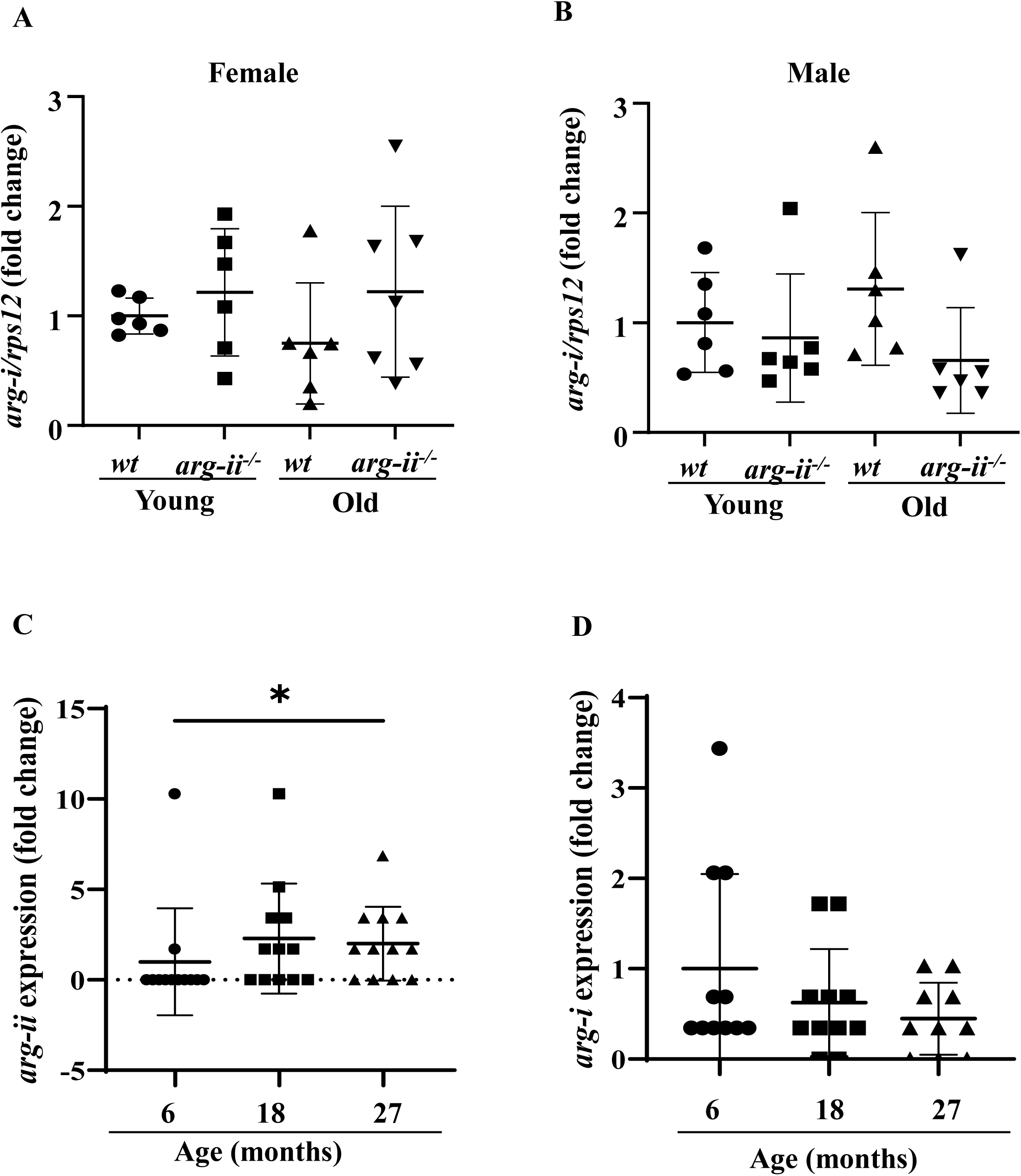
*arg-ii* and *arg-i* expression in male and female mice. *arg-i* mRNA levels of female (**A**) and male (**B**) young (3-4 months) and old (20-22 months) wild type (*wt*) heart tissues analyzed by qRT-PCR. *rps12* served as the reference (n=6 to 7 animals per group); *arg-ii* (**C**) and *arg-i* (**D**) expression extrapolated from high throughput sequencing database of male mice of 6, 18 and 27 months old (GSE201207, Wolff et al., 2023). Data are presented as the fold change to the young-*wt* group. \**p*≤0.05 between the indicated groups. *wt*, wild-type mice; *arg-ii^−/−^*, *arg-ii* gene knockout mice.

**Suppl. Fig. 2.**
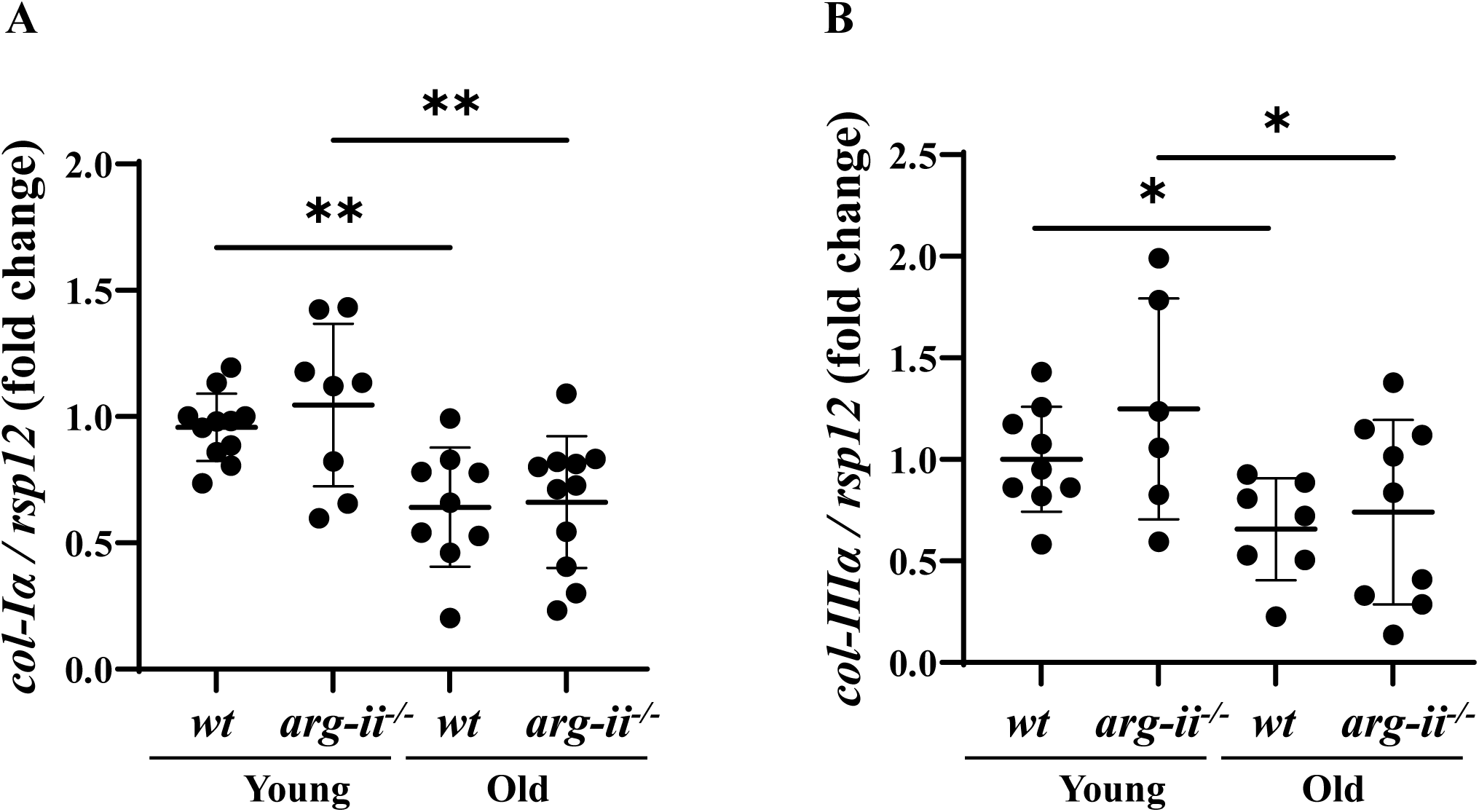
Collagen genes expression in female mice. (**A**) *Col-Iα* and (**B**) *col-IIIα* mRNA levels of young (3-4 months) and old (20-22 months) *wt* and *arg-ii*^−/−^ female heart tissues analyzed by qRT-PCR. *rps12* served as the reference (n=6 to 9 animals per group). Data are presented as the fold change to the young-*wt* group. \**p*≤0.05, \*\**p*≤0.01 between the indicated groups. *wt*, wild-type mice; *arg-ii^−/−^*, *arg-ii* gene knockout mice

**Suppl. Fig. 3.**
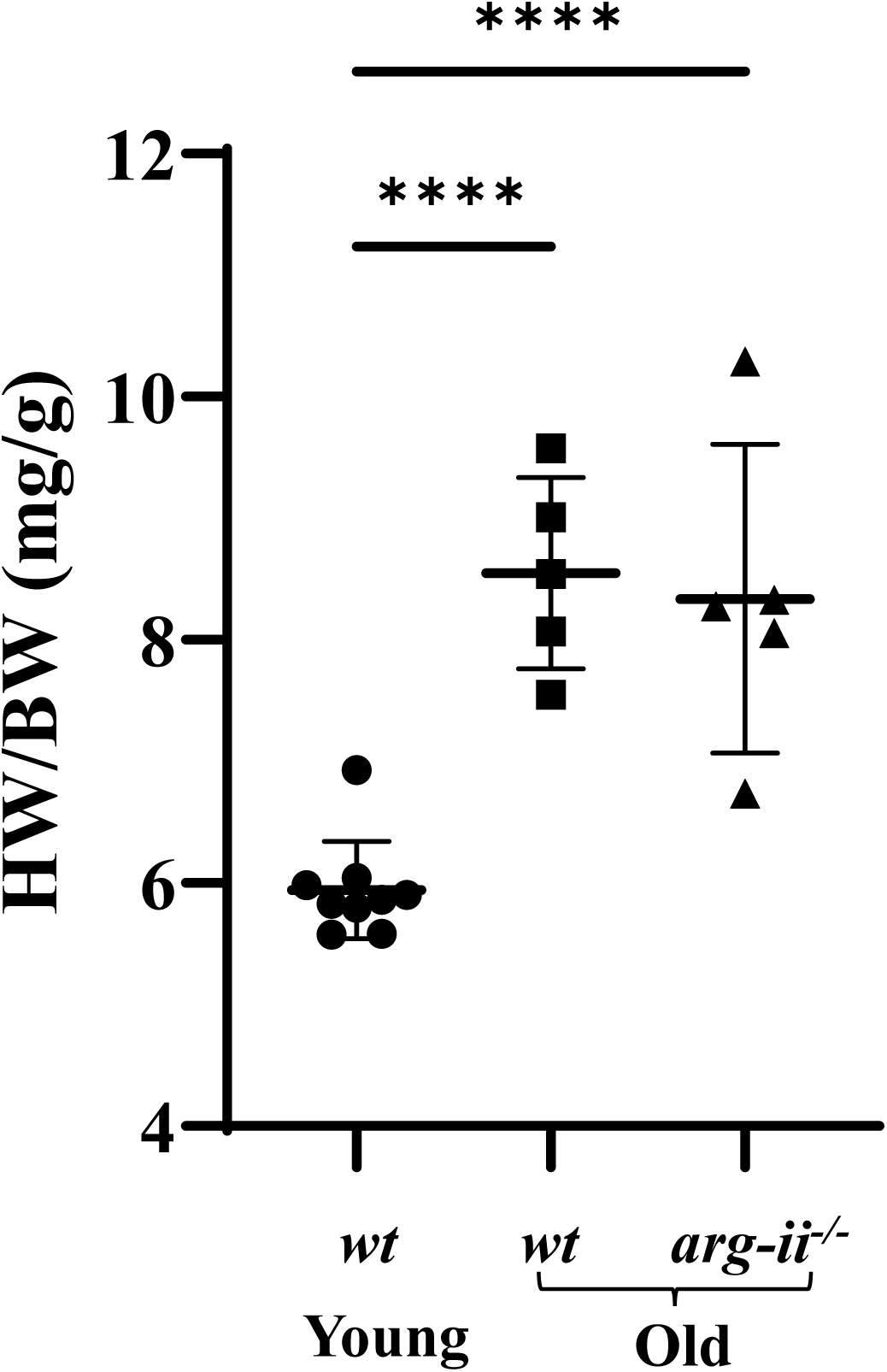
Age-related heart tissue hypertrophy. Comparison between the heart weigh to body weight ratio (HW/BW) of *wt* and *arg-ii^−/−^* young and old mice (n=5-9 per group). \*\*\*\**p* ≤ 0.001, between the indicated groups. *wt*, wild-type mice; *arg-ii^−/−^*, *arg-ii* gene knockout mice.

**Suppl. Fig. 4.**
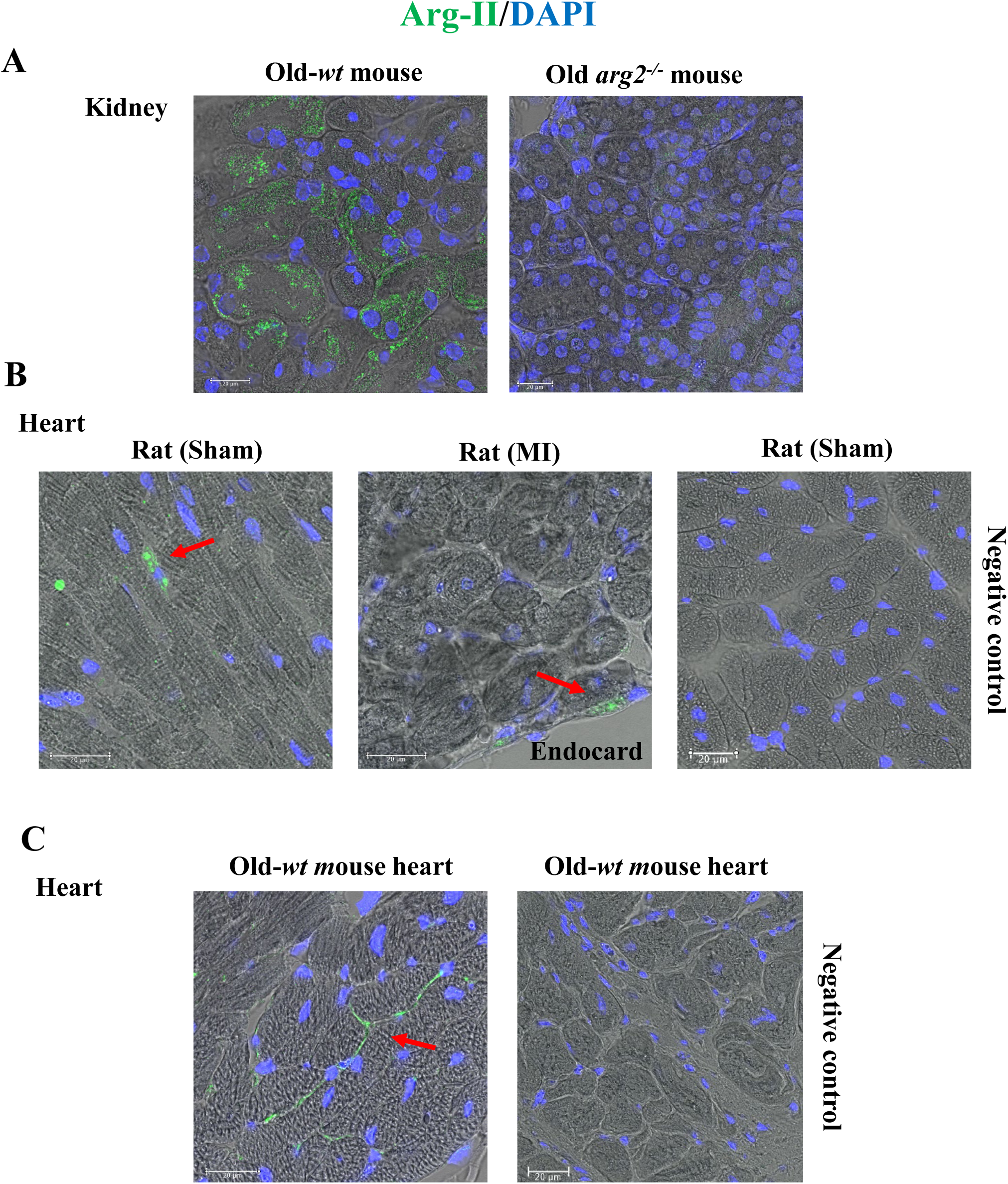
Arg-II localization in mouse and rat tissues. (**A**) Representative confocal images showing immunofluorescence staining of Arg-II (green) in old wt and *arg-ii^−/−^* kidney tissue. DAPI (blue) is used to stain nuclei. Brightfield light microscopy was combined to fluorescent signal to assess cell morphology. Scale bar = 20 µm; (**B**) Representative confocal images showing immunofluorescence staining of Arg-II (green) in heart tissue of control rat (Sham), rat under myocardial infarction followed artery ligation (MI) and relative negative control (omission of primary Ab). DAPI (blue) is used to stain nuclei. The images show Arg-II localization in non-myocytes cells. (**C**) Representative confocal images showing Arg-II (green) in old *wt* mouse tissue with relative negative control (omission of primary Ab). DAPI (blue) is used to stain nuclei.

**Suppl. Fig. 5.**
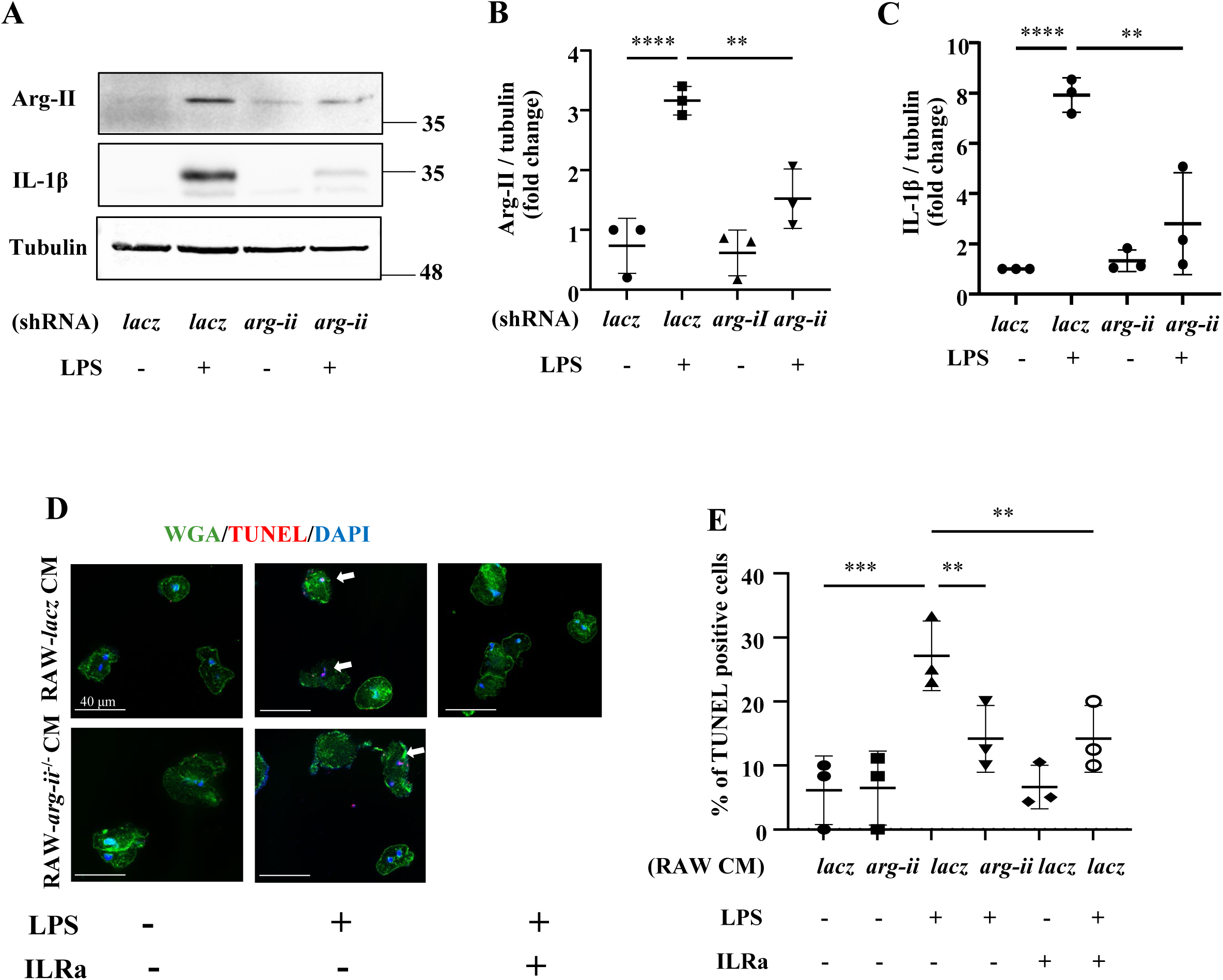
*In vitro* crosstalk between LPS-activated RAW 264.7 macrophages and cardiomyocytes. (**A**) Immunoblotting analysis of Arg-II and Il-1β (precursor) in RAW 264.7 mouse macrophages upon transduction with the rAd/U6-LacZshRNA as the control or rAd/U6-Arg-IIshRNA to silence Arg-II. RAW cells were polarized toward pro-inflammatory phenotype by incubation with lipopolysaccharide (LPS; 100 ng/mL) for 24 hours; tubulin served as protein loading control. Molecular weight (kDa) is indicated at the side of the blots. The plot graphs show the quantification of Arg-II (**B**) and Il-1β (**C**) protein signals on immunoblots (n=3); (**D**) Representative confocal images of adult isolated mouse cardiomyocytes stimulated with conditioned media (CM) from control RAW cells and RAW cells with *arg-ii* silencing, with and without LPS (24 h incubation). Wheat Germ Agglutinin (WGA)-Alexa Fluor 488-conjugate was used to stain cell membrane, and TUNEL was performed to identify apoptotic cells. DAPI is used to stain nuclei. Interleukin receptor antagonist (ILRa; 50 ng/mL) is used to prevent IL-1β binding to its receptor; (**E**) Quantification of TUNEL-positive cardiomyocytes (% in respect to total number of cells). Scale bar = 40 µm. \*\**p*≤0.01, \*\*\**p*≤0.005 and \*\*\*\**p*≤0.001 between the indicated groups.

**Suppl. Fig. 6.**
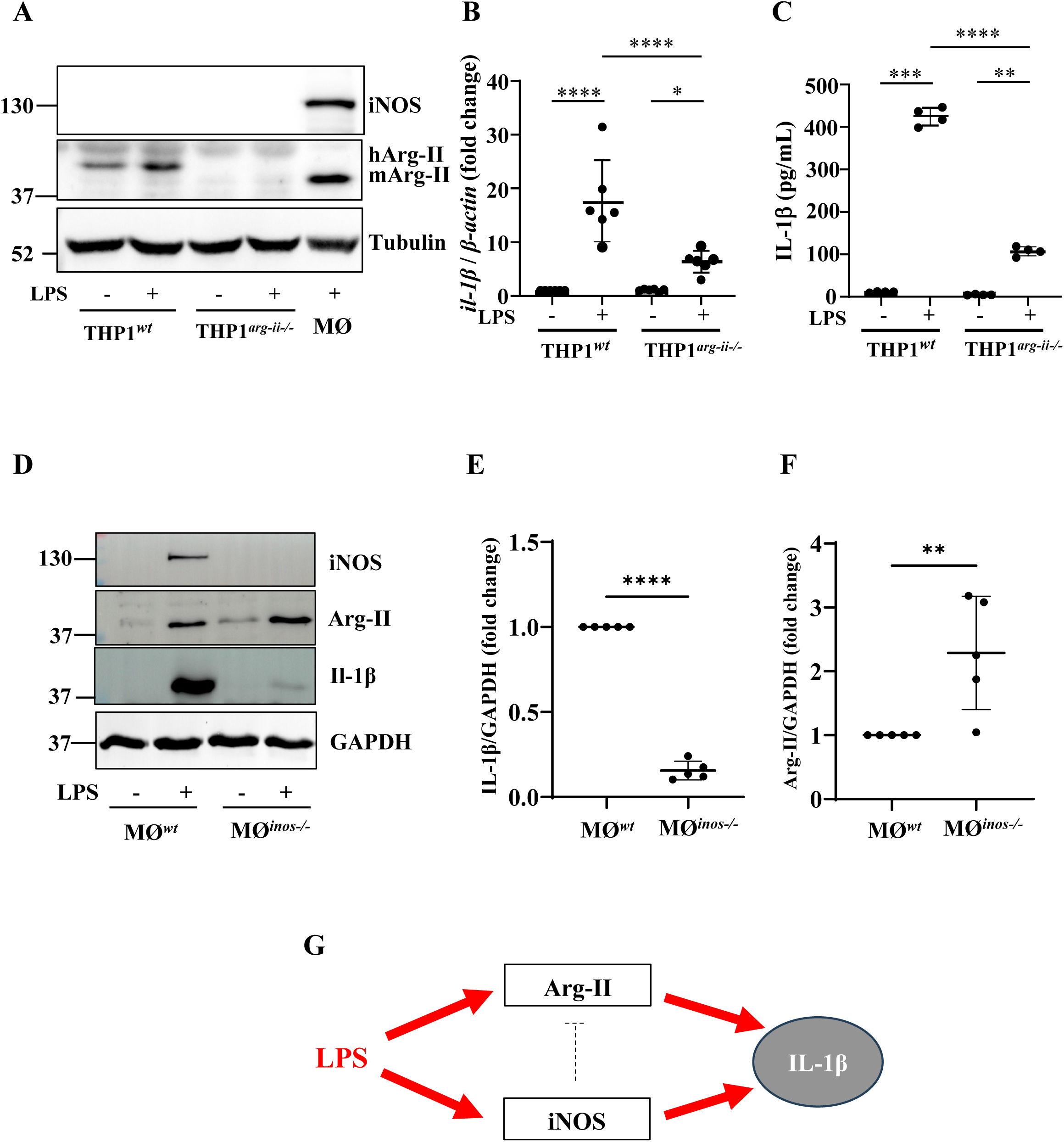
Effects of Arg-II and iNOS in regulation of IL-1β production in macrophages. (**A**) Immunoblotting analysis of Arg-II and iNOS in human THP1*^wt^* and THP1*^arg-ii-/-^* cells stimulated with LPS (100 ng/mL, 24 hours). Tubulin served as protein loading control. Mouse bone-marrow-derived macrophages (MØ) treated with LPS was used as positive control for iNOS and Arg-II detection; Molecular weight (kDa) is indicated at the side of the blots; (**B**) qRT-PCR analyzing mRNA levels of *il-1β* in the cells. *β-actin* served as internal reference (n=6); (**C**), IL-1β levels in the conditioned medium from the THP1*^wt^* and THP1*^arg-ii-/-^* cells stimulated with LPS (100 ng/mL, 24 hours) measured by ELISA; (**D**) Immunoblotting analysis showing Arg-II, iNOS, and IL-1β precursor protein levels in the bone-marrow-derived macrophages (MØ) stimulated with LPS (100 ng/mL, 24 hours). GAPDH served as protein loading controls. Molecular weight (kDa) is indicated at the side of the blots (n=5 independent experiments). The plot graphs show the quantification of IL-1β (**E**), and Arg-II (**F**) protein levels on the immunoblots. *p≤0.05, **p≤0.01, ***p≤0.005, ****p≤0.001 between the indicated groups. (**G**) Schematic illustration of IL-1β production regulated by iNOS and Arg-II in macrophages. The dotted line indicates inhibition. *wt*, wild-type; *arg-ii^−/−^, arg-ii* gene knockout; *inos^−/−^*, *inos* gene knockout; hArg-II, human Arg-II; mArg-II, mouse Arg-II.

**Suppl. Fig. 7.**
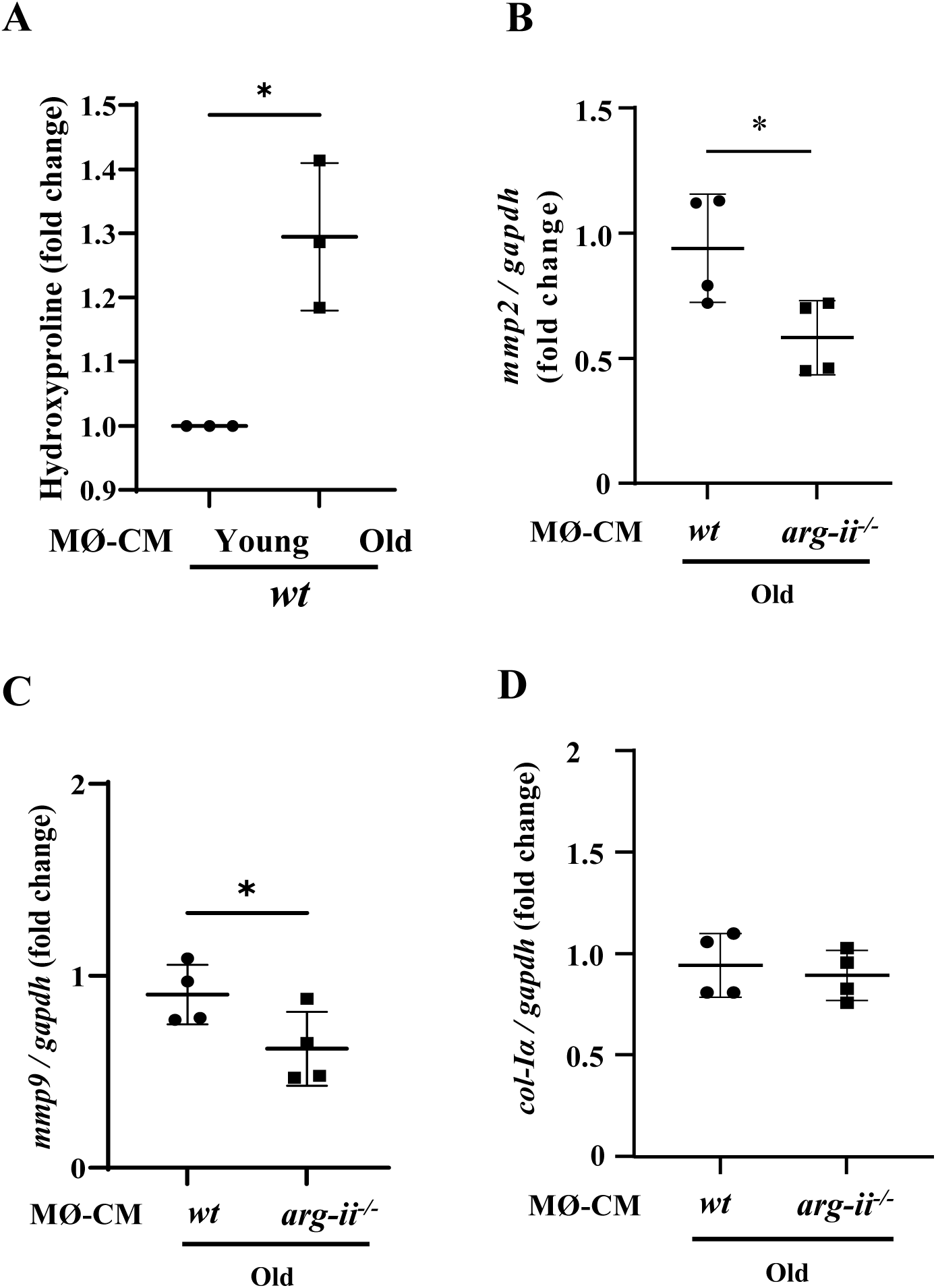
Crosstalk between splenic macrophages and cardiac fibroblasts. (**A**) Collagen production measured as hydroxyproline content in mouse *wt* fibroblasts treated with conditioned media (CM) from young and old *wt* splenic cells (96 hours of incubation) (n=3 mice in each group). mRNA expression levels of (**B**) *mmp2*, (**C**) *mmp9* and (**D**) *col-Iα* in *wt* fibroblasts treated with conditioned media (CM) from old *wt* and *arg-ii^−/−^* splenic cells (96 hours of incubation) were analyzed by qRT-PCR. *gapdh* served as the reference. (n=4 mice per group). Data are expressed as fold change to respective control group. \**p*≤0.05 between the indicated groups. MØ, splenic macrophage; CM, conditioned media; *wt*, wild-type mice; *arg-ii^−/−^*, *arg-ii* gene knockout mice.

**Suppl. Fig. 8.**
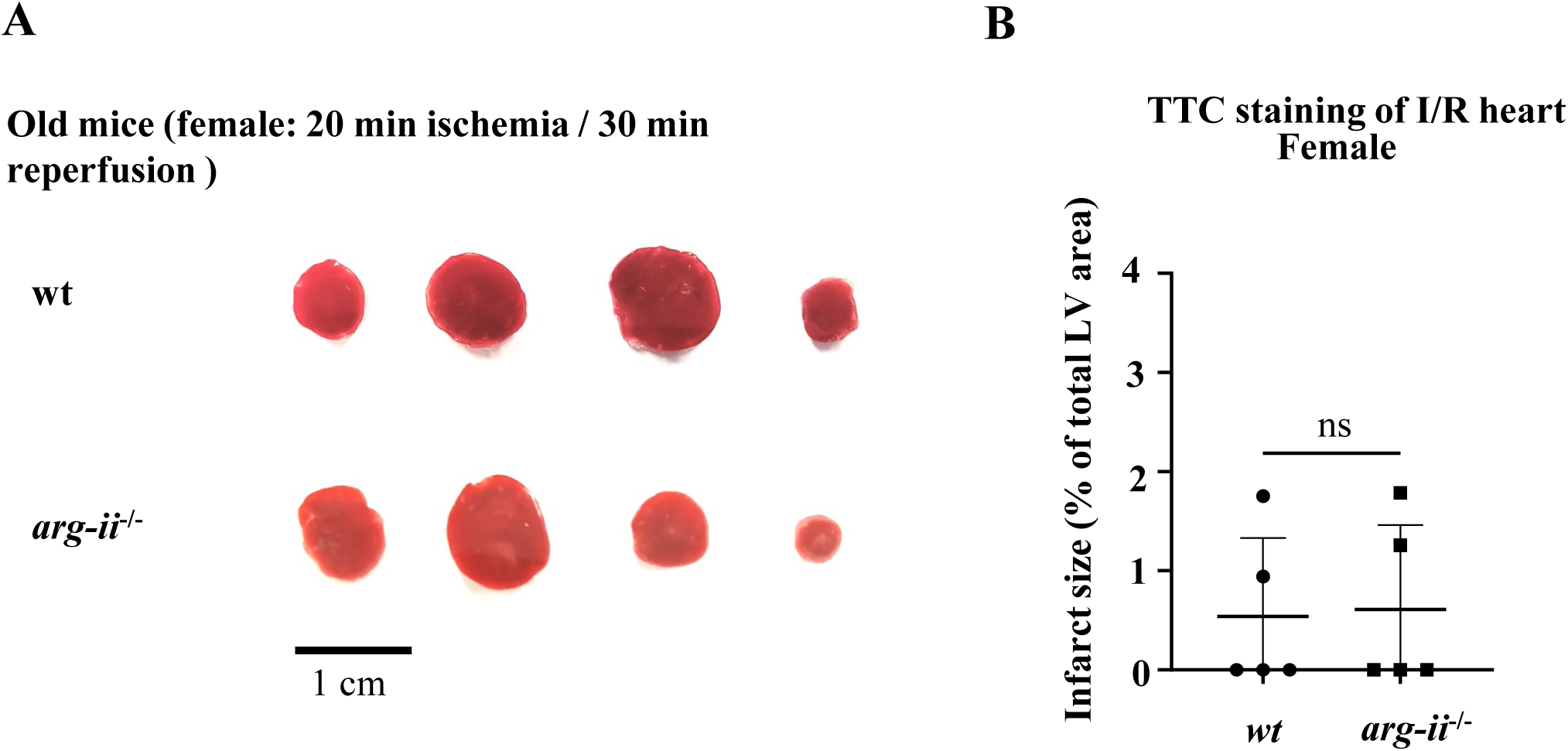
Short ischemia/reperfusion induces negligible myocardial infarct. (**A**) Representative sections of *wt* and *arg-ii^−/−^* hearts stained with 2,3,5-TTC and (**B**) relative quantification of infarcted areas. 20 minutes global ischemia is followed by 30 minutes reperfusion. *wt*, wild-type mice; *arg-ii^−/−^*, *arg-ii* gene knockout mice.

**Suppl. Fig. 9.**
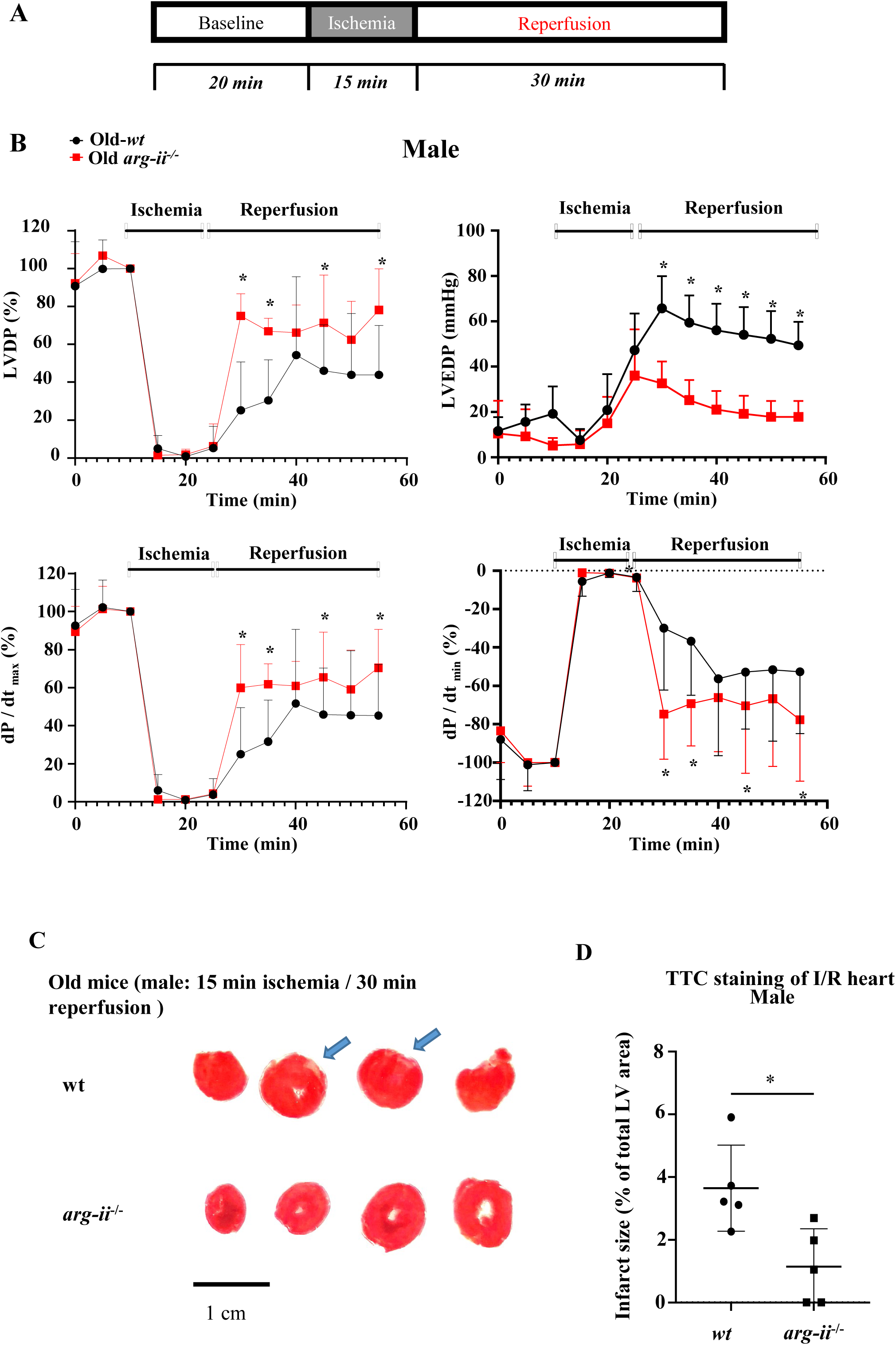
*Arg-ii* ablation improves heart function recovery from global ischemia/reperfusion injury (I/R-I) in male mice. (**A**) Protocol applied for ex vivo Langendorff-heart functional assessment of old *wt* and age-matched *arg-ii^−/−^* male mice. After baseline recordings, 15 minutes global ischemia is followed by 30 minutes of reperfusion; (**B**) Ex vivo Langendorff-heart assessment of old male *wt* (black line) and *arg-ii^−/−^* (red line) hearts. The graphs show the functional recovery of the left ventricular developed pressure (LVDP), left ventricular end diastolic pressure (LVED) and maximal rate of contraction (dP/dtmax) and relaxation (dP/dtmin). The data are expressed as % of recovery in respect to baseline values and represent the mean ± SD of data from 5 mice per group; (**C**) Representative sections of *wt* and *arg-ii^−/−^* male hearts stained with 2,3,5-TTC and (**D**) quantification of infarcted areas. *p≤0.05 between the indicated groups. *wt*, wild-type mice; *arg-ii^−/−^, arg-ii* gene knockout mice.

**Suppl. Fig. 10.**
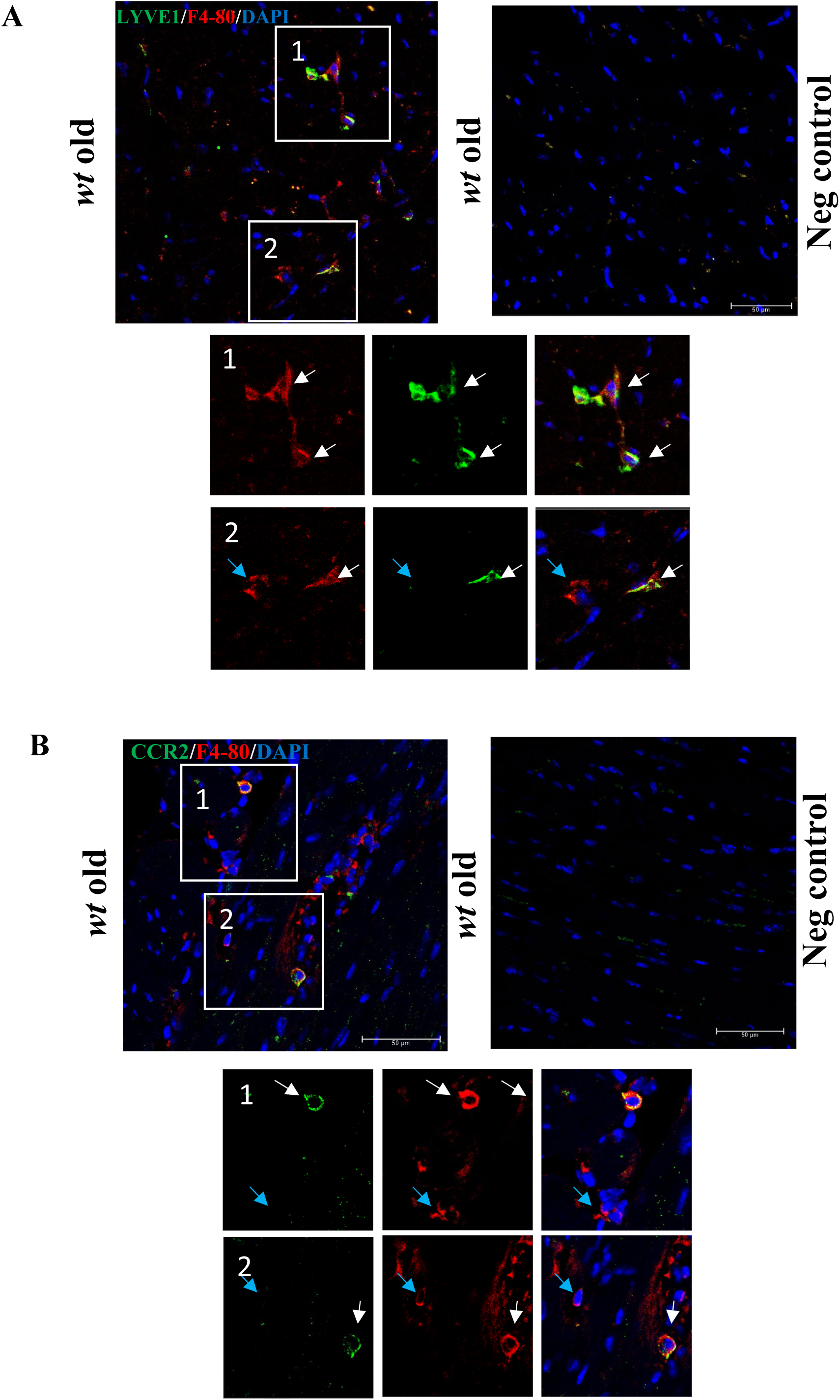
Validation of CCR2 and LYVE1 antibodies. (**A**) Confocal microscopy illustration of immunofluorescence double staining of Lyve1 (green) and F4-80 (red; macrophage marker). Scale bar = 50 µm; The negative control was generated by omitting the two primary antibodies. The enlarged region of interests (ROIs) 1 and 2 show the existence of both LYVE1^+^/F4-80^+^ and LYVE^−^/F4-80^+^ signal excluding the cross-reactivity between secondary antibodies. (**B**) Confocal microscopy illustration of immunofluorescence double staining of CCR2 (green) and F4-80 (red; macrophage marker). Scale bar = 50 µm; The negative control was again generated by omitting the two primary antibodies. ROIs 1 and 2 show the existence of both CCR2^+^/F4-80^+^ and CCR2^−^/F4-80^+^ signal excluding the cross-reactivity between secondary antibodies.

## Original blot

**Figure 4**

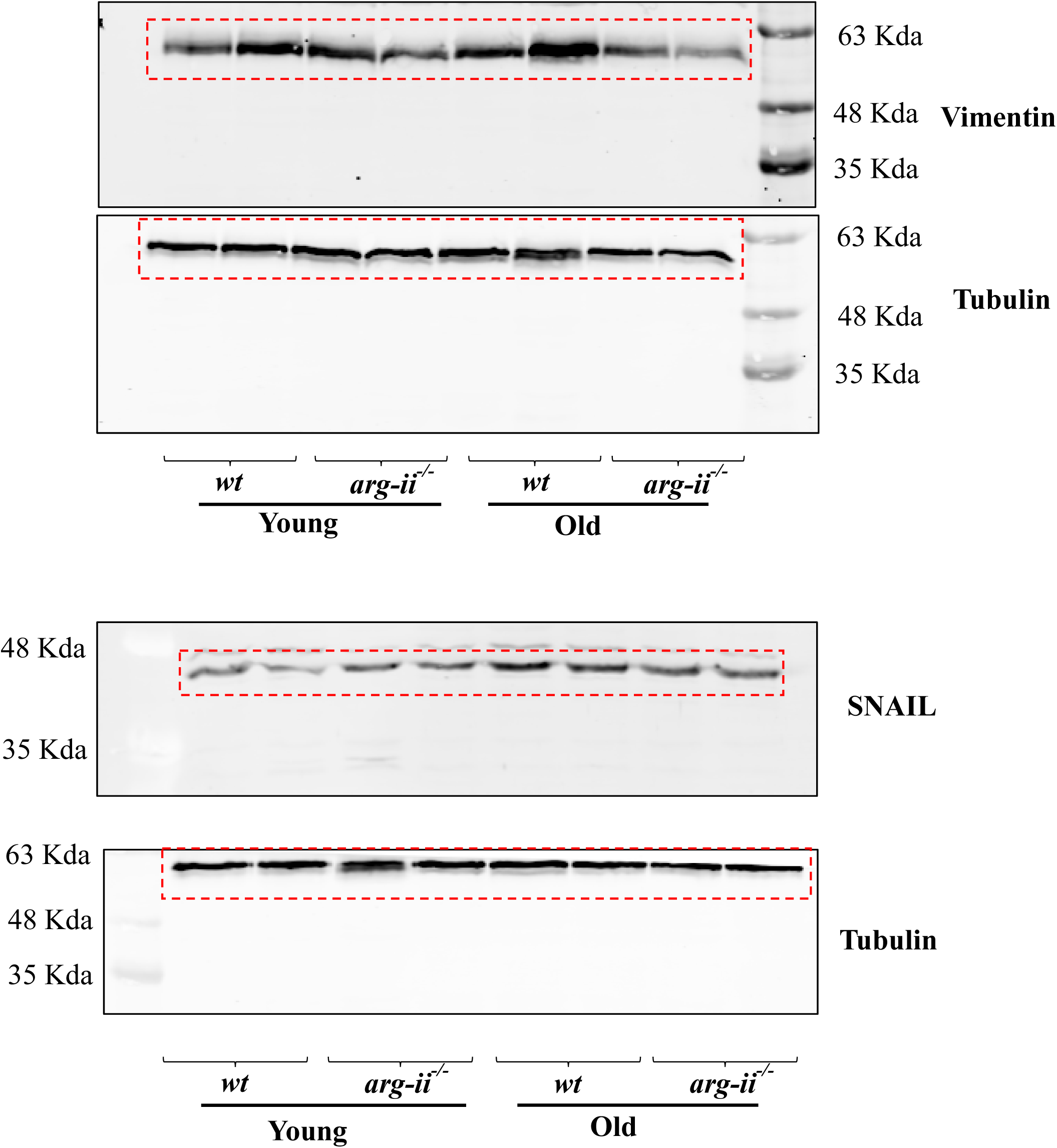
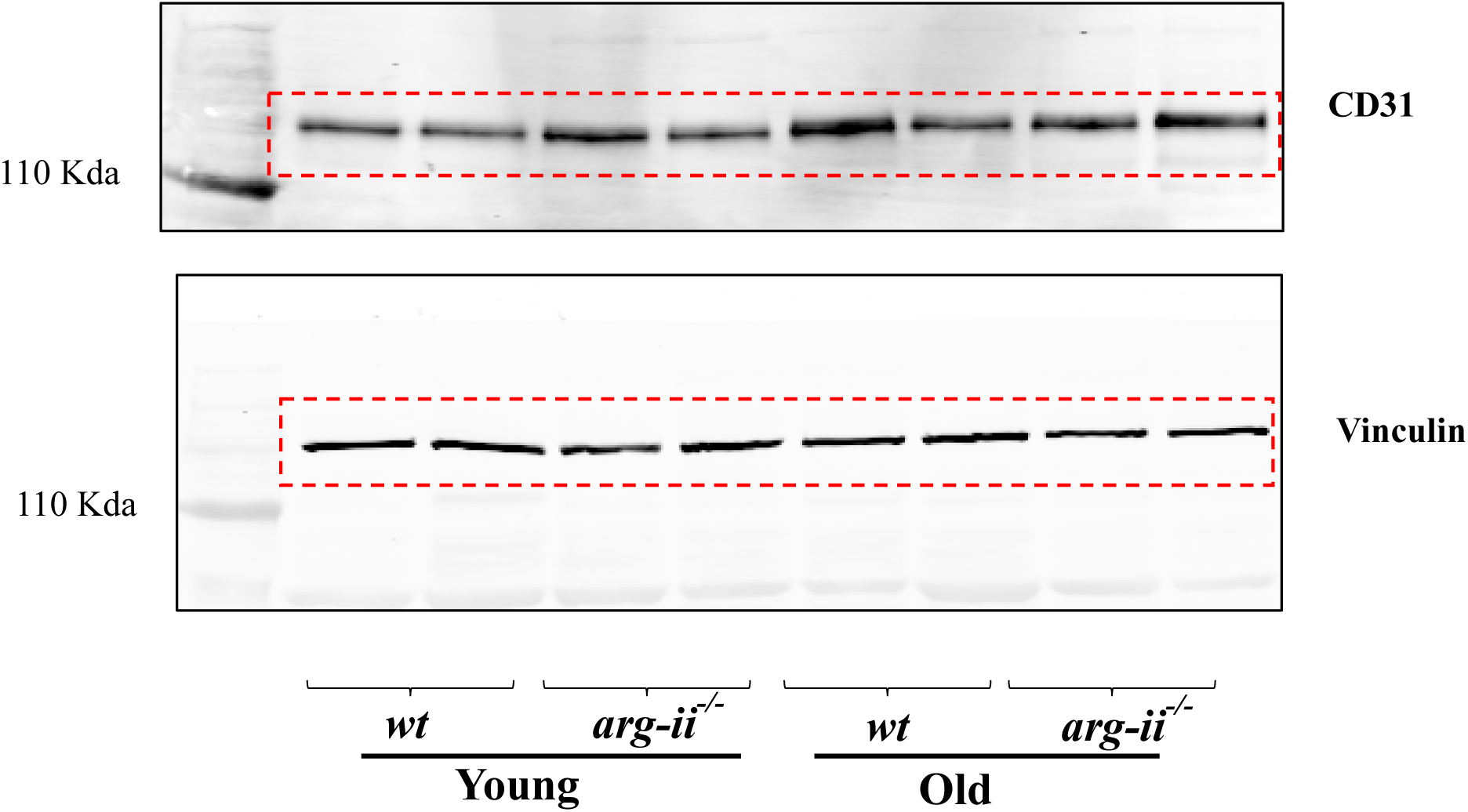

**Figure 5**

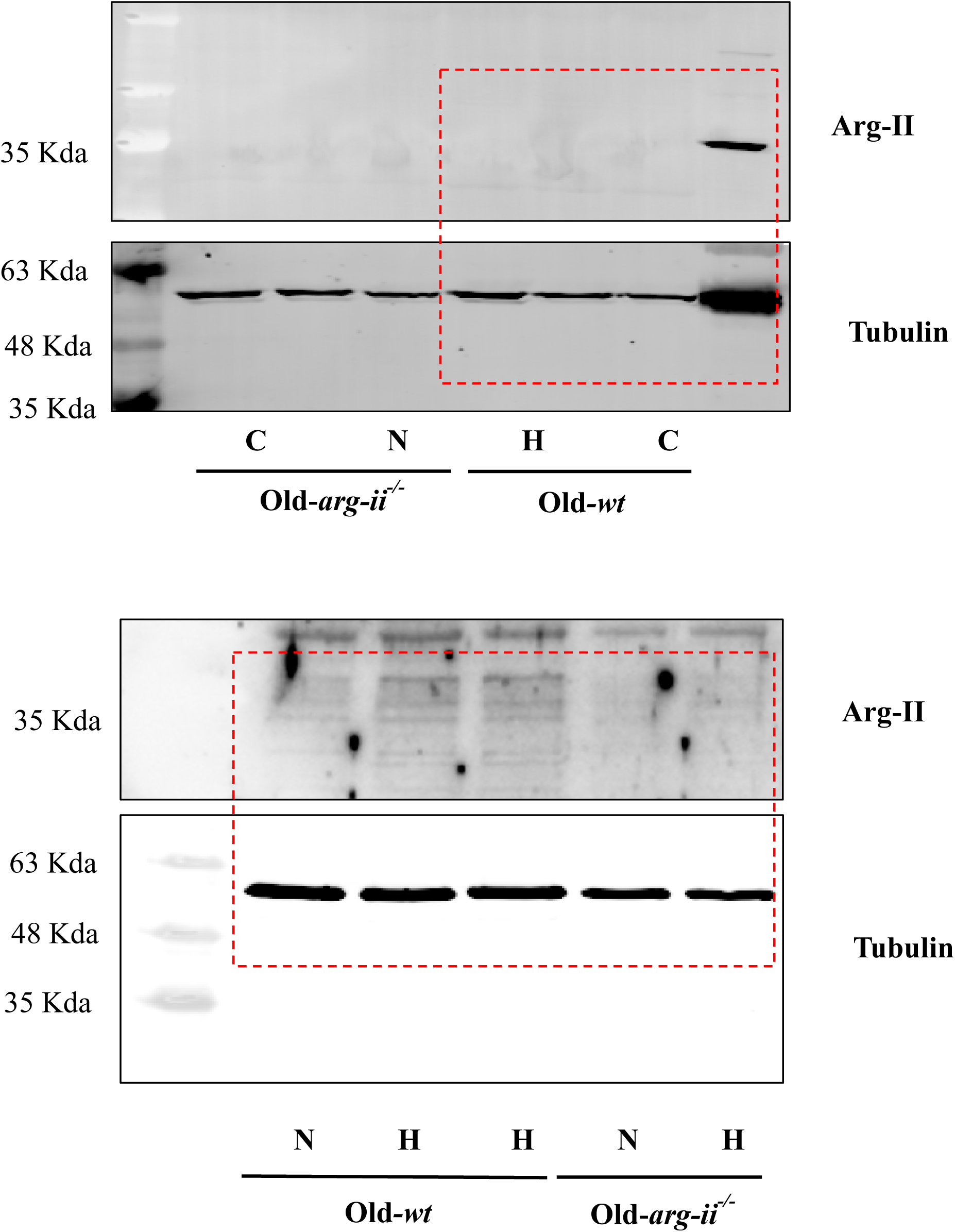

**Figure 6**

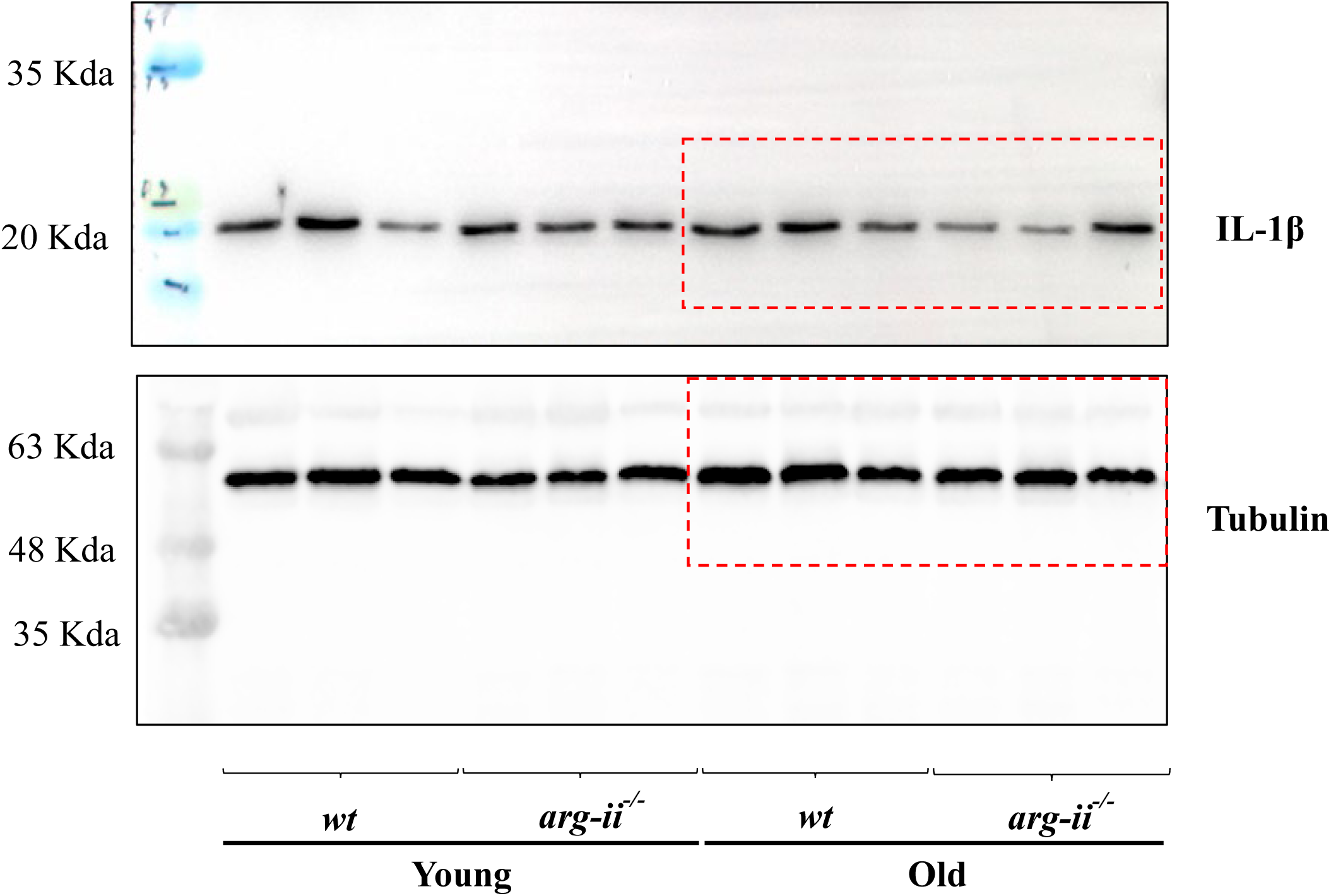

**Figure 7**

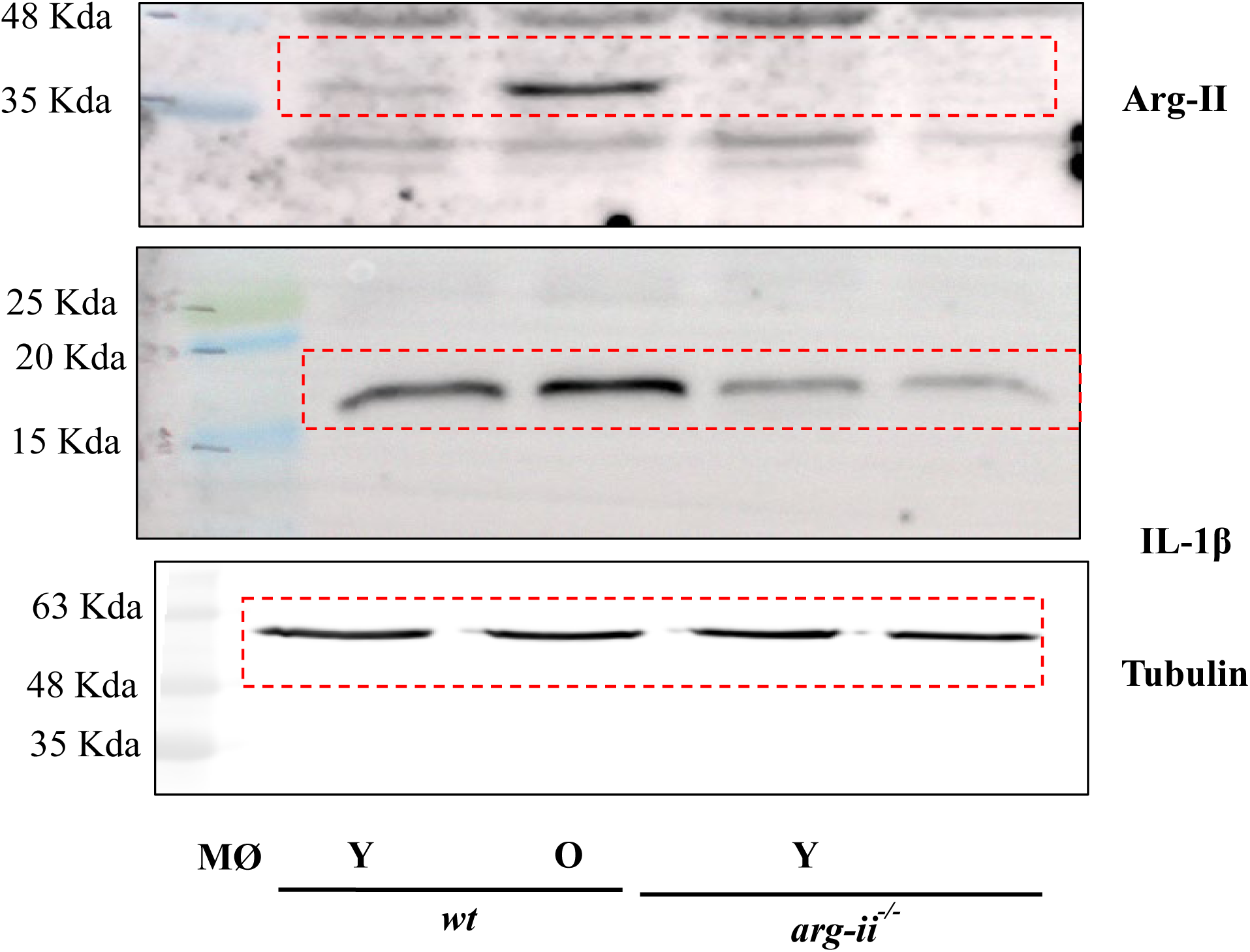

**Figure 8 F**

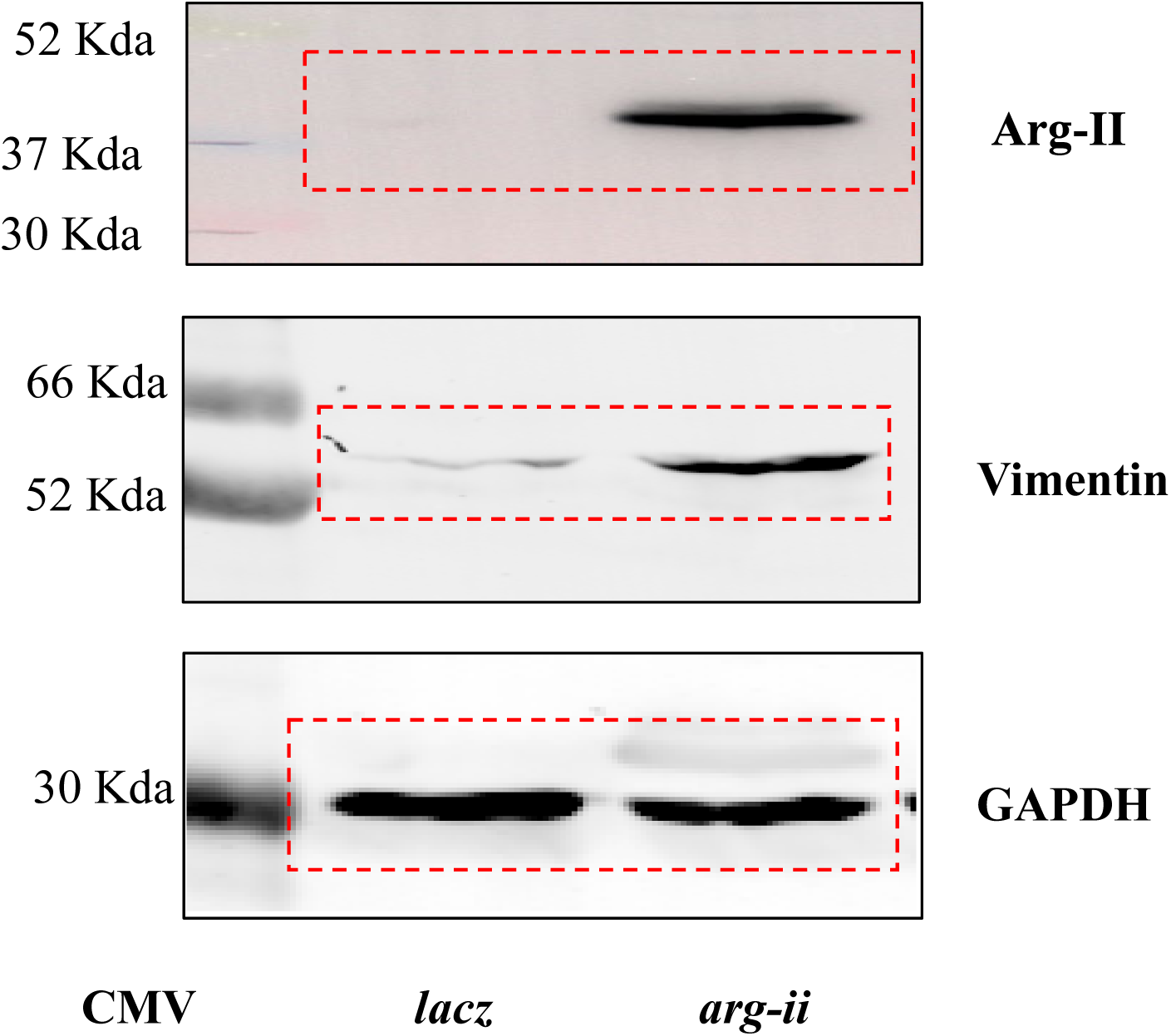

**Figure 8 M**

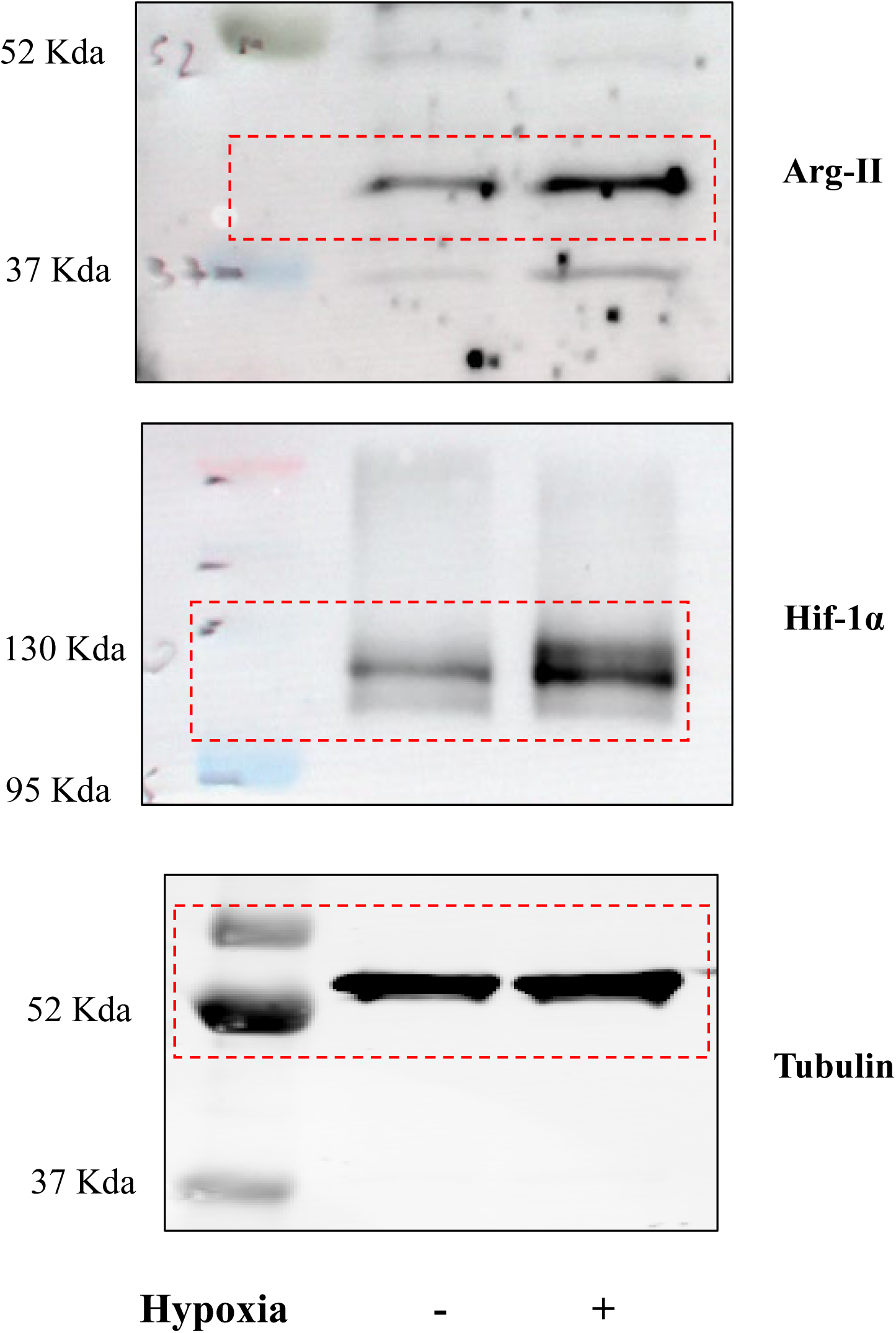

**Figure 9 A**

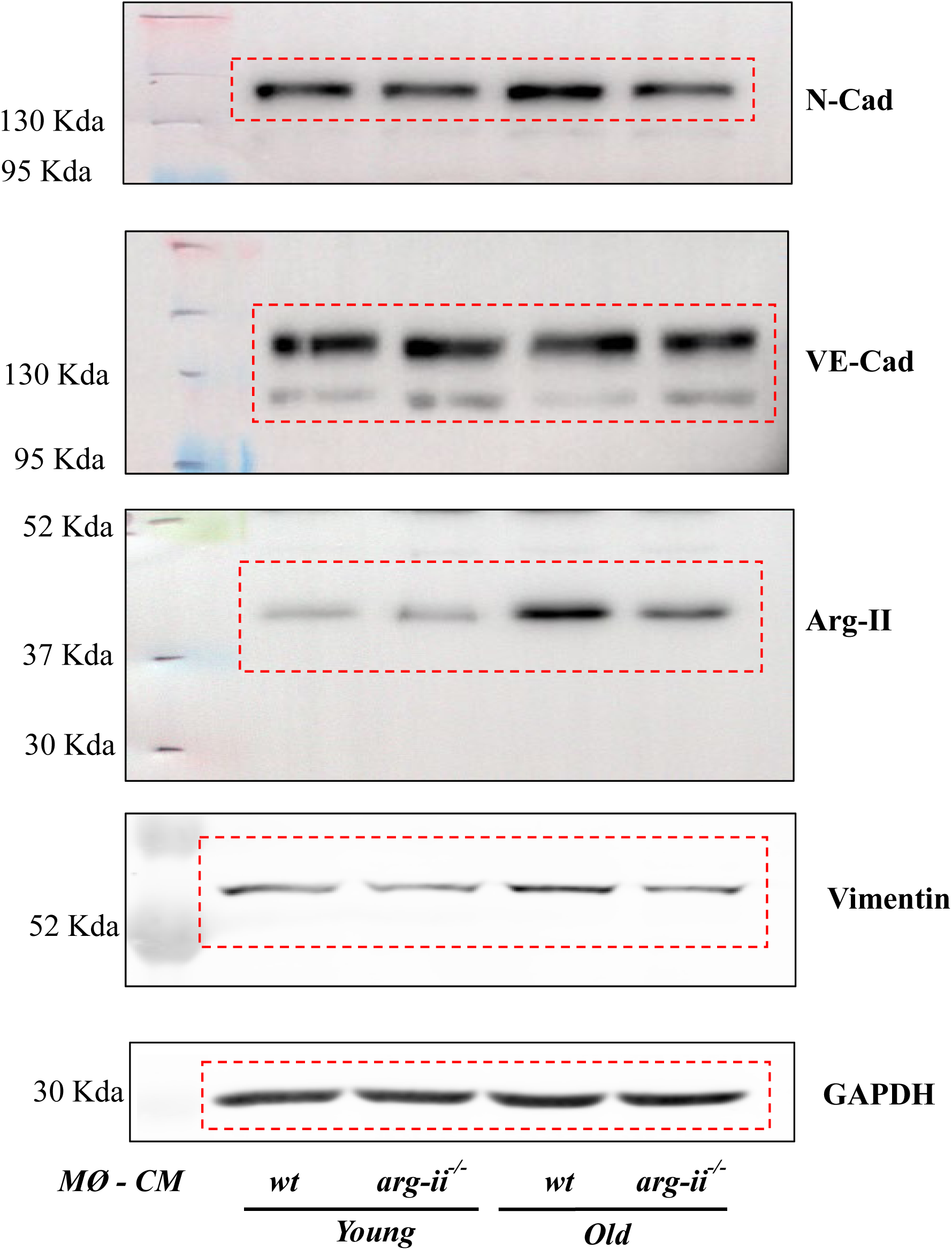

**Figure 9 F**

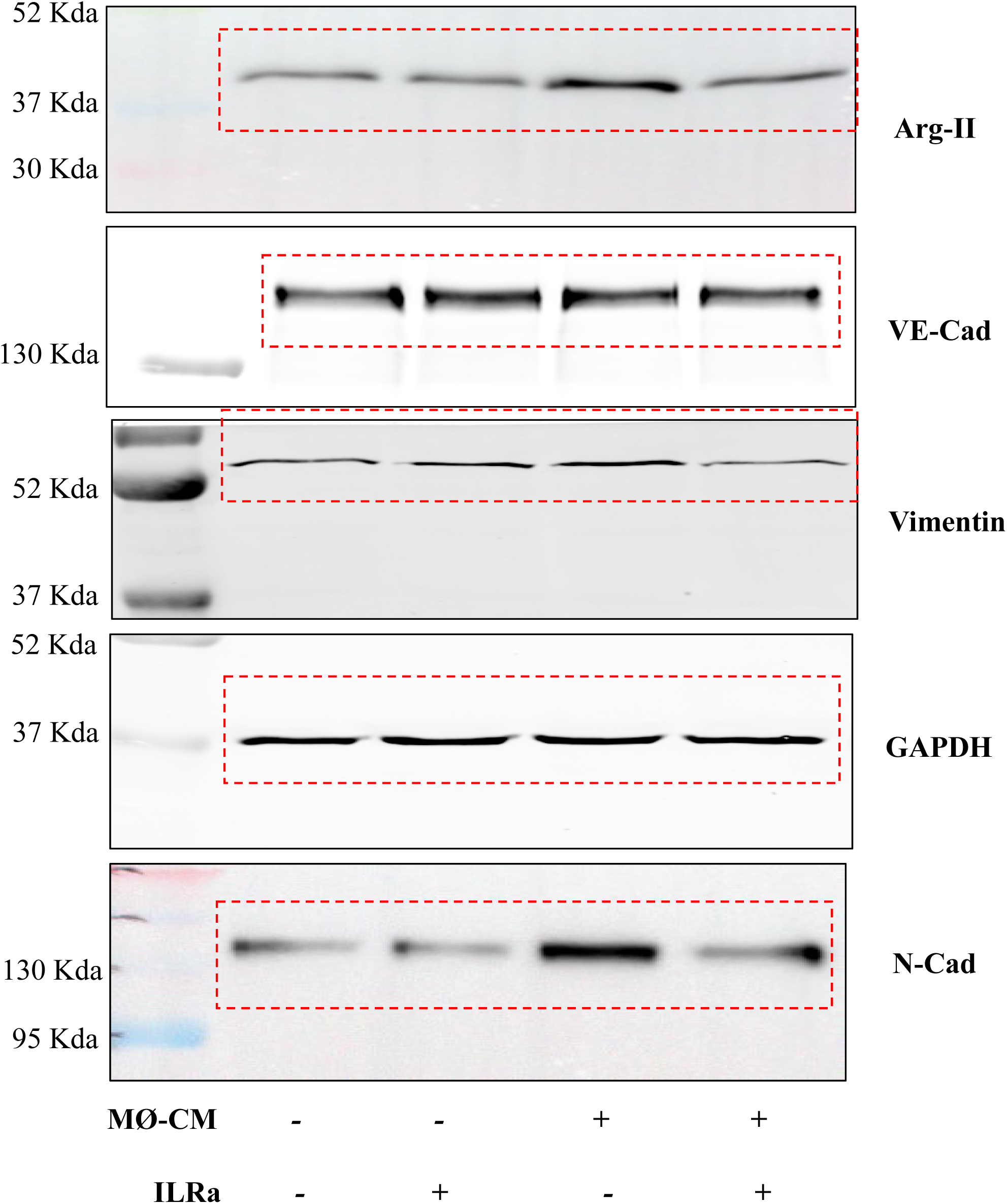

**Supplemental Figure 5**

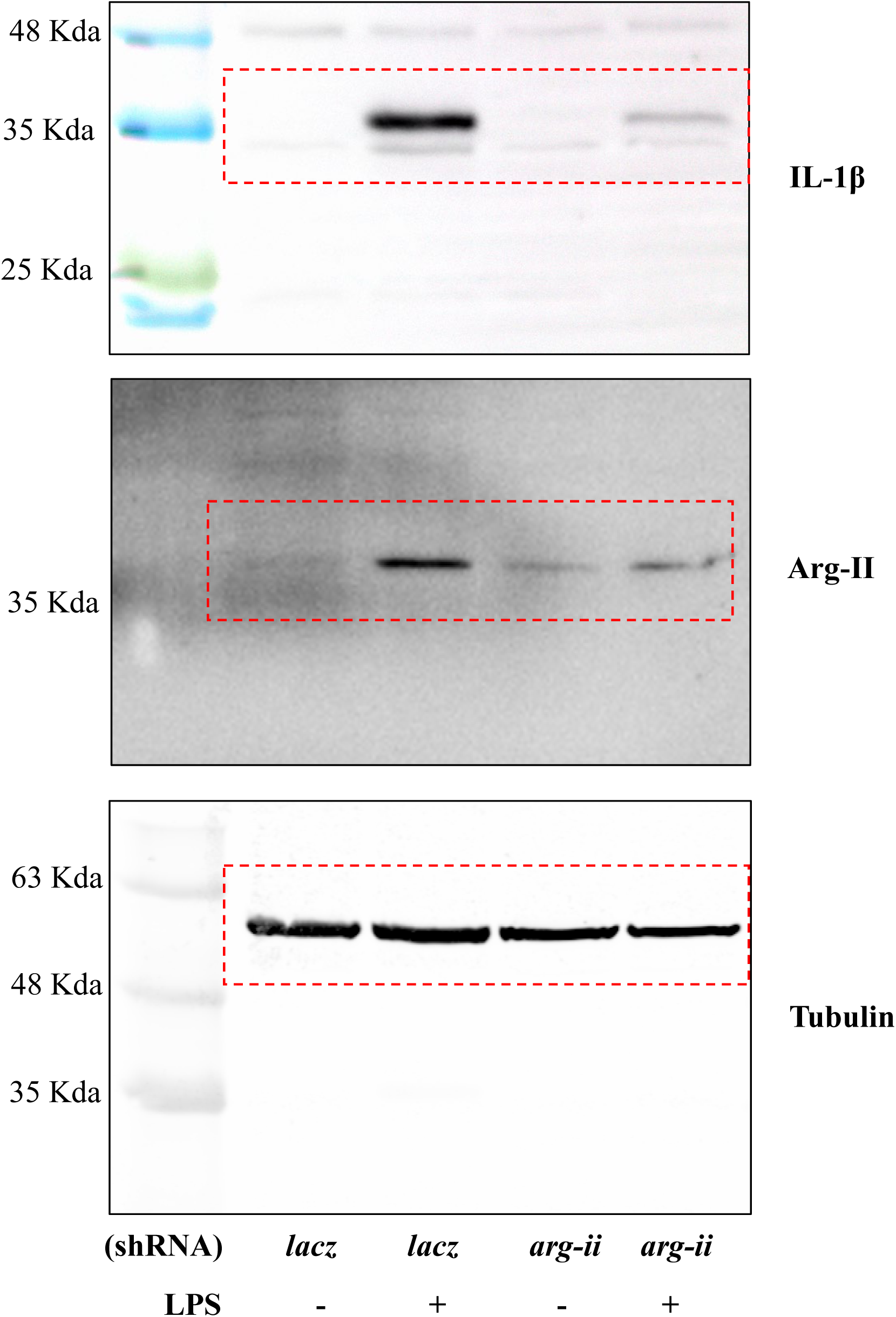

**Supplemental Figure 6 A**

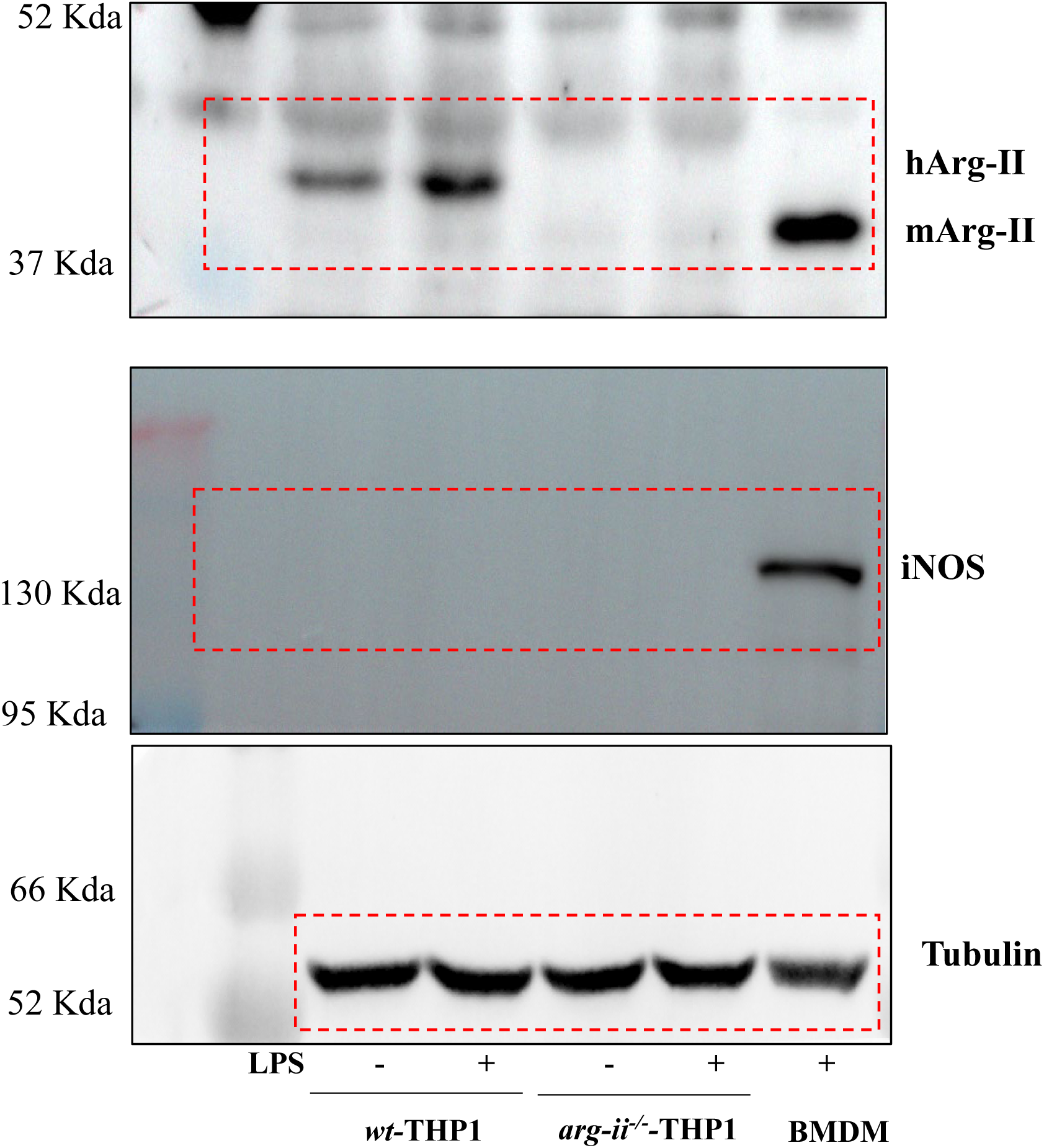

**Supplemental Figure 6 D**

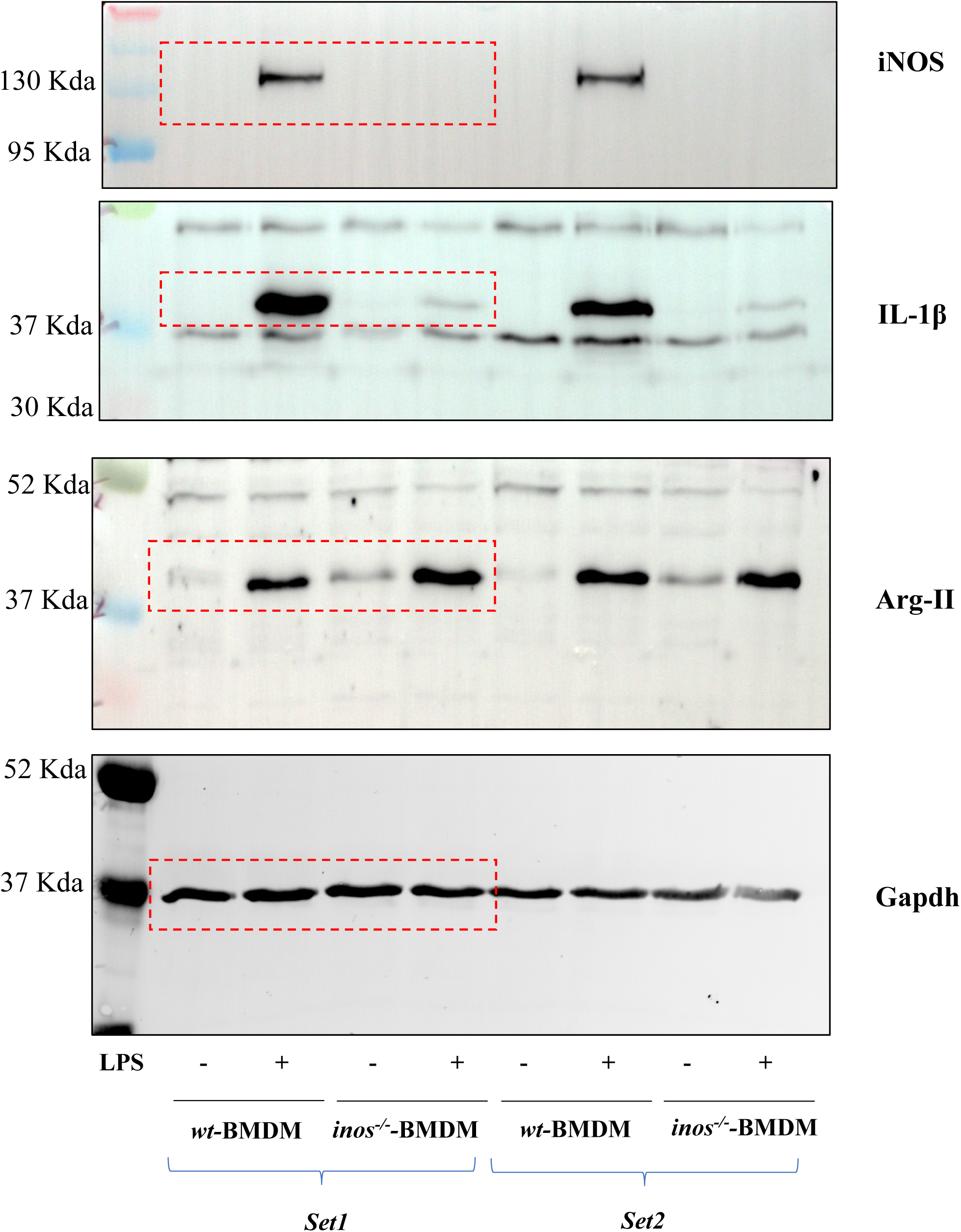

